# Differential chromatin accessibility landscape reveals the structural and functional features of the allopolyploid wheat chromosomes

**DOI:** 10.1101/2020.05.04.076737

**Authors:** Katherine W. Jordan, Fei He, Monica Fernandez de Soto, Alina Akhunova, Eduard Akhunov

## Abstract

**Background:** We have a limited understanding of how the complexity of the wheat genome influences the distribution of chromatin states along the homoeologous chromosomes. Using a differential nuclease sensitivity (DNS) assay, we investigated the chromatin states in the coding and transposon element (TE) -rich repetitive regions of the allopolyploid wheat genome.

**Results:** We observed a negative chromatin accessibility gradient along the telomere-centromere axis with mostly open and closed chromatin located in the distal and pericentromeric regions of chromosomes, respectively. This trend was mirrored by the TE-rich intergenic regions, but not by the genic regions, which showed similar averages of chromatin accessibility levels along the chromosomes. The genes’ proximity to TEs was negatively associated with chromatin accessibility. The chromatin states of TEs was dependent on their type, proximity to genes, and chromosomal position. Both the distance between genes and TE composition appear to play a more important role in the chromatin accessibility along the chromosomes than chromosomal position. The majority of MNase hypersensitive regions were located within the TEs. The DNS assay accurately predicted previously detected centromere locations. SNPs located within more accessible chromatin explain a higher proportion of genetic variance for a number of agronomic traits than SNPs located within closed chromatin.

**Conclusions:** The chromatin states in the wheat genome are shaped by the interplay of repetitive and gene-encoding regions that are predictive of the functional and structural organization of chromosomes, providing a powerful framework for detecting genomic features involved in gene regulation and prioritizing genomic variation to explain phenotypes.

## Background

The organization of chromatin affects cellular processes by controlling access to the genomic regions involved in the regulation of transcription, recombination, replication, and DNA repair [1, 2]. The elementary units of chromatin, nucleosomes, are mostly composed of histone octamers wrapped around by 147 bp of DNA connected by approximately 50 bp of linker DNA. Chromatin accessibility varies across the genome and is defined by the density of DNA-associated proteins, mostly nucleosome-forming histones, and the rate of association and dissociation of DNA-protein complexes. A broad range of nucleosome turnover rates and nucleosome occupancy levels was observed for different genomic regions, from high nucleosome occupancy and low turnover rate in heterochromatic regions to low nucleosome occupancy and high turnover rate in transcription start site regions [1].

In contrast to heterochromatic regions, nucleosomes in promoters and enhancers were shown to dynamically change between accessible and inaccessible configurations in response to developmental and environmental signals activating or suppressing gene expression [1,3,4]. These changes in chromatin states are associated with post-transcriptional histone modifications mediated by a large number of chromatin-associated proteins. For example, transition from open to closed chromatin, accompanying transcriptional suppression, could be promoted by Polycomb protein complexes [5, 6], which could also be involved in long-range interactions with distant *cis*-regulatory elements [7]. Therefore, open chromatin states reflect the regulatory potential of a genomic region and their characterization helps to accurately identify promoters, enhancers, and transcription factor binding sites.

While intergenic regions in large genomes are mostly composed of TEs, and possess largely inaccessible chromatin [8, 9], TEs along with distant *cis*-regulatory elements appear to play an active role in gene regulation and the structural organization of chromosomes. Long-range connections established between TEs and distant *cis*-regulatory elements, with their target genes were shown to contribute to gene regulation and shaping the 3D chromatin architecture [7,10–12]. In addition, interactions between the CENH3 histone-containing nucleosomes and TEs was demonstrated to be critical for the formation of active centromeres [13]. These studies provide evidence supporting the significance of TE-rich intergenic regions in defining both the structural and functional organization of chromatin in large genomes.

Combined with other methods of epigenomic profiling [14], chromatin accessibility assays helped to better understand the general principles underlying nucleosome organization across the genomes of major crops, including wheat, maize, rice, tomato, *Medicago truncatula,* and *Arabidopsis* [4,7,15–18]. An assay based on digestion with different concentrations of micrococcal nuclease (MNase) followed by the next-generation sequencing (DNS-seq) of digested genomic libraries was used to detect chromatin regions hyper-resistant or hyper-sensitive to MNase treatment [3]. DNS-seq of plant chromatin revealed “fragile nucleosomes” that showed MNase sensitive footprints (MSF) under light digest, but disappear under heavy digest conditions. These MSFs were significantly enriched in the genic and transcription factor binding regions, overlapped with the highly recombinogenic regions, and harbored genetic variants explaining most of the phenotypic variation in maize [3, 4]. Differences in nucleosome depleted regions between high and low expressed genes have also been reported in *Arabidopsis*, rice, and maize [4,15,16], linking an open chromatin state with higher gene expression. Consistent with the DNS-seq results in both plant and animal genomes, DNase I hypersensitive regions with open chromatin are often associated with proximal *cis*-regulatory elements [4,15– 19]. However, a substantial fraction of DNase I hypersensitive sites, some harboring distal *cis*-regulatory elements, were detected in intergenic regions [7,18,20,21]. Many of these intergenic accessible chromatin regions overlap with known TEs [21], suggesting their regulatory function, a possibility supported by the ability of maize TE-associated elements located within the accessible chromatin regions to drive reporter gene expression [22].

The hexaploid wheat genome (genome formula AABBDD) was formed by two recent hybridizations of three diploid progenitors [23–26], which diverged about 5.5 million years ago [27]. Previous studies demonstrated that the long-term post-hybridization adjustment of gene regulation as a consequence of increased gene dosage was accompanied by epigenetic, structural, and gene expression modifications [18,28–32]. Analysis of syntenic gene triplets in the allopolyploid genome showed that a gene expression bias towards one of the homoeologous copies was associated with changes in histone epigenetic marks, DNA methylation, and chromatin sensitivity to DNAse I and Transposase Tn5 treatments within the proximal *cis*-regulatory regions or gene body, thus connecting the chromatin and epigenetic states with imbalanced expression of duplicated genes [18,29,30]. While similar subgenome dominance in polyploid monkeyflower was accompanied by subgenome-specific epigenetic differences in the TEs near genes [33], no such dependence between DNA methylation within TEs and expression bias was obvious in wheat [30], even though a correlation between genome-specific promoter methylation and gene expression was observed [31]. The relationship between the epigenetic and chromatin states in the genic regions and TE regions near genes, and its impact on gene expression still remain unclear in understanding how mechanisms aimed at suppressing the transcriptional activity of transposable elements (TEs) while maintaining active gene expression exist with the proliferation of TEs in the wheat genome.

Extensive TE proliferation in the wheat genome [34] provides a unique opportunity to investigate the interplay between the repetitive and gene coding fractions of a large genome [35]. While the TE composition among three wheat genomes is similar, all chromosomes show a strong gradient in TE content along the centromere-telomere axis, where the distal ends have 73-89% less TE content than regions close to centromeres [34]. The intergenomic analysis of syntenic regions showed that while there is relatively low sequence similarity between the genomes in the intergenic regions, overall gene order and distance between the genes are conserved in all three wheat genomes. The conservation of this genome structure, in spite of complete replacement of TE content in the wheat genomes since their divergence from the common ancestor [27], suggests that intergenic distance rather than the intergenic sequence itself is under evolutionary pressure [34]. These results indicate that the gradient in TE abundance along the chromosomes is associated with an increase in intergenic distance from the telomere to the centromere is likely a product of selection. Here we used digestion with different concentrations of MNase to probe the genome-wide chromatin accessibility in the allopolyploid wheat genome. By investigating how chromatin accessibility changes in relation to gene density, gene expression levels, intergenic distance, TE content and composition, and chromosomal position we sought to better understand the impact of the repetitive fraction of the wheat genome on the functional and structural organization of the wheat chromosomes.

## Results

### Genomic and chromosomal patterns of differential nuclease sensitivity

Previous studies demonstrated that the D genome has an overall lower level of repressive histone marks and DNA methylation than the A and B genomes [18, 30]. These trends correlate with a slightly higher proportion of genes showing D-genome biased expression [30]. To investigate whether these patterns of epigenomic and gene expression variation are also reflected in the level of chromatin accessibility, we assessed wheat chromatin states using digestion with different concentrations of micrococcal nuclease (MNase). Genomic libraries prepared using light and heavy MNase digests were sequenced producing nearly 1.75 billion paired end (PE) reads (Table S1 in Additional File 1), of which about 1.2 billion PE reads uniquely mapped to the wheat genome [35]. This dataset was used to calculate the differential nuclease sensitivity (DNS) scores for 10-bp intervals across the genome. The high level of correlation between the two biological replicates (*r* = 0.98, *p* < 2.2 x10^-16^) (Fig. S1 in Additional File 1) is suggestive of good consistency between the experiments. Segmentation of the wheat genome based on the distribution of DNS scores was performed using the iSeg program [36], which identifies outlier regions (>1.5 standard deviations of the genome wide DNS score) corresponding to either MNase hyper-sensitive footprints (MSF) or MNase hyper-resistant footprints (MRF). A total of 177 Mb (1.26%) of the genome were classified as MSF, and 215 Mb (1.53%) of the genome were classified as MRF (Table 1, Table S2 in Additional File 1) [3].

**Table 1.** Distribution of DNS Scores across five chromosomal segments

| Comparison |  | Whole<br>Genome | A genome | B genome | D genome |
| --- | --- | --- | --- | --- | --- |
| Whole genome 2Mb windows† |  | 0.0031 | 0.0032 | -0.0016 | 0.009 |
| Segments‡ | R1 | 0.0457 | 0.0413 | 0.0398 | 0.0559 |
|  | R2a | -0.00669 | -0.0079 | -0.0146 | 0.00237 |
|  | C | -0.0131 | -0.0153 | -0.0135 | -0.0107 |
|  | R2b | -0.00949 | -0.0087 | -0.0154 | -0.00441 |
|  | R3 | 0.0431 | 0.0474 | 0.0368 | 0.0451 |
| Segmentation§ | MSF (Mb) | 177.3 | 59.4 | 67.2 | 50.7 |
|  | MRF (Mb) | 214.8 | 73.3 | 86.3 | 55.2 |
| Proximity to gene¶ | 2 kb upstream | 0.18 | 0.18 | 0.175 | 0.184 |
|  | 500 bp upstream | 0.261 | 0.256 | 0.259 | 0.268 |
|  | Gene Body | 0.0962 | 0.0926 | 0.096 | 0.10 |
|  | 2 kb downstream | 0.162 | 0.161 | 0.158 | 0.167 |
|  | Intergenic | 0.0068 | 0.0072 | 0.0050 | 0.0083 |

† Mean DNS score for 2 Mb windows
‡ Mean DNS score for entire genomic segment
§ Size in Mb of the regions considered significantly accessible or inaccessible by iSeg
¶ Mean DNS score for specific region

The genome-level DNS scores averaged across 2 Mb genomic windows were significantly higher in the D genome than both the A and B genomes (DNS_D_ = 0.009, *X*^2^= 339, *p* < 2.2 x 10^-16^, Kruskal Wallis test) (Fig. 1a, Table 1, Table S2 in Additional File 1), while the A genome DNS scores were significantly higher than the B genome (DNS_A_ = 0.0032, DNS_B_ = −0.0016, *X*^2^= 67.5, *p* < 2.2 x 10 ^-16^, Kruskal Wallis test). The genome-level DNS patterns are also supported at the chromosome level, where the D genome chromosomes have predominantly accessible chromatin (Table S2 in Additional File 1), with the chromatin of chromosomes 5D and 4B being most (DNS = 0.015) and least (DNS = −0.012) accessible, respectively.

**Figure 1.**
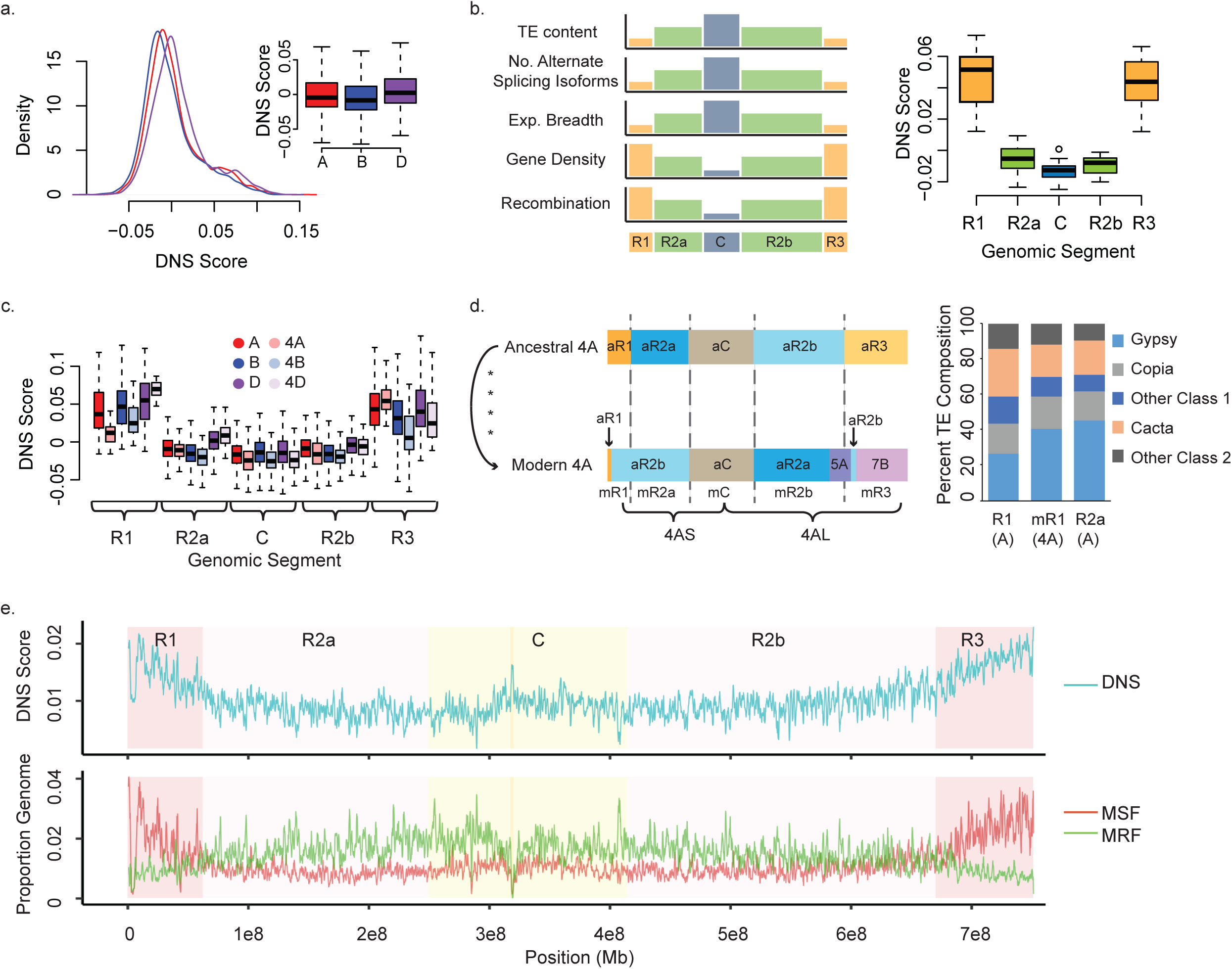
Distribution of DNS scores across the genomic regions and chromosomes. **(a)** Density of DNS scores across the whole genome in 2Mb windows for the A (red), B (blue), and D (purple) genomes. **Insert**, distribution of 2Mb window DNS scores by genome. **(b)** Distribution of DNS scores across genomic segments, distal R1, interstitial R2a, pericentromeric C, interstitial R2b, and distal R3. The relative distribution of genomic features used for the segmentation of wheat chromosomes is shown on the left. **(c)** Distribution of DNS scores by genomic segment and genome (A-red, B-blue, D-purple), homeologous chromosome group 4 DNS values are shown as lighter colors (chr4A-light pink, 4B-powder blue, 4D-light lavender). Distal segment R1 is reduced 3.6x compared to A genome R1 segments. **(d)** Structural evolution of wheat chromosome 4A and segmentation of the ancestral (aR1, aR2a, aC, aR2b, aR3) and modern (mR1, mR2a, mC, mR2b, mR3) 4A chromosomes. TE composition of the R1 segment on modern 4A (mR1), and R1 and R2a segments of remaining chromosomes from the A genome. **(e)** Top panel: Representative DNS scores for 1Mb windows across entire chromosome 3A. Genomic segments are shown in the background as dark pink for distal segments, light pink for interstitial segments, and pale yellow for the pericentromeric region, location of the centromere is dark yellow. Bottom panel: Proportion of 1Mb windows that are considered outliers for hyper-resistant regions MRF (green) and hyper-sensitive regions MSF (red) across chromosome 3A.

Previously, based on the distinct patterns of recombination rate, gene density, and expression breadth distribution, each of the 21 wheat chromosomes was partitioned into five regions, referred to as R1 and R3 for the distal ends of the short and long chromosomal arms, respectively, R2a and R2b for the interstitial regions on the short and long arms, respectively, and the C region, which represents the pericentromeric region [35, 37] (Fig. 1b). While R1 and R3 regions have high gene density and recombination rate, and reduced TE density, alternative splicing and gene expression breadth, the R2a, C, and R2b regions showed opposite trends [34,35,37]. Overall, the pericentromeric and interstitial chromosomal regions each have negative DNS scores (Table 1, Fig. 1b), while both distal regions have positive DNS scores of 0.046 and 0.053, mirroring the gradient for recombination rate (Fig. S2 in Additional File 1), gene density, and gene expression, and a directly opposite gradient for TE composition [35, 37].

The general trends of chromatin accessibility among the chromosomal segments are consistent among genomes, except that the interstitial regions of the D genome are more accessible than the corresponding regions in the A and B genomes (*X^2^*= 23.7, *p-value* = 7.1 x 10^-6^, Kruskal-Wallis test; W_AD_= 28, *p-value* = 0.0008 and W_BD_ =7, *p-value* = 2.2 x 10^-6^, Mann-Whitney-Wilcoxon test) (Fig. 1c). Overall, we observed a decline in chromatin accessibility along the telomere-centromere axis based on 2 Mb-window DNS scores spanning all chromosomes of all three wheat genomes (Fig. 1c, Figs. S3-9, Table S3). The DNS score differences between the pericentromeric and distal regions in the A and B genomes were higher than that in the D genome. For example, the DNS scores of the distal ends of the A genome were 13 and 15 times higher than the genome-wide mean (DNS_A_ = 3.2 x10^-3^), while the centromeric regions showed a 5-fold lower chromatin accessibility compared to the genome-wide mean (Table S3 in Additional File 1). Whereas in the D genome, whose mean DNS score was the highest (DNA_D_ = 9 x10^-3^), there was only five- and six-fold chromatin accessibility increase in the distal ends, and a 1.3-fold decrease in the centromeric regions. One of the likely factors affecting these differences in chromatin state between distal and pericentromeric regions is the relative position of the genomic region with respect to centromere.

To investigate this possibility, we compared the distribution of DNS scores along chromosome 4A relative to the other wheat chromosomes. Compared to its homoeologous chromosomes 4B and 4D, chromosome 4A has undergone two reciprocal translocations, and a peri- and para-centromeric inversion that has disrupted the ancestral structure of this chromosome [38, 39] (Fig. 1d). In the modern chromosome 4A, chromosomal arm 4AS is represented only by a small proportion of the ancestral 4AS arm with the majority of chromosomal segment R1 composed of the interstitial region of ancestral 4AL (Fig. 1d). The R2a segment of the modern-day 4AS arm is still represented by an interstitial chromosomal segment, but from ancestral 4AL. Present-day 4AL includes the ancestral interstitial segment of 4AS, which now makes up the interstitial region R2b of 4AL, with translocated portions of 5AL, and 7BS making up the majority of the R3 distal segment (Fig. 1d). These structural rearrangements result in the R1 distal region of 4AS now composed of the interstitial region of ancestral 4AL, and we hypothesize, based on the chromatin accessibility trends along the centromere-telomere axis of other chromosomes, that this R1 region should display a reduced DNS score similar to interstitial regions. Indeed, we observed nearly a 72% reduction in DNS score in the R1 region on chromosome 4A compared to the mean of other A genome R1 distal segments (Fig. 1c, Table S4 in Additional File 1). Moreover, in spite of homoeologous group 4 displaying the least accessible chromatin among other chromosomal groups, and chromosome 4B’s chromatin being the least accessible within the homoeologous group, the mean DNS score of chromosome 4B’s R1 segment was nearly 2.5 times higher than the mean of the chromosome 4A’s R1 segment (Fig. 1c). Considering that the B genome’s chromatin is on average more inaccessible than that in the A genome, these results indicate the R1 segment on chromosome 4A experienced a substantial reduction in chromatin accessibility. Comparison of the sequence composition between the 4A-R1 segment, and the R1 and R2a segments from other A genome chromosomes showed that the proportion of sequences represented by different classes of TEs in the 4A-R1 segment is more similar to that of the R2a interstitial segments rather than to that of the R1 segments (Fig. 1d). These results suggest that sequence composition likely plays a more important role in defining the chromatin accessibility differences between telomeric and pericentromeric chromosomal regions than the relative position on the chromosome.

### Hyper-sensitive and hyper-resistant regions of the wheat genome

Compared to the rest of the genome, we observed a significant enrichment of the MSFs in the distal R1 and R3 regions combined (Fisher’s exact test, *p-value* = 2 x 10^-4^) (Fig. 1b, Figs S3-9, Table S5 in Additional File 1). This trend was accompanied by corresponding enrichment of the MRFs in the pericentromeric and interstitial chromosomal regions including R2a, C, and R2b (Fisher’s exact test, *p-value* = 5 x 10^-3^), consistent with the observed overall trend in chromatin accessibility along the centromere-telomere axis (Fig. 1e). The 177 Mb of MSFs and 215 Mb of MRFs correspond to 2,156,684 and 2,605,884 unique genomic segments, respectively. Only 17% of MSFs and 1.8% of MRFs were located within the genic regions including annotated high-confidence (HC) gene models [35], and the 2 kb regions upstream and downstream of the coding sequences. This difference in the genomic distribution between MSF and MRF represents a significant enrichment for MSF near genes compared to that of MRF (Fig. 2a, Table S5, and Fig. S10 in Additional File 1; *p*-value = 2.2 x 10^-16^; Fisher’s Exact test (FET)). For both MSF and MRF around genes, nearly half are found within the gene body (8.1% of MSFs and 0.7% of MRF), while the other half are nearly equally distributed between the regions upstream and downstream of genes (Fig. 2a, Table S5 in Additional File 1). We detect 86% of annotated genes (90,941 genes) are located within 2 kb of at least one MSF, with an average of 4 MSF per gene, while only a total of 29,230 genes were located within at least 2 kb of MRF, with an average of 1.6 MRFs per gene. Similar proportions of MRF and MSF near genes were detected within each genome, with the exception that a higher percentage (20%) of MSF are found around genes in the D genome compared to that in the A (17%) and B (15%) genomes (Fig. S10 in Additional File 1; *p-values* < 2.2 x 10^-16^; FET). It is of note with respect to the distance from genes for MSF and MRF regions, the distance distribution reflects that MSF regions are located closer to genes than MRF (mean _MSF_ = 137 kb, mode _MSF_ = 10 kb, mean _MRF_ =175 kb, mode _MRF_ = 15 kb; Fig. 2b and Fig. S10 in Additional File 1), however both distributions are heavily skewed with averages >100 kb from annotated genes. This observation suggests that a considerable proportion of gene regulatory machinery is located in the intergenic regions of the genome, mostly composed of TEs, consistent with the recent findings in other large, complex plant genomes [7, 12].

**Figure 2.**
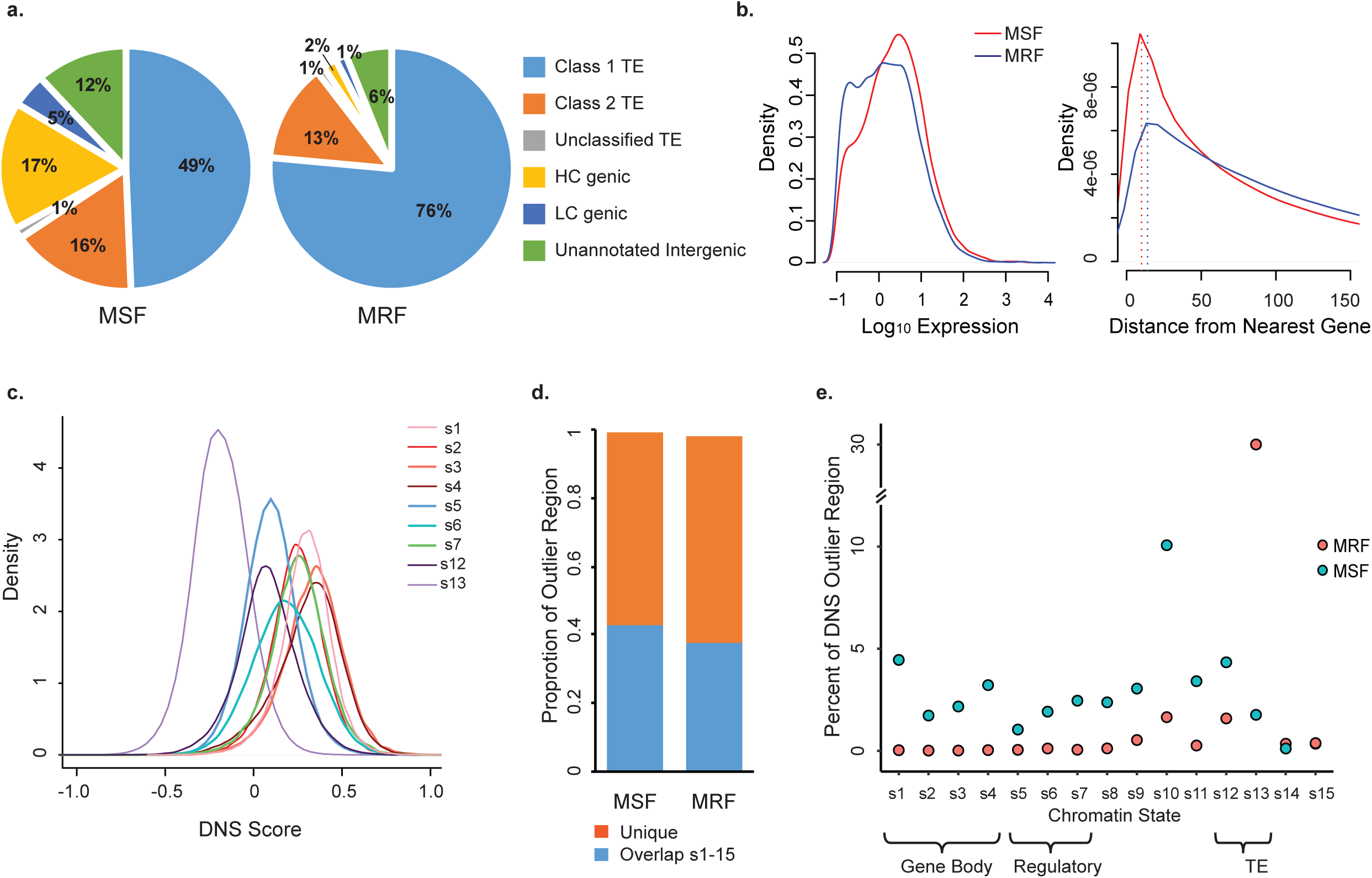
MNase hypersensitive (MSF) and hyperresistant (MRF) footprints in the wheat genome. **(a)** Distribution of MSF and MRF across the wheat genome. **(b)** Distribution of gene expression values for genes located in close proximity to MSF/MRF (left); distribution of distances (kb) between genes and MSF/MRF (right). Density peaks are marked by dashed line for both MSF (red) at 10 kb and MRF (blue) at 15 kb. **(c)** Distribution of genome-wide DNS score values calculated for nine out fifteen chromatin states identified by Li et al. (2019) [18]. **(d)** Proportions of MSF and MRF identified in our study overlapping with all fifteen chromatin states. **(e)** Proportions of MSF and MRF overlapping with each of the fifteen chromatin states.

Indeed, the majority of the MSF (67%) and MRF (91%) outliers were located within the annotated TEs (Fig 2a, Table S5 and Fig. S10 in Additional File 1). Expectedly, the proportion of MRFs within TEs was significantly enriched compared to the proportion of MSFs (*p-value* < 2.2 x 10 ^-16^; FET). While a significant enrichment of MRF was found for Class 1 retrotransposons (76% MRF vs. 49% MSF) (Table S5 in Additional File 1), this trend was not consistent for all individual TE families, with Copia transposons being over-represented in the MSFs rather than MRFs (14% MRF vs. 16% MSF, FET, *p*-value = 10^-16^). The MSFs were enriched for Class 2 transposons (16% MSF vs. 13% MRF) (Table S5 and Fig. S10 in Additional File 1; *p*-values < 2.2 x 10^-16^; FET). We detected 1.5 times more CACTA TEs in the MSFs (19% MSF vs. 13% MRF), and a 3-, 4-, and 7- fold increase in the proportion of Mutator, Mariner, and Harbinger TE families in the MSF compared to that in the MRFs (all *p*-values < 2.2 x 10^-16^; FET), suggesting DNA transposons are more frequently found in the accessible regions of the genome (Table S5 in Additional File 1). These results suggest that different classes of transposable elements show different levels of chromatin accessibility or insertion preference. The relative abundance of MRFs and MSFs within different TEs was similar among genomes, with the exception that a lower percentage (44%) of MSF was identified in retrotransposons (Class1) in the D genome compared to that in the A (52%) and B (51%) genomes, (Fig. S10 in Additional File 1; *p-values* < 2.2 x 10^-16^; FET). It should also be noted that 12% of the MSF and 6% of the MRF were located within the unannotated intergenic regions.

Further, we compared the distribution of DNS scores among the genomic regions previously classified into 15 chromatin states using the histone acetylation and methylation epigenetic marks [18]. Overall, the DNS density distributions shifted to positive values for all states, except for state 13, which is enriched for TEs (Fig. 2c, Table S6 in Additional File 1). The states 5-7 enriched for regulatory regions showed elevated DNS scores, which, however, were lower than the DNS scores of chromatin states 1-4 enriched for gene coding sequences. The majority of MSF (56%) and MRF (60%) identified in our study did not overlap with any of the 15 chromatin states (Fig. 2d). In total, these unique MSF covered 99.5 Mbp, with a total of 18,709 MSF (∼1.6 Mb) located within the 2 kb promoter regions of 12,788 high-confidence gene models. Each of the 15 chromatin states, except state 10, showed small overlap (<5%) with MSF (Fig. 2e). Chromatin states 5-7 harbored between 1-2.5% of MSF, covering 1.8 Mb in state 5, 3.4 Mb in state 6, and 4.3 Mb in state 7 (Table S6 in Additional File 1). Nearly 10% of MSF were detected in chromatin state 10 (Fig. 2e, Table S6 in Additional File 1), which was enriched for H3K27me3 histone modification marks [18] involved in facultative suppression of gene expression [40]. Consistent with our earlier analyses, showing that the majority of MRF are detected within the annotated TEs (Table S5 and Fig. S10 in Additional File 1), we found that nearly 31% of MRF are located within chromatin state 13 (Fig. 2e, Table S6 in Additional File 1). However, in spite of detecting the majority of MSF within the annotated TEs (Tables S5, S6 and Fig. S10 in Additional File 1), only a small fraction of MSF mapped to chromatin states 12 (4.3%) and 13 (1.8%). Taken together, these results indicate that the differential MNase digest has the potential to complement the functional annotation of the wheat genome by expanding the map of hyper-sensitive chromatin in the intergenic regions, which was previously shown to be enriched for long-range *cis*-regulatory elements in maize [7].

### Chromatin accessibility in the promoter regions is positively correlated with gene expression

Previous studies demonstrated a strong correlation between the levels of gene expression and epigenetic modifications in wheat [18,29–31]. We investigated the relationship between sensitivity of chromatin to treatment with different concentrations of MNase and gene expression levels. We found that gene expression levels correlate positively with DNS scores in the gene body (*r* = 0.35, *p* < 2.2 x 10^-16^), 500 bp (*r* = 0.22, *p* < 2.2 x 10^-16^) and 2 kb (*r* = 0.21, *p* < 2.2 x 10^-16^) upstream of genes. By comparing the expression levels of genes located within 2 kb from the MSFs and MRFs (Fig. 2b), we found significant differences between these two groups of genes (W= 164,110,000, *p-value* < 2.2 x 10^-16^; Wilcoxon test). Consistent with these observations, on average, genes associated with MSF showed a 30% increase in expression compared to genes located in close proximity to MRF.

In allopolyploid wheat, the contribution of each of the duplicated homoeologous genes to total expression varied across developmental stages and tissues [28, 30]. The set of previously characterized 16,746 syntenic homoeologous gene triplets [30] was evaluated for correlation between gene expression of individual gene copies in a triplet and DNS score in the genic and surrounding regions (Table S7 in Additional File 2, Fig. S11 in Additional File 1). To compare the DNS values among the homoeologous genes, we partitioned a 2 kb region (from −1 kb to +1 kb) around the CDS start site into four 500 bp-long intervals referred to as regions *a*, *b*, *c* and *d* (Fig. 3). The balanced group of gene triplets showed similar DNS profiles around the CDS start sites for each genome (Fig. 3), with a peak at 210 bp upstream of the CDS. The DNS scores for each genome were 0.375, which represents a 19% increase above the average DNS scores for all HC gene models. For the balanced triplets, there was no difference in DNS scores in any of the intergenomic comparisons for these four intervals (*p-values* range from 0.26-89, Kruskal-Wallis test) (Table S8 in Additional File 1).

**Figure 3.**
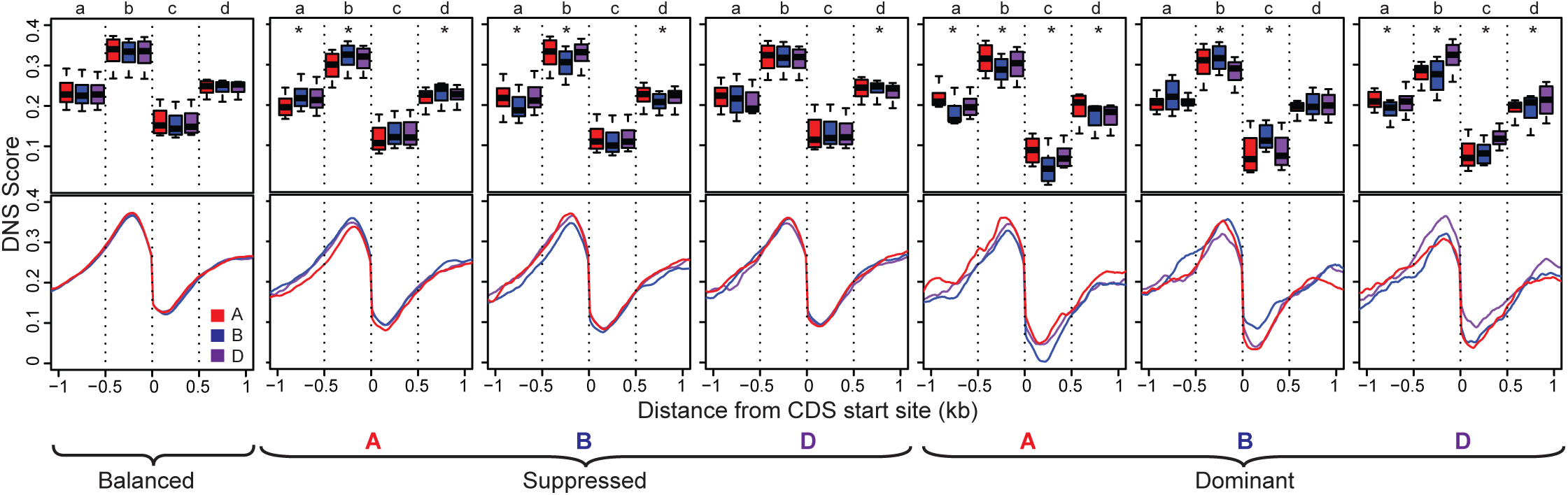
Chromatin accessibility in the homoeologous gene sets showing balanced and unbalanced (suppressed or dominant) expression. Lower panel shows distribution of DNS scores in 10 bp- long windows around the CDS start position from −1 kb to +1 kb. The upper panel shows the distribution of mean DNS scores calculated for 500 bp-long intervals *a* (−1 kb to −500 bp), *b* (−500 bp to 0), *c* (0 to +500 bp) and *d* (+500 bp to +1 kb). Statistically different comparisons of biased expression by region are marked by asterisks.

The suppression of gene expression in either A or B genomes was accompanied by a significant reduction of DNS scores in regions *a, b* and *d* compared to the non-suppressed homoeologous gene copies in other genomes (Table S8 in Additional File 1, *p-values* ≤ 10^-4^; Kruskal-Wallis test). However, for the D genome copies of genes with suppressed expression, a significant reduction in DNS score relative to other homoeologs was observed only in region *d* (*p-values* ≤ 0.01; Kruskal Wallis test). For the gene triplets with one of the genomic copies overexpressed, the corresponding dominant genome had significantly higher DNS scores in regions *b* and *c* (*p-values* ≤ 10^-5^, Table S8 in Additional File 1). These results indicate that the previously observed connection between the biased expression of duplicated genes and epigenetic modification [18,29,30] is consistent with the changes in the abundance of fragile nucleosomes in the promoters or the 5’ ends of genes.

### Chromatin accessibility of genic and intergenic regions along the chromosomes

Our analyses showed that the distribution of overall DNS scores along the centromere-telomere axis shows a strong gradient with the distal chromosomal regions possessing more accessible chromatin than the pericentromeric regions (Fig. 1b). We tested whether the patterns of chromatin accessibility along the chromosomes in the genic regions also mirrored this trend. For this purpose, we assessed the mean DNS score in the gene body, 500 bp and 2 kb upstream of the CDS start positions, 2 kb downstream of CDS end positions, and intergenic regions. When averaged across all genes, the highest DNS score was detected in the 500 bp interval upstream of the CDS, followed by the regions 2 kb upstream and downstream of CDS, then the gene body, and lastly the intergenic regions. The DNS values in these partitions were similar among all three genomes (Table 1, Fig. S12 in Additional File 1). While the chromosomal patterns of DNS distribution in the intergenic regions reflect those observed for the overall DNS score distribution, the DNS scores for the genic regions, remained mostly uniform along the chromosomes, except across the gene body (Fig. 4 and Table S3 in Additional File 1). On the contrary, the gene body DNS score for centromeric genes, on average, was even 1.5-fold higher than that in the other regions (Kruskal Wallis test, *X*^2^ = 646, *p-value* =2.2x 10^-16^) (Table S3 in Additional File 1). There was no detectable DNS difference in the immediate 500 bp upstream of the CDS for any genome or segment (DNS _500bp up_ = 0.26; Kruskal Wallis test, *X^2^*=9.3, *p-value* = 0.054) (Fig. 4, Table S3 in Additional File 1). Even though a significant DNS score difference among the five chromosomal regions was detected within 2 kb from the gene (DNS _2kb up_ = 0.18, Kruskal Wallis test, *X*^2^= 247, *p* =2.2x 10^-16^; DNS _2kb down_ = 0.16, Kruskal Wallis test, *X*^2^= 530, *p* =2.2x 10^-16^) (Fig. 4, Table S3 in Additional File 1), these differences were no more than 10% of the overall mean (fold change range 0.9 - 1.1). These results indicate that while the chromatin accessibility of intergenic regions tends to reduce from telomere to centromere, the chromatin accessibility of genic regions does not follow this trend, and remains mostly stable.

**Figure 4.**
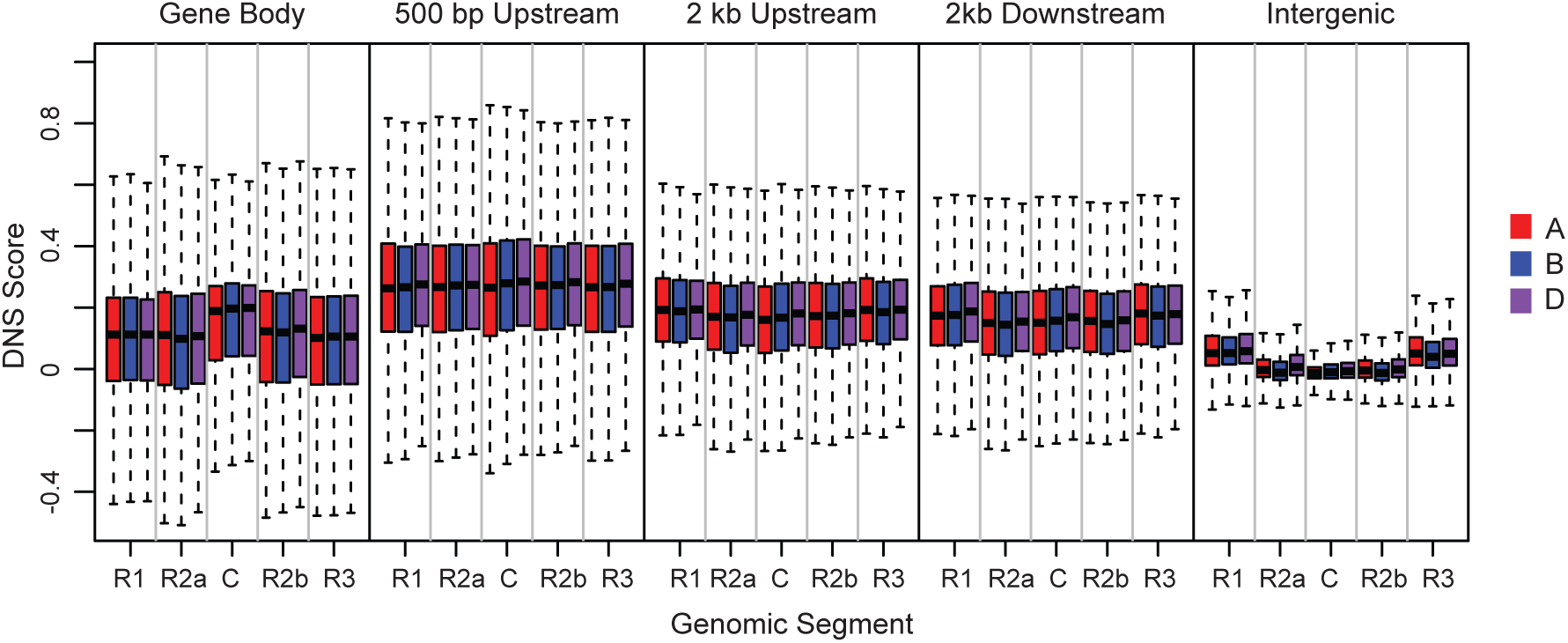
Distribution of DNS scores in the genic (gene body, 500 bp upstream or downstream of CDS, 2 kb upstream or downstream of CDS) and intergenic regions across five chromosomal segments.

### Transposable element frequency is correlated with DNS score

Our results indicate that the chromosome-level distribution of DNS scores is mostly driven by the chromatin accessibility of the intergenic regions, which is mostly composed of TEs [35]. Variation in the distribution of different classes of TEs along the chromosomes was previously reported [34,37,41]. We hypothesized that the inter-chromosomal differences in chromatin accessibility, as well as distribution of chromatin accessibility along the chromosomes is defined by the distribution of TEs. Using the annotated TEs in the wheat genome [34, 35], we evaluated the distribution of DNS scores relative to the distribution of different TE classes across genome. The most abundant class of TEs in the wheat genome is LTR retrotransposons that make up 67% of the genome [34], and the Gypsy superfamily is the predominant LTR, which comprises nearly 50% of the wheat genome. Using a 1-Mb sliding window across the genome, the Gypsy (RLG) superfamily showed a significant negative correlation between TE content and chromatin accessibility across all genomes (ρ_A-genome_ = −0.68; ρ_B-genome_ = −0.64, and ρ_D-genome_ = −0.67) (Fig. 5). On average, genomic regions with Gypsy (RLG) TEs had negative DNS scores in all three genomes, while the other common LTR, Copia (RLC) TEs, had a slightly positive DNS score (Table 2). Overall, the LTR-retrotransposons showed lower chromatin accessibility than the DNA transposons (Table 2, Fig. S13 and Table S9 in Additional File 1; Table S10 in Additional File 3). The DNA transposon regions had positive DNS scores across the TE body, where average DNS score for CACTA TEs was 0.03, and average DNS scores for the less common Mutator, Harbinger and Mariner TEs were 0.08, 0.10, and 0.24, respectively. These results indicate that a broad range of variation in chromatin accessibility exists among different classes and super-families of TEs in the wheat genome, and that chromatin accessibility of any given genomic region to a large extent is defined by the relative abundance of one or another type of TE.

**Figure 5.**
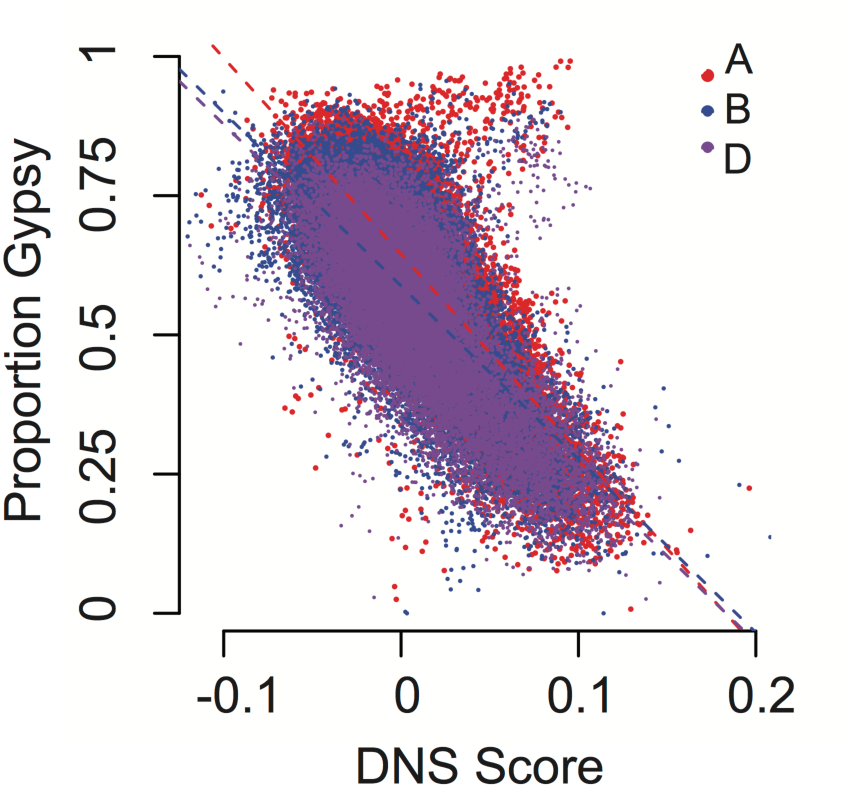
Correlation between TE Gypsy superfamily content and DNS scores. Genomes were split into 1 Mb windows, and correlation between the proportion of the Gypsy TE content and the overall window DNS score was calculated for the A genome (red), B genome (blue), and D genome (purple).

**Table 2.** DNS scores for TE superfamilies

| TE class | TE family | A genome |  | B genome |  | D genome |  |
| --- | --- | --- | --- | --- | --- | --- | --- |
|  |  | Mean<sup>†</sup> | SD<sup>‡</sup> | Mean<sup>†</sup> | SD<sup>‡</sup> | Mean<sup>†</sup> | SD<sup>‡</sup> |
| Class I | RLG | -0.0043 | 0.166 | -0.0131 | 0.268 | -0.00381 | 0.174 |
|  | RLC | 0.0087 | 0.156 | 0.0056 | 0.191 | 0.0146 | 0.152 |
|  | RLX | -0.0217 | 0.185 | -0.0244 | 0.297 | -0.0189 | 0.180 |
|  | RIX | 0.0353 | 0.248 | 0.0262 | 0.230 | 0.0364 | 0.239 |
| Class 2 | DTC | 0.0352 | 0.438 | 0.0248 | 0.981 | 0.0229 | 0.815 |
|  | DTM | 0.0854 | 0.205 | 0.0744 | 0.202 | 0.0770 | 0.195 |
|  | DTH | 0.1106 | 0.250 | 0.0867 | 0.252 | 0.0994 | 0.243 |
|  | DTT | 0.2396 | 0.259 | 0.2318 | 0.261 | 0.2245 | 0.256 |
|  | DTX | 0.2079 | 0.262 | 0.1939 | 0.271 | 0.2151 | 0.261 |
|  | DXX | 0.0736 | 0.329 | 0.0531 | 0.252 | 0.1011 | 0.211 |
| Unclassified | XXX | 0.0171 | 1.862 | -0.0878 | 3.669 | -0.0544 | 3.579 |

<sup>†</sup> DNS mean score for each TE family across the length of the annotated TEs (Wicker, et al 2018)
<sup>‡</sup> standard deviation of DNS score for each TE family

### Chromatin accessibility of TEs is associated with their chromosomal position

To investigate the relationship between the TE distribution and the chromatin accessibility along the wheat chromosomes, we compared DNS scores of different TE superfamilies in the five chromosomal segments R1, R2a, C, R2b, R3. Overall, the patterns of chromatin accessibility in the TE space in the five segments mirror the patterns observed for these regions when all sequences are considered together (Figs. 1b, 6). The chromatin in the TE-harboring regions in the distal ends was shown to be more sensitive to MNase digestion than chromatin in the pericentromeric and interstitial segments. The A and D genomes had nearly a 10-fold increase in DNS score for the TE regions on distal ends compared to the overall TE mean (DNS_A_ = 0.006, DNS_D_ = 0.006), and a 40% reduction in the centromere (Table S3 in Additional File 1). The B genome distal ends showed a 21 and 16-fold increase in the DNS score, and 4-fold reduction in the centromeric regions (DNS_B_= 0.003, Table S3 in Additional File 1). We detected similar increases in the DNS score among the common Gypsy, Copia, and CACTA TE superfamilies in the distal ends, with corresponding reductions in the centromeric regions (Fig. 6, Table S3 in Additional File 1). These findings suggest that while there are differences in the sensitivity to MNase treatment among different superfamilies of TEs, the relative position of the TE on the chromosome also correlates with the accessibility of their chromatin. The overall gradient of chromatin accessibility from centromere to telomere remains consistent for all TE superfamilies.

**Figure 6.**
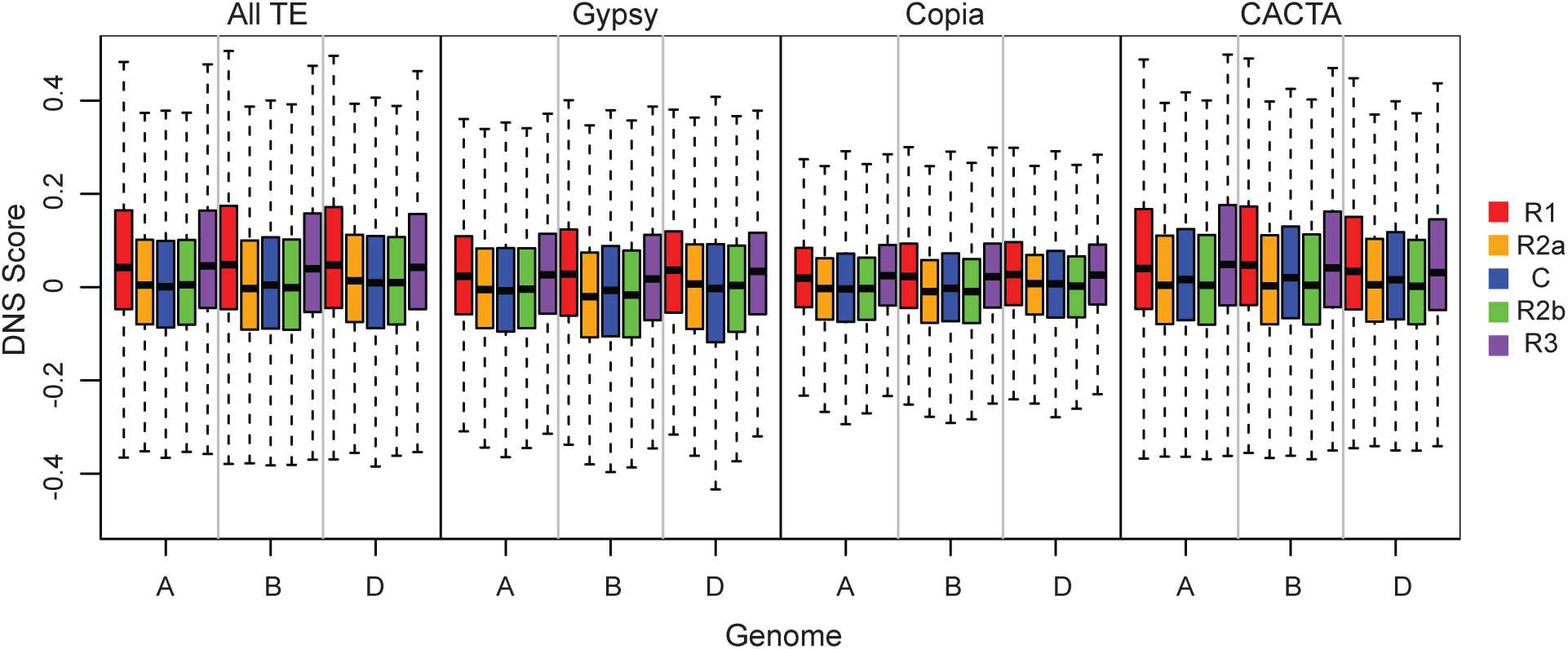
Distribution of DNS scores for distal chromosomal segments R1 (red) and R3 (purple), interstitial segments R2a (orange) and R2b (green), and centromere (blue) for different TE superfamilies.

### Chromatin accessibility of TEs is associated with their proximity to genes

While we observe a highly negative correlation between the incidence of Gypsy superfamily members in a genomic region and chromatin accessibility (Fig. 5), we found that the individual Gypsy families demonstrate variable DNS scores (Fig. S14 in Additional File 1). The Gypsy families with the most negative DNS scores were Nusif (RLG famc4, DNS_all genomes_ = - 0.21), Lila (RLG famc14, DNS_A genome_ = −0.17; DNS_B genome_ = −0.19; DNS_D genome_ = −0.15), and Daniela (RLG famc9, DNS_A genome_ = −0.14; DNS_B genome_ = −0.15; DNS_D genome_ = −0.17), while Sabrina (RLGfamc2, DNS_all genomes_ = 0.07), WHAM (RLG famc 5, DNS_all genomes_ = 0.07) and Wilma (RLGfamc6, DNS_all genomes_ = 0.08) each possess the most positive scores across the TE body in all genomes (Table S10 in Additional File 3; Fig. S14 in Additional File 1).

The chromatin accessibility of the Copia superfamily also showed variability ranging from negative to positive values. For example, the two most common families Angela (RLC_famc1) and Barbara (RLC_famc2) showed slightly positive and negative DNS scores, respectively (Fig. S14 in Additional File 1; Table S10 in Additional File 3). Less frequent Copia families, such as, famc16, had a DNS score of −0.10 in all three genomes, while TE family Bianca (famc12) possesses DNS scores greater than 0.15 in all three genomes (Fig. S14 in Additional File 1; Table S10 in Additional File 3). CACTA family 35 showed the most negative DNS scores ranging from −0.23 in the A genome, to −0.25 in the B genome. The Balduin TE family (DTC_famc8) also showed negative DNS scores across genomes (DNS_A genome_ = −0.06, DNS_B genome_ = −0.16, DNS_D genome_ = −0.15) (Fig. S14 in Additional File 1; Table S10 in Additional File 3). Three other CACTA families, Enac (DTC_famc20, DNS_all genomes_ > 0.17), DTC_famc26 (DNSall genomes > 0.17), and Benito (DTC_famc12, DNS_all genomes_ > 0.20) each had positive DNS scores in the TE regions. Variable patterns of DNS score were observed for all common superfamilies of TEs (Fig. S14 in Additional File 1; Table S10 in Additional File 3), suggesting that processes controlling chromatin structure may have different effects on different TE families.

A previous study demonstrated an enrichment or deficiency of certain TE families in the promoter regions [34]. The Gypsy TEs from Nusif and Daniela families were strongly under-represented in the gene promoters, and also showed some of the lowest DNS scores among the TE families in our dataset (Fig. S14 in Additional File 1; Table S10 in Additional File 3). Likewise, the Copia TEs from the Bianca family that were highly enriched around gene promoters were also among the TEs showing the highest DNS score. Similar trends were detected for the CACTA TEs. Both Enac and Benito that were highly enriched around the gene promoters [34] showed high DNS scores, whereas the DNS scores in the Balduin TEs that were underrepresented in the promoter regions were among the lowest in our dataset (Fig. S14 in Additional File 1; Table S10 in Additional File 3). The observed correlation between the proximity of TEs to genes and their sensitivity to the MNase treatment appear to be consistent with the earlier findings showing the spread of epigenetic modifications near the TE insertion sites [9, 42].

To test this possibility, the DNS scores of TEs from the same superfamily or family located within and outside of the 2 kb promoter regions were compared. Only 2 - 2.5% of the Gypsy TEs were found within the 2 kb regions upstream of CDS, but showed significantly higher (*p-value* < 2.2 x 10^-16^; Mann-Whitney U test) sensitivity to MNase than Gypsy TEs located outside of the promoter regions (Fig. 7a). The difference between Gypsy elements’ DNS score in proximity to genes translates to a 12-, 5-, and 14-fold increase for the A, B, and D genomes, respectively, compared to the DNS scores for the Gypsy elements > 2 kb away from genes. Both Copia and CACTA TEs found within the 2 kb promoter region also showed significantly higher DNS scores compared to the TEs outside of the promoter regions (Fig. 7a). The DNS scores for Copia TEs within promoters ranged from 0.08 to 0.09, while TEs outside of the promoter region had mean scores of 0.005, 0.002, and 0.01 for the A, B, and D genomes, respectively. Likewise, for the CACTA TEs the average DNS score was 0.02 away from genes and 0.18 near genes. Similar trends were observed for the less common TE superfamilies including the LINE (RIX), and unclassified LTR retrotransposons (RLX), Mutator (DTM), Harbinger (DTH), Mariner (DTT), and the unclassified DNA transposons (DTX and DXX) (Fig. S15 in Additional File 1), suggesting that the accessible chromatin characteristic of the genic regions also extends to the neighboring TEs.

**Figure 7.**
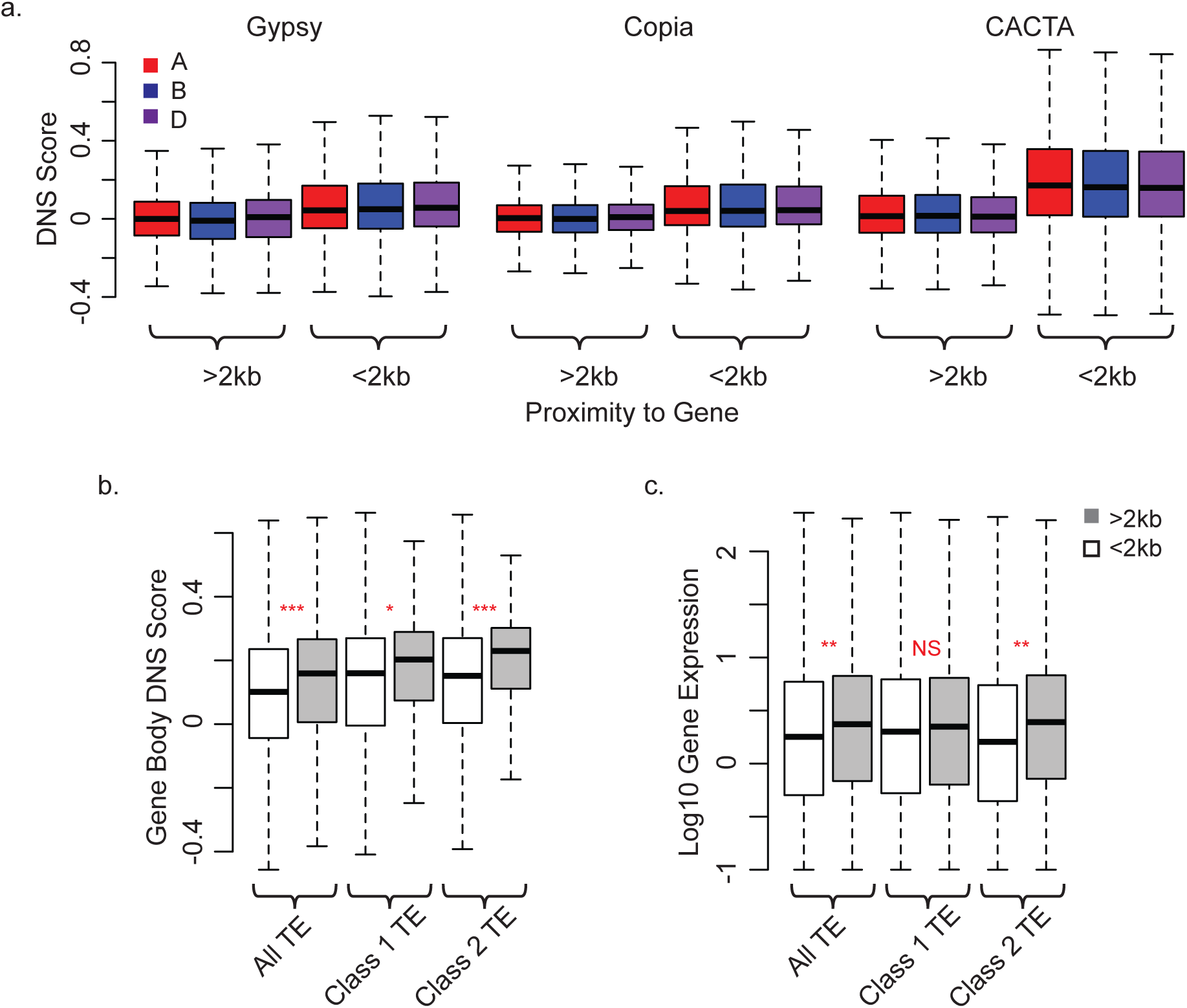
Chromatin accessibility of TEs and neighboring genes. **(a)** DNS scores by genome for TE super-families located within and outside of 2 kb promoter regions of the genes for Gypsy superfamily, (Mann-Whitney U Test: W_A_= 85484000, W_B_ = 81997000, W_D_=6067000; all *p*-values < 2.2 x 10^-16^); Copia superfamily (W_A_ = 60126000, W_B_ = 60412000, W_D_ =56955000; all *p*-values < 2.2 x 10^-16^); and CACTA superfamily (W_A_ = 91146000, W_B_ = 113700000, W_D_ = 86315000; all *p*-values < 2.2 x 10^-16^). **(b)** DNS scores across gene bodies for all genes that have a TE located within 2kb of gene or more than 2kb from gene. Distributions are given for all TEs (W= 39630000, *p*-value < 2.2 x 10^-16^), class 1 TE (W=8089200, *p*-value = 2.7 x 10^-05^), and class 2 TE (W=7616500, *p*-value < 2.2 x 10^-16^). *p*-values are denoted by red stars above boxplot for each comparison: * 0.05 < *p* <10^-5^; ** 10^-5^ < *p* <10^-10^, *** 10^-10^ < *p* <10^-16^, NS= p > 0.05 **(c)** Log10 gene expression values for all genes that have a TE located within 2 kb of gene or more than 2 kb from a gene. Distributions are given for all TEs (W= 4768500, *p*-value = 1.5 x 10^-07^), class 1 TE (W=94150, *p*-value = 0.11), and class 2 TE (W=908540, *p*-value = 2.5 x 10^-8^).

We further investigated the impact of TE insertion into the promoter regions < 2 kb or > 2 kb away from a gene on chromatin accessibility of the gene body and the levels of gene expression (Fig. 7b). We found that the insertion of both LTR and DNA TEs into the promoter regions < 2 kb from a gene coincided with both decreased gene body chromatin accessibility and reduced gene expression (Fig. 7b, c).

### Gene spacing correlates with chromatin accessibility in the intergenic regions

Given the significant differences across chromosomes in chromatin accessibility in the intergenic regions compared to genes (Fig. 1b, Fig. 4, Table 1), and the relationship between TE proximity to genes and TE chromatin accessibility (Fig. 7; Fig. S15 in Additional File 1), it is possible that the physical spacing of genes along the chromosomes could be one of the factors influencing the global distribution of chromatin sensitivity to MNase digest along the centromere-telomere axis. On average there is an 8-, 5-, and 6-fold increase in gene density in the distal chromosomal regions compared to the centromeric regions for the A, B, and D genomes, respectively [35]. To assess the relationship between the gene density and chromatin accessibility, the DNS scores were compared among four intergenic interval ranges defined based on the physical distances between the adjacent genes: <10 kb, 10 kb - 100 kb, 100 kb -1 Mb, and >1 Mb. The mean DNS scores were calculated for each intergenic interval in 1-kb windows, until the midpoint of intergenic distance of adjacent genes is reached (Fig. 8a).

**Figure 8.**
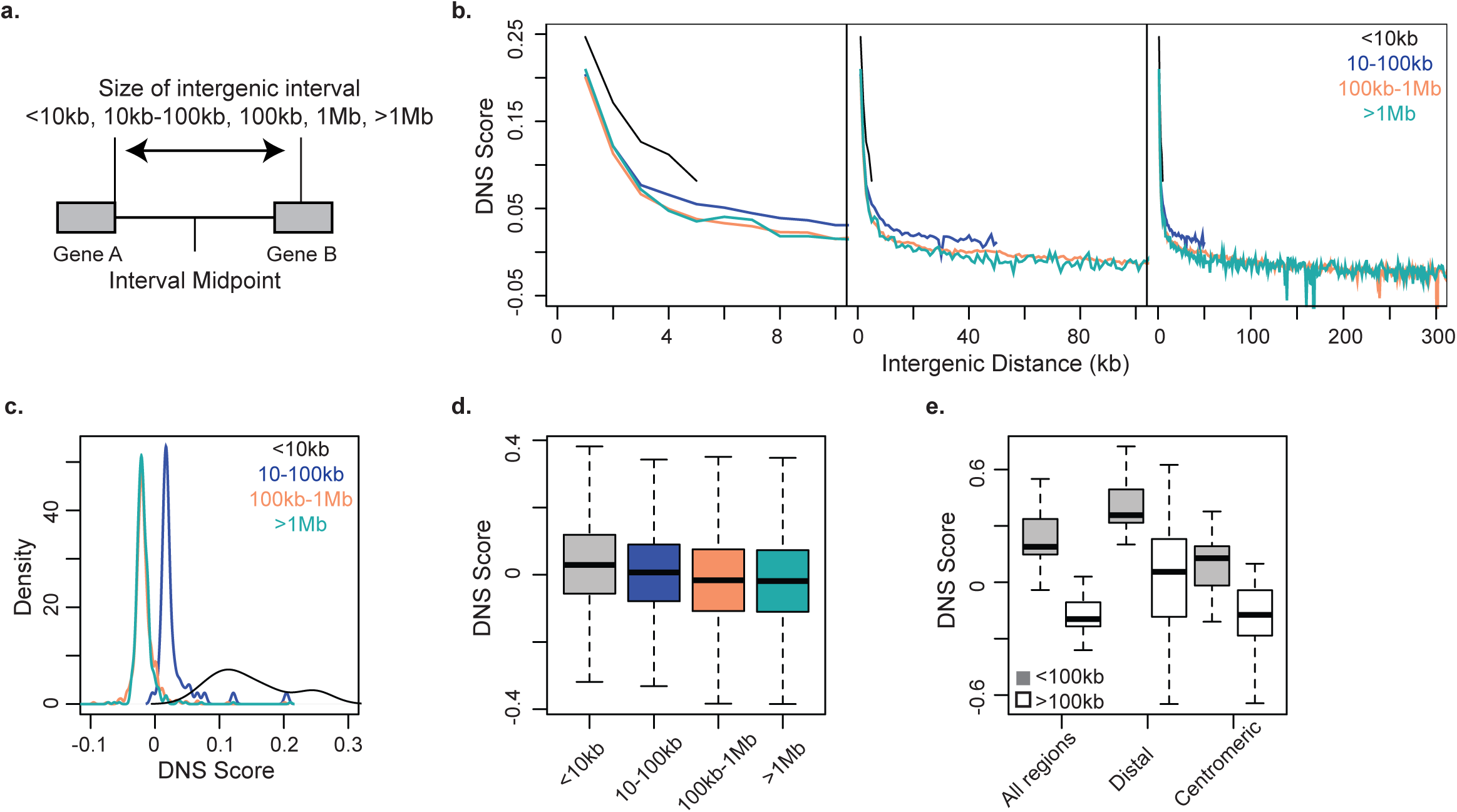
Relationship between the gene spacing and sensitivity to MNase treatment. **(a)** All intergenic regions were classified into 4 groups based on the distance between the adjacent genes: <10 kb (black), 10 kb – 100 kb (blue), between 100 kb −1 Mb (apricot), and >1 Mb (teal). **(b)** DNS scores for three groups of intergenic distances shown on scale up to 10 kb (**left panel**), 100 kb **(middle panel)** and 300 kb (**right panel**). **(c)** Density plot showing DNS distribution for 1 kb windows for each intergenic distance group. **(d)** Distribution of DNS scores for all Gypsy LTRs for the same intergenic distance group, representing a significant effect of intergenic distance on Gypsy LTR DNS score (*X^2^* = 23266; *p-value* < 2.2 x 10^-16^, Kruskal Wallis test). **(e)** Distribution of DNS scores for intergenic segments < 100kb (gray) and more than 100kb (white) from neighboring genes for all regions in the genome, distal segments only, and centromeric segments only. Comparisons result in significant differences for all 3 categories (W_all segments_ = 3023, *p-value* < 2.2.x 10^-16^; W _Distal._= 2661, p-value 6.7 x 10^-12^; W _Centr._=2118, *p-value* = 2.9 x 10^-4^).

For intergenic intervals <10 kb, the DNS score within the first 1 kb window was higher than that for larger intergenic intervals (Fig. 8b), and decreased to a value of 0.08 at the midpoint of the neighboring genes. DNS scores for intergenic intervals where neighboring genes are located more than 10 kb apart reach this value (DNS= 0.08) within 3kb, and continue to decrease as intergenic distance increases between genes (Fig. 8b). For intergenic intervals 10 kb - 100 kb, 100 kb −1 Mb, and >1 Mb, DNS scores eventually reach a DNS value that does not change with distance. We refer to this point as the background DNS score, which represents the peak of distribution of DNS scores for each intergenic interval (Fig. 8c). Our results show background DNS values were similar for 100 kb −1 Mb, and >1 Mb intervals (DNS = −0.017), however intergenic intervals within the 10-100 kb range showed a higher background DNS score (DNS = 0.018) (Figs 8b, c; Table S11 in Additional File 1), suggesting that once genes are located more than 100 kb from its neighboring gene, the chromatin state is predominantly inaccessible and does not change significantly with more intergenic distance.

We further investigated the effect of intergenic interval sizes on the DNS score distribution among Gypsy TEs, the most common TE superfamily in the wheat genome (Fig. 8d). We detected a significant difference (*X^2^*= 23,266, *p-value*=2.2 x 10^-16^, Kruskal Wallis test) in the DNS score of Gypsy TEs when they are located in intergenic intervals of different sizes. Gypsy TEs located within the larger intergenic intervals showed a lower sensitivity to MNase digest than those located within the intervals of smaller size, suggesting some connection between the physical spacing of genes in the wheat genome and chromatin accessibility of TEs in the intergenic intervals.

A confounding effect on the chromosomal gradient of DNS scores is the difference in gene density and ultimately intergenic distance between adjacent genes in the distal and centromeric regions. The average intergenic distance between genes in the distal regions is 69.9 kb, while in the centromeric region it is 418.3 kb; this results in the intergenic intervals on the distal ends predominantly falling into the 10 kb - 100 kb range (Fig. S16 in Additional File 1), while the majority of the intergenic intervals in the centromeric region fall into the 100 kb −1 Mb range. By selecting a random sample of intergenic distance intervals across all genomic regions from the <100 kb range, and >100 kb range, we confirmed a significant difference in chromatin accessibility based on intergenic distance (Fig 8e, *p-value* < 2.2.x 10^-16^ Kruskal Wallis test). Further, to remove the confounding effect of position along the centromere-telomere axis on our estimates of chromatin accessibility in these intergenic intervals, we compared DNS scores of intergenic intervals between these two distance ranges separately for the distal and pericentromeric regions of the chromosomes. We found significant differences in DNS scores (*p-value* < 2.2.x 10^-16^, Mann-Whitney test) for intergenic intervals < 100 kb and >100 kb in both comparisons, mirroring the chromosomal gradient in chromatin accessibility (Figs. 1, 8e). Similar results were obtained by taking random samples of each of the intergenic intervals with size ranges of 10 kb - 100 kb, 100 kb −1 Mb, and >1 Mb from distal and centromeric regions (Table S11 in Additional File 1). These results suggest that, in addition to the correlation observed between the region’s DNS score and its position on the centromere-telomere axis, there is a connection between the chromatin accessibility and distance between genes.

However, the established relationships among these factors are not always straightforward. For example, even though we observed a 2.5-fold reduction in DNS score for the R1 distal region on chromosome 4A (Fig. 1c), in comparison to the same region on chromosome 4B, the average distances between genes in these regions were very similar, with average distances on 4A and 4B being 79.7 kb and 76.4 kb, respectively. This trend on the chromosome 4A-R1 region coincided with 1.5-fold increase in the proportion Gypsy TEs, and 1.5-fold reduction in the proportion CACTA TEs, compared to the R1 regions from other A genome chromosomes, indicative of a connection between chromatin accessibility and the TE composition of the intergenic regions.

### Chromatin accessibility of the wheat centromeric regions

Centromeric chromatin is formed by nucleosomes where histone H3 is replaced by its centromeric variant CENH3 [43]. In wheat, centromeric nucleosomes are associated with centromeric satellite sequences mostly composed of LTR transposable elements from the Cereba family, which is also enriched in the centromeres of barley chromosomes [34,44,45]. While the locations of the centromeres on the wheat chromosomes mostly conincided with the regions enriched for Cereba-like elements, the location of the centromere on chromosome 4D in the reference cultivar Chinese Spring was repositioned from that of the Cereba-like repeats [46]. Even though the centromere is composed of condensed heterochromatin, centrometric chromatin in *Drosophila* and yeast showed regions of high sensitivity to MNase digest [43]. We used the differential digest with MNase to investigate the chromatin accessibility around the centromeric regions of wheat chromosomes identified by chromatin immunoprecipitation with antibodies against CENH3 [35, 44].

In most cases, the depth of read coverage under both light and heavy digest with MNase was lower in the centrometric regions than in other chromosomal regions (Fig. 9; Fig. S17 and Table S12 in Additional File 1). These regions of low read coverage also conincided with a high frequency of Cereba LTR transposons, and increased DNS score. The latter is the result of higher read coverage obtained by the centromeric chromatin digest with a low rather than a high concentration of MNase digest. The increased DNS score peak appears to be one of the characteristic signatures of centromeric chromatin in wheat.

**Figure 9.**
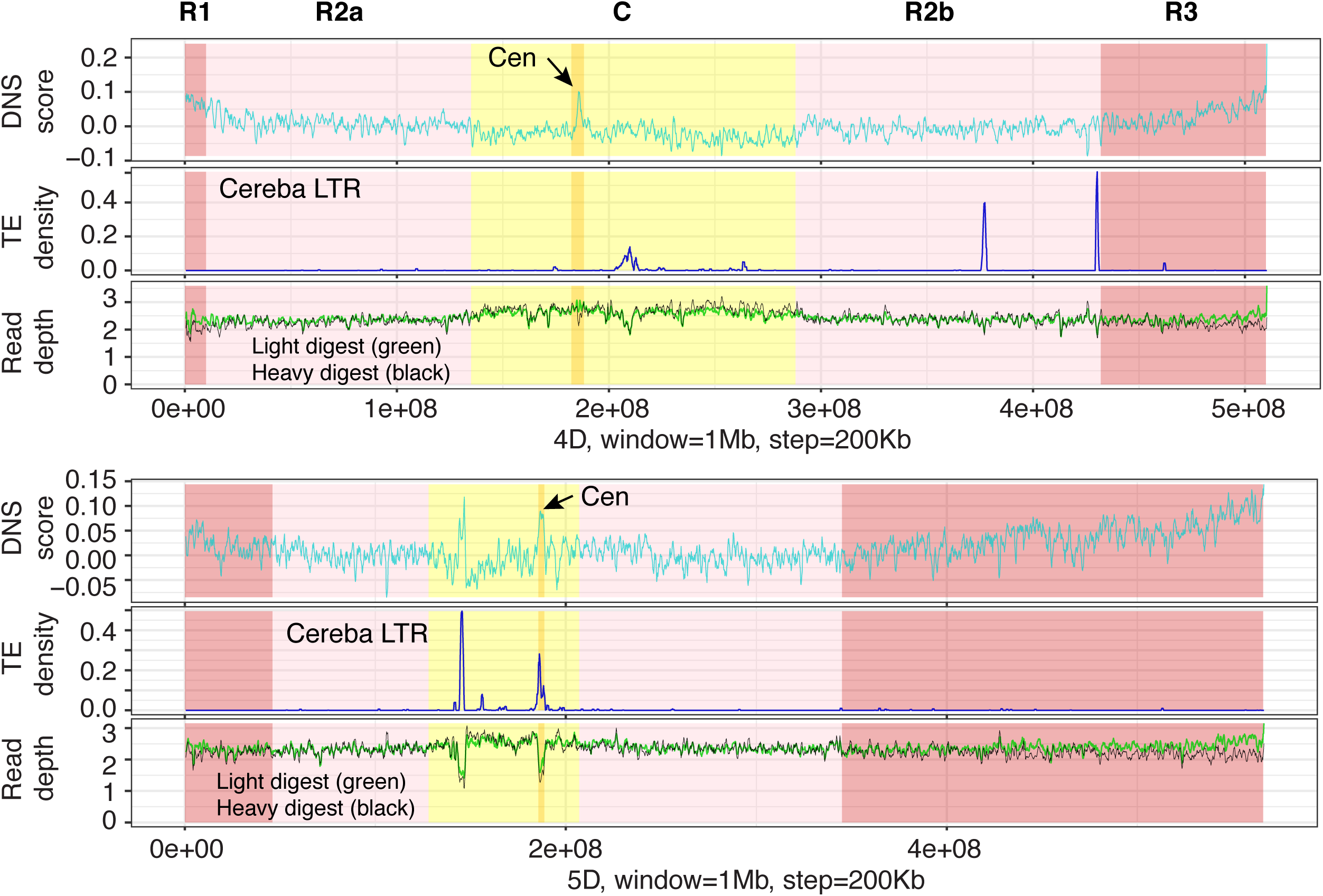
Sensitivity of centromeric chromatin to differential MNase digest. Distribution of DNS scores (top panel), Cereba LTR density (middle panel), and depth of read coverage obtained by light and heavy MNase digest (bottom panel) for chromosomes 4D and 5D. Each chromosome is divided into five regions R1, R2a, C, R2b, and R3. The location of centromere (Cen-dark yellow) is based on previously published studies [35] that used anti-CENH3 chromatin immunoprecipitation method.

On chromosome 4D, the lowest depth of read coverage and increased transposon density located at position 209.4 Mb did not conincide with the location of centromere, which was identified between positions 182.3 and 188.2 Mb by CENH3 localization (Table S12 in Additional File 1). Contrary to that, the DNS score peak was located at position 185.2 Mb within the centromeric region confirming the ability of differential digest with MNase to accurately identify centromeric chromatin. The only exception from these dependencies was found on chromosome 5D, which possesses two regions of increased Cereba transposon density, both showing decreased read depth coverage and increased DNS score peaks. However, only one of these two regions was consistent with the CENH3 centromeric chromatin localization (Fig. 9).

### Partitioning genetic variance among genic regions with different chromatin accessibility

A previous study in maize demonstrated that MSF regions harbor SNPs that explain up to 40% of the phenotypic variance for major agronomic traits [4]. To investigate the impact of chromatin states within genic regions on the phenotypic variation in wheat, we ranked the entire wheat genome from the most closed to most open regions by DNS score, and binned them into 5 bins, each comprising 20% of the genome. SNPs from the 1000 exome project [47] that are located in gene bodies or within 1 kb flanking region of the genes were extracted and placed into their representative DNS score bin. We compared the proportion of phenotypic variance explained by the SNPs from the bin with the most closed chromatin to that explained by SNPs from the bin with the most open chromatin. The genetic variance was estimated using the GCTA-GREML method for plant height, grain filling period, harvest weight, and stress susceptibility (Fig. 10). Consistent across all analyzed traits, an increase in chromatin accessibility was associated with an increase in the proportion of phenotypic variance explained (Table S13 in Additional File 4, Table S14 in Additional File 1). On average across all traits, the proportion of phenotypic variance explained by the SNPs located within the most open chromatin regions was more than 3-fold greater than the variance explained by SNPs located in regions with the most closed chromatin (0.56 open to 0.17 closed, 69% increase) (Fig. 10).

**Figure 10.**
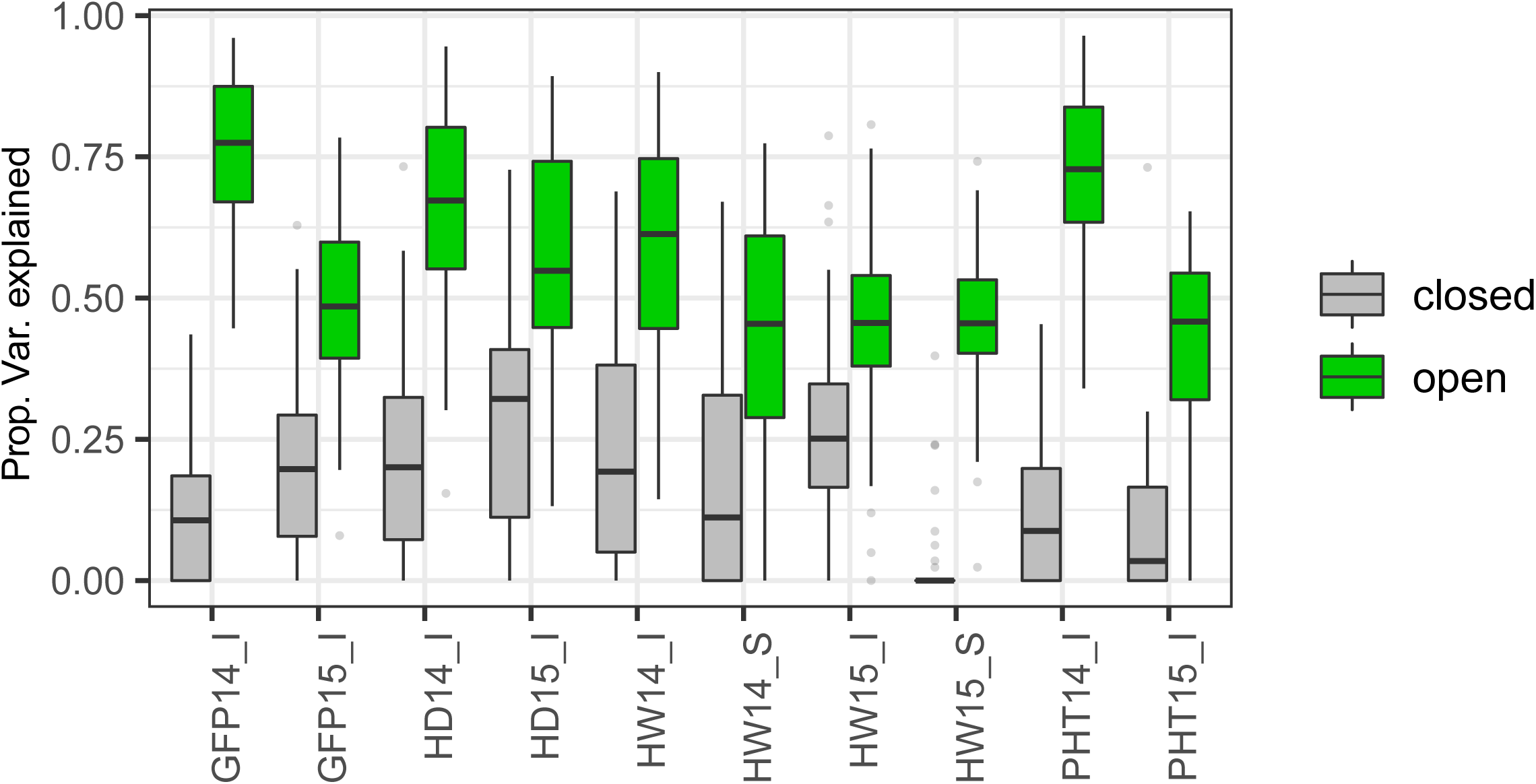
Proportions of phenotypic variation explained by SNPs located in the genomic regions with low (closed chromatin - gray) and high (open chromatin - green) levels of chromatin accessibility. The DNS scores were used to assign each genomic region to the top and bottom 20^th^ percentiles of DNS score distribution. The estimates of variance were performed for plant height (PHT14_I, PHT15_I), grain filling period (GFP14_I, GFP15_I), harvest weight (HW14_I, HW15_I), and stress susceptibility (HW14_S, HW15_S) traits collected for field trials perfromed in 2014 and 2015 [47].

## Discussion

Our results show that the functional and structural features of individual wheat genomes and chromosomes are reflected in the patterns of chromatin accessibility assessed by the differential MNase digest. The overall chromatin accessibility of the wheat D genome, which merged with the AB genomes about 10,000 years ago, was substantially higher than that of the A and B genomes that merged less than 1.3 million years ago [27,48–51]. The intergenomic differences in chromatin accessibility in the D genome were observed across all genomic regions including gene coding sequences, regions upstream and downstream of genes, and intergenic regions. These observations are consistent with the lower abundance of repressive H3K27me3 histone marks across the gene body and a higher gene expression level in the D genome [30]. The post-hybridization accumulation of epigenetic changes over time [32,52,53] is one of the possible factors that might influence genome-level chromatin states, resulting in higher levels of chromatin accessibility in the D genome. However, a lack of substantial differences among the genomes in the proportion of methylated CpG, CHH, and CHG sites [31] indicates that an increase in the D genome’s chromatin accessibility is not directly associated with differential DNA methylation.

Another likely factor is the composition and relative abundance of the repetitive portion of the genome. The D genome is nearly 1 Gb smaller than the A and B genomes due to the loss of 800 Mb of Gypsy LTR TEs [34, 54], which in our study showed the strongest negative correlation with chromatin accessibility in all three wheat genomes. Overall, TEs in the D genome tend to be younger than TEs in the other genomes [34], which is indicative of more recent TE activity in the D genome lineage compared to that in the A and B genomes. The increased gene density and its accompanying reduction in intergenic region sizes [35], and the spread of accessible chromatin states from genes to surrounding TEs we observed in our study, likely contribute to the overall increased chromatin accessibility in the wheat D genome.

Differential MNase-seq revealed a number of genomic regions detectable only under light digest conditions [3, 4], and occupied by either transcription factors or nucleosomes with conformations making linker DNA more accessible. We detected an abundance of MSFs in the proximal regulatory regions, where the levels of chromatin accessibility showed a positive correlation with gene expression [1,3,55,56], and were predictive of the unbalanced expression levels among the duplicated homoeologs, supporting the results of chromatin accessibility studies conducted using the DNase-seq and ATAC-seq approaches in wheat [18, 29]. However, the majority of MSF (67%) identified in our study were located in the TE-rich intergenic regions, with an average distance of 137 kb from the closest gene, and overlapped with annotated TEs. These results are consistent with the prevalence of putative distant *cis*-regulatory elements in crops with large genomes (maize, barley) [7,12,21], and combined with the demonstrated regulatory potential of TE elements derived from regions with open chromatin [7, 22], suggest that genome size expansion driven by TE proliferation in the wheat genomes has the potential to diversify gene expression regulatory pathways. However, TE space expansion does not directly correlate with the MSF frequency, which appears to be conditioned by the regional gene density. The enrichment of MSF in the distal chromosomal regions with high gene/low TE density compared to the MSF frequency in the pericentromeric regions characterized by low gene/high TE density is consistent with this hypothesis, and is supported by the finding that total size of distant accessible chromatin regions (dACRs) with regulatory potential does not scale linearly with an increase in genome size [12].

We found that the chromatin accessibility of neighboring TEs and genes, and the levels of gene expression tend to correlate. The TEs located closer to genes have higher levels of chromatin accessibility than respective TEs from the same family located farther from genes. Likewise, genes having TEs located within 2 kb from the start site tend to have lower chromatin accessibility and expression levels than genes having TEs located more than 2 kb from the gene. We also observed an effect of TE type on chromatin accessibility in the promoter regions, with some families from the Class 1 and Class 2 TEs showing lowest and highest chromatin accessibility, respectively. Our results indicate that cellular mechanisms aimed at maintaining silenced TEs and active gene expression are sensitive to both the physical spacing between these two genomic features, as well as to the types of TEs, and appear to be consistent with models in which transcriptional activation or suppression of TEs can affect the expression of adjacent genes [57, 58] through epigenetic mechanisms [9, 59]. In addition, it appears that the rate of transition from the accessible to inaccessible chromatin states between genic and repetitive intergenic regions is also affected by the physical spacing between genes, with a faster rate of transition in the pericentromeric regions that have lower gene density. Taken together, these observations suggest that the size and composition of intergenic regions might play an important role in shaping the organization of the expressed portion of the wheat genome and its regulation.

Our study shows that the distribution of chromatin accessibility in the intergenic regions follows a negative gradient along the centromere-telomere axis consistent with the previously defined five chromosomal segments with distinct patterns of gene density, expression, recombination rate and diversity: two distal (R1, R3), two pericentromeric (R2a, R2b) and one centromeric (C) [37]. While this chromatin accessibility gradient was consistent for all major classes of TEs, it was not observed for the genic regions indicating that the distribution of chromatin states in the intergenic sequences along the chromosomes have little effect on the chromosomal patterns of chromatin accessibility within the genes. We suggested that the chromosome position could be one of the factors that influence the large-scale chromatin accessibility trends across the genome. However, the analysis of the structurally re-arranged chromosome 4A, where the distal R1 region is made up of the former interstitial R2b region [38, 39], demonstrated that the previously established chromatin states remain mostly unchanged after relocation to a different chromosomal position, suggesting that the distribution of chromatin accessibility along the chromosomes is driven by factors other than the position on the centromere-telomere axis alone. The lack of substantial changes in chromatin since the occurrence of chromosome 4A’s structural re-arrangement suggests that global chromatin states tend to remain stable, at least within short evolutionary time scales, and are primarily defined by the sequence composition of a genomic region.

One of the likely factors that underlies the chromosomal patterns of chromatin accessibility is the physical spacing between genes. The recent analyses of TE composition in the wheat genome found that in spite of the lack of sequence conservation in the intergenic sequences among the wheat homoeologous chromosomes the physical spacing between genes remains conserved [34], indicating the importance of this factor for genome organization and function. Our results show that chromatin accessibility in the intergenic regions decays as a function of distance from a gene, with the rate of decay positively influenced by the physical distance to the adjacent gene. It appears that longer intergenic regions harboring a larger number of TEs are more effectively targeted for TE silencing and chromatin suppression than shorter intergenic regions, thereby creating a gradient of chromatin accessibility along the telomere-centromere axis. These results are in line with the predictions based on the modeling of TE propagation, response of a host genome to TE propagation, and accumulation of silenced TEs in a host genome [60]. This model suggests that lower TE deletion rates, resulting in TE accumulation in genome, could lead to more effective silencing of duplicated TE copies through siRNA-mediated DNA methylation pathway, thus increasing the genome size [60]. However, this model does not explain the origin of a gene density gradient along the telomere-centromere axis and preferential accumulation of TEs in the pericentromeric regions. If TE insertion near genes is negatively selected due to its detrimental effects on gene expression [59], and at the same time TE retention is under positive selection as a part of pathways needed to control TE proliferation [60], one might suggest that TE distribution is defined by the efficiency of selection in different parts of a genome, which in turn is strongly influenced by recombination rate. Strong suppression of recombination in the pericentromeric regions of large wheat chromosomes, and associated with this reduction in the efficiency of selection [61, 62] could potentially create conditions for the disproportionate accumulation of TEs in the pericentromeric regions compared to that in the distal regions. This factor in turn could be responsible for the chromatin accessibility gradient along the wheat chromosome arms. Whether this chromatin architecture plays any functional role or it is simply the consequence of gene spacing distribution remains unclear, but considering recent reports that showed the involvement of intergenic TEs in the developmental regulation of 3D chromatin architecture and gene expression [7,11,12,21,22,58], as well as the evolutionary conservation of gene spacing in the wheat genome [34] and the abundance of accessible chromatin / MSF in the intergenic space [7,12,21], it is possible that such chromatin organization is of functional importance. Further studies incorporating comparative 3D chromatin structure analysis will likely shed some light on the functional role of chromosomal patterns of chromatin accessibility observed in our study.

Based on the DNS-seq read coverage, the lowest levels of chromatin accessibility in our dataset were observed for the wheat centromeres. However, we found that the wheat centromeric nucleosomes have regions that are more sensitive to light than heavy digest conditions. This observation probably reflects the previously demonstrated unconventional conformation of centromeric nucleosomes carrying the CENH3 variant of H3 histone [43]. This differential sensitivity to MNase concentration produced the highest local DNS score peaks in the centromeric regions that in nearly all cases coincided with the previously detected CENH3 signals [35, 44]. This trend was still consistent even for wheat chromosome 4D, which showed repositioning of the CENH3 signal location among different wheat cultivars [46]. On all chromosomes, except 4D, the highest DNS score peaks coincided with an increased density of Cereba LTR elements and CENH3 signal. On chromosome 4D of cultivar Chinese Spring, CENH3 centromeric signal overlapped with the DNS score peak, and was shifted away from the Cereba LTR density peak. Unusual patterns of read coverage, DNS score, Cereba LTR, and CENH3 signals were observed on chromosome 5D, which showed two well-separated peaks for DNS score, Cereba LTR density, and read coverage. However, only one of these regions overlapped with a CENH3 signal detected using CENH3 immunofluorescence [46] and immunoprecipitation [35, 44], suggesting that not in all cases coincidence of DNS score and Cereba LTR density peaks is predictive of centromere location.

Here, we showed that chromatin accessibility is a strong predictor of the effect of SNP variation on phenotype, indicating that the developed map of chromatin states across the wheat genome is useful for prioritizing SNPs in genomic selection experiments or detecting causal SNPs in gene mapping studies or GWAS. Consistently, the regions of the maize genome with high chromatin accessibility harbored SNP variants explaining a substantial proportion of phenotypic variance for a number of agronomic traits [4,7,12]. The value of chromatin accessibility data for detecting causal genomic regions was also previously demonstrated for maize where DNase I chromatin accessibility was used to predict distantly located enhancers genome-wide and for the *b1*, *bx1* and *tb1* genes [7,12,21].

## Conclusions

The chromatin accessibility map of the wheat genome reflects the distribution of functional and structural features across the wheat genome and reveals a close connection between the repetitive and gene-coding sequences that have the potential to influence gene expression regulation. The state of chromatin is one of the dimensions in the genome-to-phenome maps being constructed connecting genomic variation with the molecular-, tissue- and organism-level phenotypes [63]. The relevance of this dimension for effective translation of genomic variant effects to phenotypes has been demonstrated by the enrichment of functionally active genomic elements in the regions with accessible chromatin and an increased proportion of phenotypic variation explained by SNPs from these regions. By combining the developed chromatin accessibility map with other functionally relevant genomic attributes (transcriptome, metabolome, proteome etc.) we can both improve our ability to predict phenotypic outcomes of any particular genome, and select genomic targets for engineering a biological system to obtain the desired effects.

## Methods

### Nuclei Isolation and Differential MNase Digestion

Wheat cultivar Chinese Spring was grown in greenhouse conditions with 16:8-hour light:dark cycle. Two-week old leaf tissue was collected and immediately flash frozen in liquid nitrogen. Nuclei were isolated using a modified protocol by Vera et al [3]. Briefly 4 g of frozen tissue were ground using mortar and pestle under liquid nitrogen and were cross linked for 10 minutes in ice cold fixation buffer (15 mM PIPES-NaOH, pH 6.8, 80 mM KCl, 20 mM NaCl, 0.32mM sorbitol, 2 mM EDTA, 0.5 mM EGTA, 1 mM DTT, 0.15 mM spermine, 0.5 mM spermidine, 0.2 200 µM PMSF, and 200 µM phenanthroline, and 1% formaldehyde). The cross-linking was stopped by adding glycine to a final concentration of 125 mM and incubating at room temperature for 5 minutes. Nuclei were isolated by adding Triton-X 100 to a final volume of 1% and rotated for 5 minutes, then filtered through 1 layer of miracloth. Nuclear suspensions were divided in 2 aliquots and then suspended in 15mL of 50% volume:volume Percoll:PBS cushion, then centrifuged for 15 minutes at 4°C at 3,000 x g. Nuclei were transferred from the Percoll interphase to a new tube, diluted 2X in PBS buffer, and pelleted by centrifugation for 15 minutes at 4°C 2,000 x g. Pellets were resuspended in 15mL of ice cold MNase digestion buffer (50 mM HEPES-HCl, pH 7.6, 12.5% glycerol, 25mM KCl, 4mM MgCl_2_, 1mM CaCl_2_), and pelleted again by centrifugation for 15 minutes at 4°C 2,000 x g. Pellets were resuspended in 2 mL of MNase digestion buffer. A 100 µL aliquot of the resuspended nuclei were stained with 1µg/mL DAPI in PBS buffer, and quantified using hemacytometer on a confocal microscope.

The remaining nuclei were split into 60µL aliquots containing 3,000 nuclei each and flash frozen in liquid nitrogen. Nuclei were digested by micrococcal nuclease (NEB) using 100 U/mL (heavy) and 10 U/mL (light) for 20 minutes at room temperature. Digestion was terminated by adding 10mM EGTA. To break the cross-links, digestions were treated overnight at 65°C in 1% SDS and 100 µg/mL proteinase K. DNA was extracted using phenol-chloroform extraction and precipitated in ethanol. Digested DNA was resuspended in 40 µg/mL RNaseA (Qiagen), and run on a 1% agarose gel to confirm the heavy and light digest.

### Biological replicates and libraries for sequencing

Two separate biological replicates of nuclei were thawed to room temperature and split into 8 separate 60ul aliquots. For each replicate, 4 separate light digestions (10 U/ml) and 4 separate heavy digestions (100U/ml) were carried out for 20 minutes at room temperature. Digestions were stopped with the addition of 0.5M EGTA. DNA was extracted in the same manner described above, and then the 4 samples of each like digestion were combined to produce 2 replicate light digestions, and 2 replicate heavy digestions, resulting in 4 total libraries. Prior to library preparation digested DNA samples were subjected to 100-200 bp size selection using the Pippin prep system (Sage Science). The DNA-seq libraries were constructed from 500 ng of size selected DNA with the GeneRead DNA library I core kit (Qiagen, cat #180434) and GeneRead Adapter I set B (Qiagen, cat # 180986) according to Qiagen protocol with one exception: seven PCR cycles were performed for the library enrichment. The sizes of resulting libraries were validated on the 7500 DNA Bioanalyzer chip. To test the quality of library preparations, two out of four barcoded libraries prepared using the high and low concentrations of MNase were pooled in the equimolar amounts and sequenced with 2 x 75 bp Illumina MiSeq run using MiSeq 150 cycles reagent kit v3. Then each of the four libraries was sequenced on 2 lanes of HiSeq 2500 system (8 lanes total) using a 2 x 50 bp sequencing run producing a total of 1,749,823,029 reads.

### Data Processing and DNS Score Calculation

Raw fastq files were run through quality control using Illumina NGSC Toolkit v2.3.3, and aligned to Chinese Spring RefSeqv1 genome [35] using the HISAT2 v2.0.5 alignment program [64]. Paired end reads were retained if 70% of the read length had a quality cutoff score of ≥ 20. Only uniquely mapped reads were retained for further analysis. BED files were made from each alignment, using the Bedtools v2.26.0 bamtobed [65] conversion to get the coordinates where reads align, then depth of reads was measured in 10bp intervals using bedmap--count option. The read coverage (number of reads that map) for each 10bp interval was normalized by taking the total number of reads mapped for the whole genome and then dividing by million. To get the differential MNase score for each 10bp interval, we subtracted the normalized depth of coverage of the heavy digest from the normalized depth of coverage of the light digest [4]. For instance, for each 10 bp interval on a chromosome we obtain normalized depth of coverage for both light and heavy digests for each replicate, and then calculate the differential depth for each replicate (2 reps). Correlation between the replicates was 0.98 (*p-value* < 2.2x 10^-16^) (Figs S1a,b in Additional File 1), therefore for estimates across chromosomes, segments, and windows, the mean values of the reps are presented in plots and tables. Negative scores reflect DNS hyper-resistant (inaccessible) loci, while positive scores reflect DNS hyper-sensitive (accessible) loci. The bedmap’s “– sum” and “—mean” were used to process DNS scores from genomic informative intervals, *i.e*. whole gene models, 500 bp upstream of CDS (positions ranging from −500 to −1 of start of annotated HC gene models), 2 kb upstream of CDS (positions ranging from −2000 to −1 of start of HC gene models), 2kb downstream of end of CDS (positions ranging from (+1 to 2000 from end of HC gene models), intergenic space (positions ranging more than 2 kb from end of HC gene model, and more than - 2 kb from the start of the adjacent HC gene models), annotated TE space [34], 1 Mb and 2 Mb windows across entire genome. To make DNS values comparable across regions, all DNS values presented in this paper represent the average DNS score for 10 bp intervals within each informative genomic region.

### MNase Hypersensitive (MSF) and Hyper-resistant (MRF) Regions

We performed a segmentation analysis using the iSeg algorithm [36] to identify distinctly accessible (hypersensitive) and inaccessible (hyper-resistant) regions of the genome. A biological cutoff for genome-wide significance of sd = 1.5, was used to identify regions either accessible or inaccessible to MNase digest. Replicates were run separately and regions that were found to surpass the biological cutoff in both replicates were considered either accessible (hypersensitive, MSF), or inaccessible (hyper-resistant, MRF). These MSF and MRF regions were mapped in relation to genomic features using the closest features tool from the BedOps suite [65] to examine their relative distribution within the genome and their proximity to genic space and TE regions. Segmentation analysis scores are highly correlated with DNS values (Figure S1c-f in Additional File 1).

### Gene expression analysis

A subset of gene expression values for cultivar Chinese Spring was selected from the wheat genome expression database [30]. We selected 5 replications of non-stressed CS leaves and shoots 14 days old from the recent meta-analysis [30] to match our tissue type and age. Gene expression for high confidence (HC) genes was calculated as the average expression across 5 biological replicates from the study. Genes were considered expressed if mean expression was ≥ 0.1 tpm (73,437 HC genes). Gene expression values were log_10_-transformed and correlated with DNS score for certain genic regions (2 kb upstream, 500 bp upstream, gene body, 2 kb downstream, intergenic space). Recently, the transcriptional landscape of wheat was released, which discussed partitioning of 1:1:1 triplets into seven categories based on relative expression contribution. We grouped the syntenic triplets into these categories using same criteria as previously described using the gene expression data from 5 reps of Chinese Spring expression data [30]. Only syntenic triplets that had a sum of ≥ 0.5 tpm were used in this analysis, leaving 12,601 total triplet sets for analysis (Table S6 in Additional File 2).

### Transposable Element Enrichment

Coordinates of various TE superfamily/family content within the CS genome were obtained from the recently released version of the wheat genome [35]. We associated patterns of DNS scores and iSeg densities with TE family content and frequency across the genome. Spearman correlation test was used to test the correlation between the proportion of Gypsy TEs and DNS score for 1-Mb sliding windows with 200 kb step. Only those windows that contained each type of TEs were used in analysis.

### Effect of chromatin accessibility on genetic variance

The previously published phenotypic data described in our study by He et al. (2019) was used for variance partitioning [47]. The Best Linear Unbiased Estimates were obtained by fitting a model with fixed genotype effects and all other effects as random in an individual year. The trait values from the rainfed and irrigated (I) trials were used to calculate the stress susceptibility index [66]. For each trait, the year and environment were added as a suffix to the trait name. The following traits were included into the analyses: days to heading in 2014 (HD14_I); days to heading in 2015 (HD15_I); plant height in 2014 (PHT14_I); plant height in 2015 (PHT15_I); grain filling period in 2014 (GFP14_I); grain filling period in 2015 (GFP15_I); harvest weight of grain in 2014 (HW14_I); harvest weight of grain in 2015 (HW15_I); stress susceptibility index for harvest weight in 2014 (HW14_S) and 2015 (HW15_S).

Using the DNS scores calculated for 10-bp long intervals across genome, we ranked the entire genome from the most closed to the most open chromatin intervals based on the DNS score distribution. Intervals were split into 5 groups, each representing 20% of the genome based on accessible chromatin score. SNPs extracted from the 1000 wheat exomes project [47] were filtered to retain one SNP every 10 kb with MAF > 0.002 resulting in a total of 239,000 variable sites. SNPs that fell within the gene bodies and within 1 kb flanking regions of genes were extracted, and grouped into 5 bins of different DNS score distributions. A total of 10,000 SNPs were randomly selected from the most closed (0-20% bin) and most open (80-100% bin) bins of the genome, and the proportion of phenotypic variance explained by these two groups of SNPs was estimated using the GCTA-GREML method, as previously described [47, 67]. The variance partitioning with these randomly selected SNP sets was repeated 50 times, and the proportions of phenotypic variance for each trait, V(G)/V(p), were extracted from each calculation.

## Abbreviations

DNS: differential nuclease sensitivity
MNase: micrococcal nuclease
TE: transposable element
CENH3: centromeric variant of H3 histone

## Declarations

### Ethics approval and consent to participate

Not applicable.

### Consent for publication

Not applicable.

### Availability of data and materials

Raw sequence data is available for download from the NCBI BioProject PRJNA564769. The DNS score and read coverage tracks are made available through the Triticeae Toolbox database (https://triticeaetoolbox.org).

### Authors’ contributions

KJ isolated wheat chromatin, performed MNase treatment, analyzed chromosomal patterns of chromatin accessibility distribution across the genome and its impact on gene expression, and wrote the first draft of the manuscript. FH processed raw data to generate chromatin accessibility scores, analyzed locations of centromeres relative to chromatin states and TE density and partitioned genetic variance. MF and AA prepared DNS-seq libraries, evaluated their quality and generated NGS data. EA proposed the idea, coordinated data analyses, interpreted results, and wrote the manuscript. All authors read and approved the final manuscript.

## Acknowledgements

We would like to thank the KSU Integrated Genomics Facility and KU Medical Center’s Genome Sequencing Facility for help with performing next-generation sequencing, D. Andresen for assistance with the computing resources of the KSU Beocat cluster funded by NSF CHE-1726332 and NIH P20GM113109 grants.

## Additional File 1

File format: PDF

Table S1. Alignment Statistics for DNS-Seq

Table S2. Genome and Chromosome DNS Scores and Proportion of MSF/MRF regions

Table S3. Pericentromeric-Distal Comparisons of DNS Scores and Relative Fold Changes

Table S4. Homoeologous Chromosome Group 4 DNS Segment Comparison

Table S5. MSF and MRF Region Annotation

Table S6. Chromatin States DNS Scores and Outlier Overlap

Table S8. Comparison of DNS Scores for Triplets Around Genes

Table S9. TE Superfamily Correlations of DNS Score and TE Density

Table S11. Intergenic Distance Effect on DNS Score

Table S12. Centromere Mapping with DNS Score, Cereba Density, and Read Depth

Table S14. Phenotypic Variance by Region Summary

Figure S1. Correlation of DNS Scores and MRF/MSF outliers

Figure S2. Recombination Rate and DNS Score Correlation

Figure S3. DNS Score and Proportion of MRF/MSF regions for Homoeologous Chromosomes 1

Figure S4. DNS Score and Proportion of MRF/MSF regions for Homoeologous Chromosomes 2

Figure S5. DNS Score and Proportion of MRF/MSF regions for Homoeologous Chromosomes 3

Figure S6. DNS Score and Proportion of MRF/MSF regions for Homoeologous Chromosomes 4

Figure S7. DNS Score and Proportion of MRF/MSF regions for Homoeologous Chromosomes 5

Figure S8. DNS Score and Proportion of MRF/MSF regions for Homoeologous Chromosomes 6

Figure S9. DNS Score and Proportion of MRF/MSF regions for Homoeologous Chromosomes 7

Figure S10. MRF/MSF Outlier Region Descriptions Annotation

Figure S11. DNS Scores Around Genes by Genome

Figure S12. Categorized Syntenic Triplet Expression Contribution

Figure S13. DNS Scores for Common TE Superfamilies

Figure S14. DNS Scores by Family of Common TE Superfamilies

Figure S15. TE DNS Scores Relative to Gene Proximity

Figure S16. Intergenic Distance Distribution for Distal and Centromeric Regions

Figure S17. Sensitivity of Centromeric Chromatin to Differential MNase Digest

## Additional File 2

File format: TXT

Table S7. Triplets, Designation Category, and Expression (separate txt file)

## Additional File 3

File format: TXT

Table S10. TE Family DNS Mean Scores (separate excel file)

## Additional File 4

File format: TXT

Table S13. Phenotypic Variance by Decile Raw Data (separate excel file)

