## Additional File 1 for "Differential chromatin accessibility landscape reveals the structural and functional features of the allopolyploid wheat chromosomes"

Table S1. Alignment Statistics for DNS-Seq

Table S2. Genome and Chromosome DNS Scores and Proportion of MSF/MRF regions

Table S3. Proximal-Distal Comparisons of DNS Scores and Relative Fold Changes

Table S4. Homoeologous Chromosome Group 4 DNS Segment Comparison

Table S5. MSF and MRF Region Annotation

Table S6. Chromatin States DNS Scores and Outlier Overlap

Table S8. Comparison of DNS Scores for Triplets Around Genes

Table S9. TE Superfamily Correlations of DNS Score and TE Density

Table S11. Intergenic Distance Effect on DNS Score

Table S12. Centromere Mapping with DNS Score, Cereba Density, and Read Depth

Table S14. Phenotypic Variance by Region Summary

Figure S1. Correlation of DNS Scores and MRF/MSF outliers

Figure S2. Recombination Rate and DNS Score Correlation

Figure S3. DNS Score and Proportion of MRF/MSF regions for Homoeologous Chromosomes 1

Figure S4. DNS Score and Proportion of MRF/MSF regions for Homoeologous Chromosomes 2

Figure S5. DNS Score and Proportion of MRF/MSF regions for Homoeologous Chromosomes 3

Figure S6. DNS Score and Proportion of MRF/MSF regions for Homoeologous Chromosomes 4

Figure S7. DNS Score and Proportion of MRF/MSF regions for Homoeologous Chromosomes 5

Figure S8. DNS Score and Proportion of MRF/MSF regions for Homoeologous Chromosomes 6

Figure S9. DNS Score and Proportion of MRF/MSF regions for Homoeologous Chromosomes 7

Figure S10. MRF/MSF Outlier Region Descriptions Annotation

Figure S11. Categorized Syntenic Triplet Expression Contribution

Figure S12. DNS Scores Around Genes by Genome

Figure S13. DNS Scores for Common TE Superfamilies

Figure S14. DNS Scores by Family of Common TE Superfamilies

Figure S15. TE DNS Scores Relative to Gene Proximity

Figure S16. Intergenic Distance Distribution for Distal and Centromeric Regions

Figure S17. Sensitivity of Centromeric Chromatin to Differential MNase Digest

### Table S1. Alignment Statistics for MNase-Seq

| Library | Total Reads (PE) | QC Passed Reads (PE) | Aligned (total reads) | Uniquely Aligned (total reads) |
|---|---|---|---|---|
| Light Digest Rep1 | 416,522,545 | 402,865,390 | 780,813,754 | 589,984,079 (71%) |
| Heavy Digest Rep 1 | 434,239,251 | 417,622,726 | 782,477,672 | 589,871,191 (68%) |
| Light Digest Rep2 | 429,145,753 | 414,587,048 | 797,782,573 | 577,209,203 (67%) |
| Heavy Digest Rep2 | 469,915,480 | 444,680,355 | 862,303,550 | 632,946,226 (67%) |
| **Average per library** | 437,455,757 | 419,938,880 | 805,844,387 | 597,502,675 |
| **Totals** | 1,749,823,029 | 1,679,755,519 | 3,223,377,549 | 2,390,010,699 (68%) |

**Table S2. Genome and Chromosome DNS Scores and Proportion MSF/MRF regions**

| Comparison | Region | Length (Mb) | MSF (Mb) | Proportion MSF | MRF (Mb) | Proportion MRF | DNS score† |
|---|---|---|---|---|---|---|---|
| Genomes | A | 4930 | 59.38 | 0.01205 | 73.30 | 0.01487 | 0.0032 |
| | B | 5176 | 67.22 | 0.01299 | 86.33 | 0.01668 | -0.0016 |
| | D | 3946 | 50.66 | 0.01284 | 55.17 | 0.01398 | 0.009 |
| | Whole Genome | 14052 | 177.26 | 0.01261 | 214.80 | 0.01529 | 0.0031 |
| Chromosomes | chr1A | 594 | 7.18 | 0.01209 | 8.59 | 0.01446 | 0.0061 |
| | chr1B | 689 | 8.99 | 0.01305 | 11.32 | 0.01644 | 0.0000 |
| | chr1D | 495 | 6.49 | 0.01310 | 6.58 | 0.01329 | 0.0131 |
| | chr2A | 780 | 8.98 | 0.01151 | 11.83 | 0.01517 | 0.0000 |
| | chr2B | 801 | 10.81 | 0.01349 | 13.05 | 0.01629 | 0.0026 |
| | chr2D | 651 | 8.22 | 0.01263 | 9.17 | 0.01408 | 0.0073 |
| | chr3A | 750 | 8.82 | 0.01177 | 11.23 | 0.01497 | 0.0018 |
| | chr3B | 830 | 10.89 | 0.01313 | 14.09 | 0.01698 | -0.0028 |
| | chr3D | 615 | 7.74 | 0.01259 | 8.65 | 0.01407 | 0.0074 |
| | chr4A | 744 | 9.13 | 0.01228 | 11.74 | 0.01578 | -0.0008 |
| | chr4B | 673 | 7.79 | 0.01157 | 12.28 | 0.01824 | -0.0117 |
| | chr4D | 509 | 5.98 | 0.01175 | 8.07 | 0.01585 | -0.0004 |
| | chr5A | 709 | 8.63 | 0.01217 | 10.05 | 0.01417 | 0.0067 |
| | chr5B | 713 | 9.77 | 0.01370 | 11.00 | 0.01542 | 0.0051 |
| | chr5D | 565 | 7.55 | 0.01336 | 7.33 | 0.01297 | 0.0150 |
| | chr6A | 617 | 7.07 | 0.01146 | 9.28 | 0.01503 | 0.0005 |
| | chr6B | 720 | 9.14 | 0.01270 | 12.21 | 0.01696 | -0.0046 |
| | chr6D | 473 | 6.08 | 0.01285 | 6.64 | 0.01404 | 0.0088 |
| | chr7A | 736 | 9.56 | 0.01299 | 10.59 | 0.01438 | 0.0084 |
| | chr7B | 750 | 9.82 | 0.01309 | 12.38 | 0.01651 | -0.0004 |
| | chr7D | 638 | 8.60 | 0.01348 | 8.73 | 0.01369 | 0.0114 |

† DNS mean of 2Mb windows across each genome and chromosome

**Table S3. Proximal-Distal Comparisons of DNS Scores and Relative Fold Changes**

| Comparison | Genome | overall mean† | R1 mean‡ (FC) | C mean ‡ (FC) | R3 mean‡ (FC) |
|---|---|---|---|---|---|
| 2Mb windows | A | 0.0032 | 0.0423 (13.2) | -0.0154 (-4.8) | 0.047 (14.7) |
| | B | -0.0016 | 0.0468 (29.3) | -0.0133 (-8.3) | 0.0368 (23.0) |
| | D | 0.009 | 0.054 (6.0) | -0.0117 (-1.3) | 0.0451 (5.0) |
| Gene Body | A | 0.0929 | 0.0887 (1.0) | 0.135 (1.5) | 0.0853 (0.9) |
| | B | 0.0959 | 0.0941 (1.0) | 0.152 (1.6) | 0.0882 (0.9) |
| | D | 0.1002 | 0.0896 (0.9) | 0.152 (1.5) | 0.0912 (0.9) |
| 500bp Upstream | A | 0.256 | 0.259 (1.0) | 0.244 (1.0) | 0.257 (1.0) |
| | B | 0.259 | 0.26 (1.0) | 0.266 (1.0) | 0.256 (1.0) |
| | D | 0.268 | 0.268 (1.0) | 0.267 (1.0) | 0.269 (1.0) |
| 2kb Upstream | A | 0.18 | 0.187 (1.0) | 0.16 (0.9) | 0.19 (1.1) |
| | B | 0.175 | 0.189 (1.1) | 0.168 (1.0) | 0.183 (1.0) |
| | D | 0.184 | 0.192 (1.0) | 0.176 (1.0) | 0.192 (1.0) |
| 2kb Downstream | A | 0.161 | 0.173 (1.1) | 0.148 (0.9) | 0.17 (1.1) |
| | B | 0.158 | 0.175 (1.1) | 0.158 (1.0) | 0.171 (1.1) |
| | D | 0.167 | 0.183 (1.1) | 0.168 (1.0) | 0.176 (1.1) |
| Intergenic | A | 0.00719 | 0.0704 (9.8) | -0.00423 (-0.6) | 0.068 (9.5) |
| | B | 0.00498 | 0.0675 (13.6) | 0.00157 (0.3) | 0.0523 (10.5) |
| | D | 0.00833 | 0.0763 (9.2) | 0.0048 (0.6) | 0.0639 (7.7) |
| all TE | A | 0.00594 | 0.0647 (10.9) | 0.0029 (0.5) | 0.0657 (11.1) |
| | B | 0.00292 | 0.063 (21.6) | -0.0123 (-4.2) | 0.0472 (16.2) |
| | D | 0.00612 | 0.0648 (10.6) | 0.00266 (0.4) | 0.0578 (9.4) |
| Gypsy | A | -0.00447 | 0.0266 (6.0) | -0.0147 (-3.3) | 0.0281 (6.3) |
| | B | -0.0132 | 0.032 (2.4) | -0.0173 (-1.3) | 0.0193 (1.5) |
| | D | -0.00381 | 0.0324 (8.5) | -0.0179 (-4.7) | 0.0275 (7.2) |
| Copia | A | 0.00848 | 0.0283 (3.3) | -0.00079 -(0.1) | 0.0328 (3.9) |
| | B | 0.0057 | 0.033 (5.8) | 0.00034 (0.1) | 0.032 (5.6) |
| | D | 0.0146 | 0.0371 (2.5) | 0.0084 (0.6) | 0.0338 (2.3) |
| CACTA | A | 0.0347 | 0.0648 (1.9) | 0.0248 (0.7) | 0.0704 (2.0) |
| | B | 0.0248 | 0.0603 (2.4) | 0.0119 (0.5) | 0.0506 (2.0) |
| | D | 0.0229 | 0.0547 (2.4) | 0.0154 (0.7) | 0.0514 (2.2) |

† DNS mean for whole genome specified in each comparison
‡ DNS mean for each comparison only in the particular chromosomal segment
(FC) fold change with respect to overall mean
Intergenic regions are segments of the genome more than 2kb away from HC genes

**Table S4. Homoeologous Chromosome Group 4 DNS Segment Comparison**

| Comparison | Region | A genome† | Chr4A‡ | Change§ | B genome† | Chr4B‡ | Change § | D genome† | Chr4D‡ | Change § |
|---|---|---|---|---|---|---|---|---|---|---|
| Segment | R1 | 0.0423 | 0.0119 | 0.2813 | 0.0468 | 0.0292 | 0.6239 | 0.054 | 0.0684 | 1.2667 |
| | R2a | -0.0081 | -0.0112 | 1.3827 | -0.0152 | -0.0217 | 1.4276 | 0.00209 | 0.00914 | 4.3732 |
| | C | -0.01545 | -0.0133 | 0.8608 | -0.0133 | -0.0248 | 1.8647 | -0.0117 | -0.022 | 1.8803 |
| | R2b | -0.00862 | -0.0148 | 1.7169 | -0.015 | -0.0196 | 1.3067 | -0.00433 | -0.00681 | 1.5727 |
| | R3 | 0.043 | 0.0576 | 1.3395 | 0.0335 | 0.013 | 0.3881 | 0.0433 | 0.029 | 0.6697 |
| Whole Chromosome | | 0.00273 | -0.00076 | -0.2784 | -0.00158 | -0.0117 | 7.4051 | 0.009 | -0.000419 | -0.0466 |

† mean of 2Mb windows for each segment for whole genome
‡ mean of 2Mb windows for homeologous group 4 chromosomes
§ Relative change for group 4 to overall genome mean

**Table S5**. MSF and MRF Region Annotation ((The International Wheat Genome Sequencing Consortium (IWGSC) 2018)

| **Designation** | **MSF (Hyper-sensitive Footprints)** | | | | **MRF (Hyper-resistant Footprints)** | | | | **Enrichment (FET) ‡** |
|---|---|---|---|---|---|---|---|---|---|
| **Location** | Total | A | B | D | Total | A | B | D | Overall |
| **Total** | 2,156,684 | 731,872 | 818,940 | 605,872 | 2,605,884 | 890,919 | 1,038,272 | 676,693 | |
| **Genic HC†** | 368,071 | 123,751 | 125,613 | 118,707 | 48,143 | 16,349 | 17,813 | 13,981 | MSF |
| **Genic LC†** | 95,896 | 9,859 | 11,661 | 9,177 | 38,180 | 12,053 | 15,612 | 10,515 | MSF |
| **All TE** | 1,440,502 | 496,600 | 565,201 | 378,701 | 2,360,569 | 811,588 | 942,999 | 605,982 | MRF |
| **Class 1 TE** | 1,064,531 | 380,996 | 415,498 | 268,037 | 1,992,012 | 698,840 | 793,881 | 499,291 | MRF |
| **Gypsy (RLG)** | 752,110 | 274,887 | 292,029 | 185,194 | 1,509,315 | 531,480 | 602,806 | 375,029 | MRF |
| **Copia (RLC)** | 225,201 | 79,602 | 87,828 | 57,771 | 333,760 | 121,519 | 130,383 | 81,858 | MSF |
| **Class 2 TE** | 346,059 | 106,278 | 137,264 | 102,517 | 341,587 | 104,477 | 137,235 | 99,875 | MSF |
| **CACTA (DTC)** | 271,957 | 82,639 | 108,948 | 80,370 | 301,012 | 92,630 | 120,054 | 88,328 | MSF |
| **Mutator (DTM)** | 14,305 | 4,391 | 5,531 | 4,383 | 8,792 | 2,533 | 3,223 | 3,036 | MSF |
| **Mariner (DTT)** | 29,201 | 8,890 | 11,731 | 8,580 | 12,313 | 3,216 | 5,461 | 3,636 | MSF |
| **Harbinger (DTH)** | 7,287 | 2,488 | 2,578 | 2,221 | 5,221 | 1,628 | 2,084 | 1,509 | MSF |
| **Unclassified TE** | 29,912 | 9,326 | 12,439 | 8,147 | 26,970 | 8,271 | 11,883 | 6,816 | MSF |
| **Not annotated** | 252,215 | 101,662 | 116,465 | 99,287 | 158,992 | 50,929 | 61,848 | 46,215 | |

† Includes gene body and within 2 kb up- or downstream of gene body

‡ Fisher Exact Test enrichment comparing total numbers of MSF and MRF regions (*p-values* $< 10^{-16}$)

**Table S6.** Chromatin States DNS Scores and Outlier Overlap

| Chromatin State† | DNS Mean | DNS Std Dev | MSF Overlap ‡ (bp) | % MSF § | MRF Overlap ‡ (bp) | % MRF § |
|---|---|---|---|---|---|---|
| s1 | 0.293 | 0.144 | 7,891,543 | 4.45 | 67,818 | 0.03 |
| s2 | 0.246 | 0.150 | 3,064,078 | 1.73 | 28,057 | 0.01 |
| s3 | 0.335 | 0.170 | 3,849,585 | 2.17 | 31,231 | 0.01 |
| s4 | 0.307 | 0.194 | 5,704,560 | 3.22 | 74,358 | 0.03 |
| s5 | 0.094 | 0.166 | 1,847,416 | 1.04 | 105,585 | 0.05 |
| s6 | 0.179 | 0.190 | 3,400,949 | 1.92 | 248,310 | 0.12 |
| s7 | 0.250 | 0.162 | 4,353,181 | 2.46 | 116,610 | 0.05 |
| s8 | 0.195 | 0.208 | 4,210,225 | 2.38 | 255,387 | 0.12 |
| s9 | 0.093 | 0.167 | 5,403,994 | 3.05 | 1,135,641 | 0.53 |
| s10 | 0.119 | 0.175 | 17,845,989 | 10.07 | 3,540,057 | 1.65 |
| s11 | 0.178 | 0.206 | 6,044,327 | 3.41 | 559,136 | 0.26 |
| s12 | 0.077 | 0.177 | 7,685,771 | 4.34 | 3,409,378 | 1.59 |
| s13 | -0.188 | 0.176 | 3,126,621 | 1.76 | 69,152,473 | 32.19 |
| s14 | -0.444 | 2.757 | 203,072 | 0.11 | 747,055 | 0.35 |
| s15 | -4.405 | 15.535 | 668,356 | 0.38 | 746,968 | 0.35 |

† Previously defined in Li et al. [18] using histone acetylation and methylation epigenetic marks

‡ Amount of overlap (bp) of bed files defining chromatin states [18], and MSF and MRF outliers using DNS-seq

§ The percent of overlapped features in terms of the total size of MSF and MRF outliers

**Table S7. Triplets, designation category, and expression**. Separate text file (on server).
/home/kjordan/Chromatin/Revised_Manuscript /TableS7.Triplets_category_contribution.txt

**Table S8. Comparisons of DNS scores for Triplets Around Genes**

| Triplet Designation | Region* | Triplet Category | Test-statistic § | p-value† | A-B test (p-value)‡ | A-D test (p-value)‡ | B-D test (p-value)‡ |
|---|---|---|---|---|---|---|---|
| Balanced | *a* | | 0.337 | 0.84 | 0.6299 | 0.6063 | 0.984 |
| | *b* | | 1.67 | 0.43 | 0.2398 | 0.3092 | 0.7907 |
| | *c* | | 2.69 | 0.26 | 0.1392 | 0.9972 | 0.1777 |
| | *d* | | 0.236 | 0.89 | 0.6347 | 0.7228 | 1 |
| Suppression | *a* | Asup | 18.87 | 7.98E-05 | 3.53E-05 | 0.001658 | 0.2812 |
| | | Bsup | 17.06 | 0.0001505 | 0.0006578 | 0.8619 | 0.0001291 |
| | | Dsup | 4.13 | 0.1269 | 0.4575 | 0.02815 | 0.3522 |
| | *b* | Asup | 17.22 | 0.0001821 | 0.0001358 | 0.002337 | 0.1566 |
| | | Bsup | 18.29 | 0.0001068 | 0.0003075 | 0.6616 | 0.0001649 |
| | | Dsup | 1.71 | 0.426 | 0.7695 | 0.2211 | 0.3293 |
| | *c* | Asup | 7.42 | 0.02453 | 0.01822 | 0.01926 | 0.9698 |
| | | Bsup | 4.2315 | 0.1205 | 0.08419 | 0.8876 | 0.06929 |
| | | Dsup | 0.822 | 0.663 | 0.361 | 0.5932 | 0.7538 |
| | *d* | Asup | 11.07 | 0.003955 | 0.001816 | 0.3352 | 0.0154 |
| | | Bsup | 15.65 | 0.0003989 | 0.0002789 | 0.2283 | 0.003028 |
| | | Dsup | 8.85 | 0.01199 | 0.7992 | 0.03222 | 0.002775 |
| Dominant | *a* | Adom | 43.37 | 3.82E-10 | 7.36E-09 | 2.27E-05 | 9.55E-05 |
| | | Bdom | 7.31 | 0.02583 | 0.01141 | 0.3843 | 0.05227 |
| | | Ddom | 22.24 | 1.48E-05 | 4.27E-06 | 0.1502 | 0.001583 |
| | *b* | Adom | 19.78 | 5.10E-05 | 1.57E-05 | 0.01295 | 0.03015 |
| | | Bdom | 23.01 | 1.01E-05 | 0.1045 | 0.001611 | 3.11E-06 |
| | | Ddom | 48.35 | 3.17E-11 | 0.6221 | 9.06E-10 | 4.01E-09 |
| | *c* | Adom | 34.45 | 3.30E-08 | 5.05E-08 | 0.01691 | 5.27E-05 |
| | | Bdom | 29.15 | 4.68E-07 | 4.19E-06 | 0.1868 | 4.19E-06 |
| | | Ddom | 53.32 | 2.65E-12 | 0.2539 | 1.34E-09 | 7.46E-11 |
| | *d* | Adom | 18.3 | 0.0001062 | 0.0002036 | 0.0002385 | 0.8935 |
| | | Bdom | 1.27 | 5.30E-01 | 0.5205 | 0.1965 | 0.8724 |
| | | Ddom | 12.3 | 0.002133 | 0.1579 | 0.002899 | 0.00464 |

*Region *a*: 1kb to 500bp upstream; *b*: 500bp to CDS start; *c*: CDS start to 500bp downstream; *d*: 500bp -1kb downstream

§ Kruskal Wallis test statistic (chi-square)

† Kruskal Wallis p-value

‡ Wilcoxon test p-value between 2 genomes

**Table S9. TE Superfamily Correlations of DNS Scores and TE Density**

| Type | Family | A cor † | A Windows ‡ | B cor † | B Windows ‡ | D cor † | D Windows ‡ | Genome Composition § |
|---|---|---|---|---|---|---|---|---|
| Class 1 | RLG- Gypsy | -0.6768 | 24677 | -0.6368 | 25904 | -0.6734 | 19759 | 46.7 |
| | RLC -Copia | 0.1338 | 24675 | 0.0444 | 25904 | 0.148 | 19758 | 16.7 |
| | RLX -Unclassified LTR | -0.2243 | 24528 | -0.2163 | 25851 | -0.321 | 19720 | 3.2 |
| | RIX -Line | 0.2727 | 23537 | 0.2207 | 25252 | 0.1967 | 19297 | 0.9 |
| | SIX -Sine | 0.0919 | 8718 | 0.034 | 12546 | 0.0436 | 10606 | 0.01 |
| Class 2 | DTC -CACTA | -0.2579 | 24676 | -0.1317 | 25901 | -0.295 | 19750 | 15.5 |
| | DTM -Mutator | 0.1747 | 23649 | 0.1361 | 25261 | 0.1484 | 19508 | 0.38 |
| | DTX -Unclassified terminal repeats | 0.2682 | 24033 | 0.2114 | 25398 | 0.1815 | 19558 | 0.21 |
| | DTH -Harbinger | 0.357 | 21282 | 0.2448 | 23141 | 0.2572 | 18289 | 0.16 |
| | DTT -Mariner | 0.0193 | 24487 | 0.0006 | 25794 | -0.0609 | 19711 | 0.16 |
| | DXX -Unclassiried Type2 | -0.3836 | 9116 | -0.2178 | 12010 | -0.3345 | 8593 | 0.06 |
| | DTA- hAT | -0.1758 | 1750 | -0.1149 | 2064 | -0.1783 | 1296 | 0.01 |
| Unclassified | XXX -Unclassified Repeats | 0.3822 | 24409 | 0.2921 | 25820 | 0.4099 | 19672 | 0.68 |

† Spearman correlation of proportion of TE content per 1MB window and DNS score of same window
‡ Windows with TE present are used for correlation
§ Whole genome percentage of TE type (IWGSC, 2018)

**TableS10. TE family DNS Mean Scores (separate excel file on server)**
/home/kjordan/Chromatin/Revised_Manuscript TableS10.DNSmeans_subfamilies_formatted.xlsx

**Table S11. Intergenic Distance Effect on DNS Score**

| Category | Comparison | DNS Mean† | DNS SD† | P-Value D vs C‡ |
|---|---|---|---|---|
| Intergenic Distance < 10kb | all regions | 0.148 | 0.064 | 0.4 |
| | Distal (R1 and R3) | 0.162 | 0.053 | |
| | Centromeric (C) | 0.146 | 0.06 | |
| 10kb-100kb | all regions | 0.0279 | 0.0325 | 3.68 x 10-12 |
| | Distal (R1 and R3) | 0.0378 | 0.013 | |
| | Centromeric (C) | 0.00629 | 0.0168 | |
| 100kb-1Mb | all regions | -0.0172 | 0.0198 | 2.2 x 10-16 |
| | Distal (R1 and R3) | 0.0069 | 0.028 | |
| | Centromeric (C) | -0.0168 | 0.026 | |
| Intergenic Distance >1Mb | all regions | -0.178 | 0.0175 | 1.19 x 10-6 |
| | Distal (R1 and R3) | 0.0045 | 0.027 | |
| | Centromeric (C) | -0.0159 | 0.015 | |

† Mean and standard deviation of DNS score of all 1kb windows in each distance category
‡ Wilcoxon test statistic comparing distribution of DNS scores for randomly matched sample size

**Table S12. Centromere Mapping with DNS Score, Cereba Density, and Read Depth**

| Chromosome | Centromere Position† | Heavy Depth‡ | Light Depth‡ | Read Depth Peak§ | Proportion Cereba* | Cereba Peak§ | DNS Score* | DNS Peak§ | DNS/CENH3 Consistency |
|---|---|---|---|---|---|---|---|---|---|
| chr1A | 210.2 - 215.8 | 0.8154 | 0.9587 | 214.0 | 0.5018 | 214.6 | 0.0830 | 213.0 | yes |
| chr1B | 237.7 - 243.5 | 0.7760 | 0.8062 | 244.0 | 0.5357 | 244.0 | 0.0856 | 240.0 | yes |
| chr1D | 166.2 - 173.8 | 1.6257 | 1.7586 | 173.2 | 0.1301 | 173.0 | 0.0944 | 170.4 | yes |
| chr2A | 326.3 - 327 | 0.9175 | 1.0723 | 326.4 | 0.5280 | 327.8 | 0.0639 | 326.4 | yes |
| | 339.4 - 342 | 0.9228 | 1.0253 | 339.4 | 0.4055 | 339.4 | 0.0417 | 340.6 | yes |
| | 359.3 - 359.5 | 2.8237 | 2.6973 | 359.4 | 0.0849 | 357.4 | 0.0415 | 356.4 | yes |
| chr2B | 344.4 - 351.3 | 0.8363 | 1.0396 | 346.8 | 0.4447 | 347.8 | 0.0945 | 347.6 | yes |
| chr2D | 264.4 - 272.5 | 1.4505 | 1.7621 | 269.4 | 0.3588 | 269.0 | 0.1179 | 270.8 | yes |
| chr3A | 316.9 - 319.9 | 0.5680 | 0.7032 | 318.8 | 0.5255 | 318.0 | 0.0696 | 318.2 | yes |
| chr3B | 345.8 - 347 | 1.7936 | 1.9935 | 348.0 | 0.2524 | 348.2 | 0.0615 | 348.4 | yes |
| | 348.5 - 349 | 1.7936 | 1.9935 | 348.0 | 0.2524 | 348.2 | 0.0615 | 348.4 | yes |
| chr3D | 237.1 - 243.2 | 1.0633 | 1.2509 | 242.2 | 0.3157 | 240.6 | 0.0939 | 240.8 | yes |
| chr4A | 264.1 - 267.9 | 0.5172 | 0.6945 | 264.4 | 0.5767 | 264.6 | 0.0962 | 264.2 | yes |
| | 315.1 - 315.7 | 1.3570 | 1.4808 | 314.8 | 0.3083 | 314.8 | 0.0443 | 314.6 | yes |
| chr4B | 303.9 - 304.4 | 1.8836 | 1.8716 | 303.4 | 0.2988 | 303.8 | 0.0396 | 299.4 | close - within 4Mb |
| | 317.8 - 319.6 | 0.8784 | 0.9664 | 318.8 | 0.3989 | 319.0 | 0.0483 | 316.6 | yes |
| chr4D | 182.3 - 188.2 | 1.8283 | 1.8047 | 209.4 | 0.1388 | 209.4 | 0.1094 | 185.2 | yes DNS - not Cereba |
| chr5A | 108.9 - 109.1 | 2.0596 | 2.1262 | 109.0 | 0.0667 | 108.2 | 0.0268 | 104.0 | Region R2b |
| | 248.7 - 249 | 1.0245 | 1.1053 | 248.4 | 0.4413 | 248.6 | 0.0657 | 248.2 | yes |
| | 252.5 - 255.1 | 0.7941 | 1.0226 | 252.8 | 0.5566 | 253.0 | 0.0893 | 252.8 | yes |
| chr5B | 198.9 - 202.5 | 1.8467 | 1.9865 | 200.4 | 0.2238 | 201.6 | 0.0556 | 196.2 | yes |
| chr5D | 185.6 - 188.7 | 1.2719 | 1.5883 | 186.0 | 0.2817 | 185.6 | 0.0930 | 186.2 | yes |
| | | 1.0841 | 1.4133 | 146.2 | 0.4953 | 145.0 | 0.1086 | 146.2 | not annotated |
| chr6A | 283.3 - 288.7 | 1.0025 | 1.1268 | 285.0 | 0.2947 | 289.2 | 0.0935 | 284.4 | yes |
| | 290.7 - 292.5 | 0.9300 | 1.2746 | 287.8 | 0.2947 | 289.2 | 0.0817 | 287.6 | close - within 3Mb |
| chr6B | 323 - 327.5 | 0.7531 | 0.9624 | 325.4 | 0.4673 | 325.2 | 0.0921 | 326.0 | yes |
| chr6D | 211.9 - 217.4 | 1.6014 | 1.9367 | 213.4 | 0.2654 | 213.2 | 0.0869 | 213.4 | yes |
| chr7A | 360.2 - 363.8 | 1.1393 | 1.3561 | 361.2 | 0.2804 | 361.4 | 0.0929 | 361.8 | yes |
| chr7B | 288.2 - 288.3 | 2.6389 | 2.6562 | 288.2 | 0.0912 | 288.0 | 0.0398 | 289.8 | yes |
| | 294.4 - 294.6 | 1.6792 | 1.7328 | 293.6 | 0.3789 | 293.8 | 0.0504 | 292.2 | yes |
| | 296.4 - 296.5 | 2.1153 | 2.2123 | 296.4 | 0.1649 | 296.4 | 0.0346 | 296.2 | yes |

| | | | | | | | | | |
|---|---|---|---|---|---|---|---|---|---|
| | 308 - 310.1 | 0.9300 | 1.0972 | 308.2 | 0.5207 | 308.2 | 0.0702 | 308.0 | yes |
| chr7D | 336.3 - 341.7 | 1.3632 | 1.6799 | 338.0 | 0.3118 | 337.0 | 0.0951 | 339.0 | yes |

† Position of CENH3 from IWGSC, 2018
‡ Minimum Depth of Coverage for 1Mb windows in region
§ 1Mb window position where read depth is minimal, Cereba proportion, or DNS value is maximal
* Peak value for Cerebra proportion and DNS score for 1Mb window

**Table S13. Phenotypic Variance by Decile Raw Data**. Separate excel file.

**Table S14. Phenotypic Variance by Region Summary**.

| **Phenotype** | **Open Regions†** | | **Closed Regions‡** | | ***p-value*§** |
|---|---|---|---|---|---|
| | **Mean** | **SD** | **Mean** | **SD** | |
| HW14_S | 0.4388 | 0.1986 | 0.1806 | 0.1863 | 2.80E-08 |
| HW15_S | 0.4572 | 0.1230 | 0.0249 | 0.0757 | 2.00E-16 |
| HD14_I | 0.6723 | 0.1764 | 0.2119 | 0.1695 | 2.40E-15 |
| HD15_I | 0.5710 | 0.1837 | 0.2889 | 0.1920 | 6.40E-10 |
| PHT14_I | 0.7123 | 0.1602 | 0.1282 | 0.1416 | 5.90E-15 |
| PHT15_I | 0.4331 | 0.1413 | 0.0936 | 0.1333 | 2.00E-16 |
| GFP14_I | 0.7636 | 0.1323 | 0.1137 | 0.1226 | 2.20E-16 |
| GFP15_I | 0.5006 | 0.1548 | 0.1966 | 0.1555 | 1.30E-12 |
| HW14_I | 0.5936 | 0.1913 | 0.2285 | 0.1896 | 4.03E-06 |
| HW15_I | 0.4475 | 0.1781 | 0.2728 | 0.1782 | 3.60E-12 |

† Regions comprised of the 20% most open segments in the genome
‡ Regions comprised of the 20% most closed segments in the genome
§ Pairwise Wilcoxon test p-value comparing open and closed regions

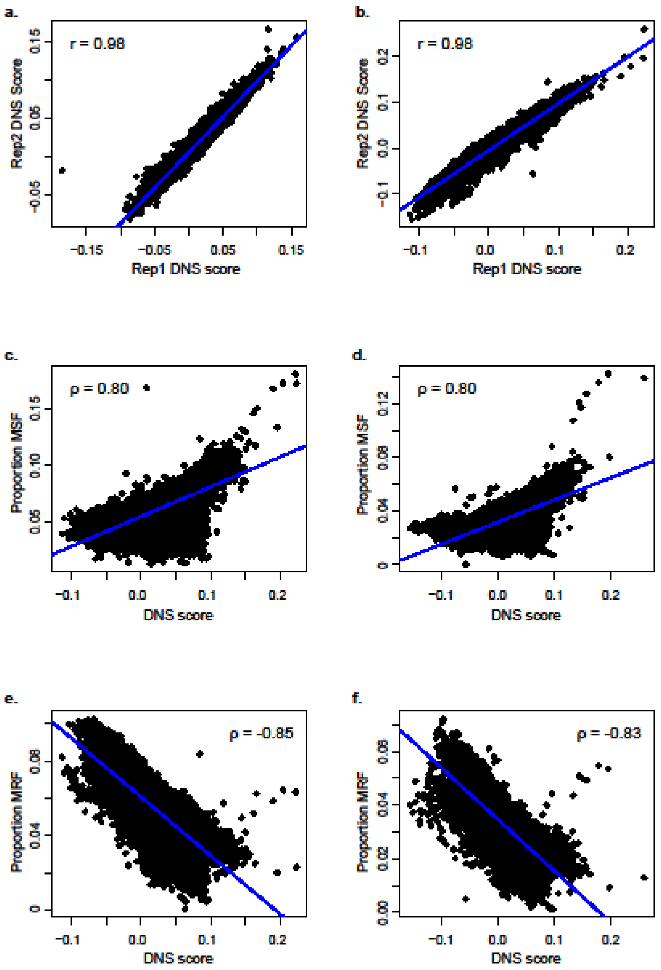


**Figure S1**. Correlation between replicates of DNS scores and iSeg results. **(a)** DNS scores of rep1 and rep2 using 2Mb windows across the genome, regression line is shown in blue, and Pearson correlation coefficient is displayed on the graph. **(b)** DNS scores of rep1 and rep2 using 1Mb windows across the genome, regression line is shown in blue, and Pearson correlation coefficient is displayed on the graph. **(c)** iSeg results as proportion of 1Mb window that is an MSF outlier with 1Mb window DNS score for replicate 1, regression line is shown in blue, and Spearman's rho is displayed on the graph. **(d)** iSeg results as proportion of 1Mb window that is an MSF outlier with 1Mb window DNS score for replicate 2, regression line is shown in blue, and Spearman's rho is displayed on the graph. **(e)** iSeg results as proportion of 1Mb window that is an MRF outlier with 1Mb window DNS score for replicate 1, regression line is shown in blue, and Spearman's rho is displayed on the graph. **(f)** iSeg results as proportion of 1Mb window that is an MSF outlier with 1Mb window DNS score for replicate 2, regression line is shown in blue, and Spearman's rho is displayed on the graph.

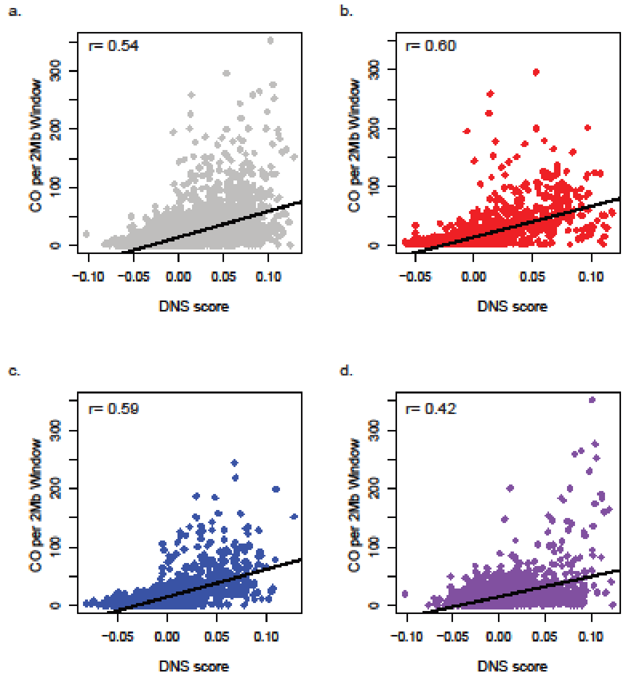


**Figure S2**. Recombination rate positively correlates with DNS score. **(a)** All genomes together, **(b)** A genome, **(c)** B genome, **(d)** D genome. For each comparison, recombination events were summed for all individuals in the spring wheat NAM population (Jordan, et al, 2018) for 2Mb windows across the genome, and correlated with 2Mb DNS score. Linear regression lines are displayed for each correlation and Pearson correlation coefficient is displayed on the figure.

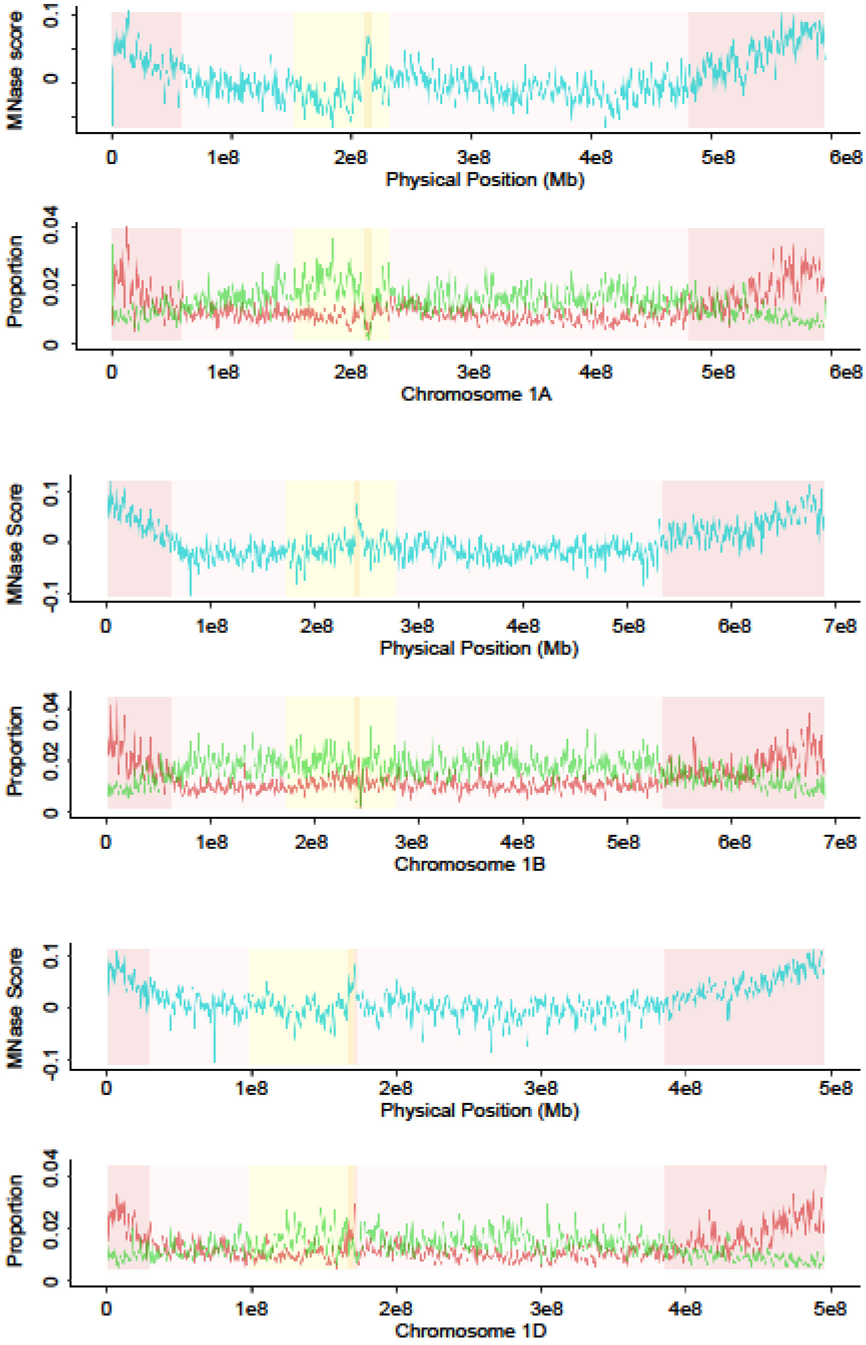


**Figure S3**. Distribution of chromatin accessibility across chromosomes 1A, 1B and 1D. For each chromosome, **top panel** shows distribution of DNS scores calculated for 1Mb windows, and **bottom panels** show proportion of MSF (red) and MRF (green) within 1Mb windows across chromosome. Genomic segments are shown in the background as dark pink for distal segments, light pink for interstitial segments, and pale yellow for the proximal region, location of the centromere is dark yellow.

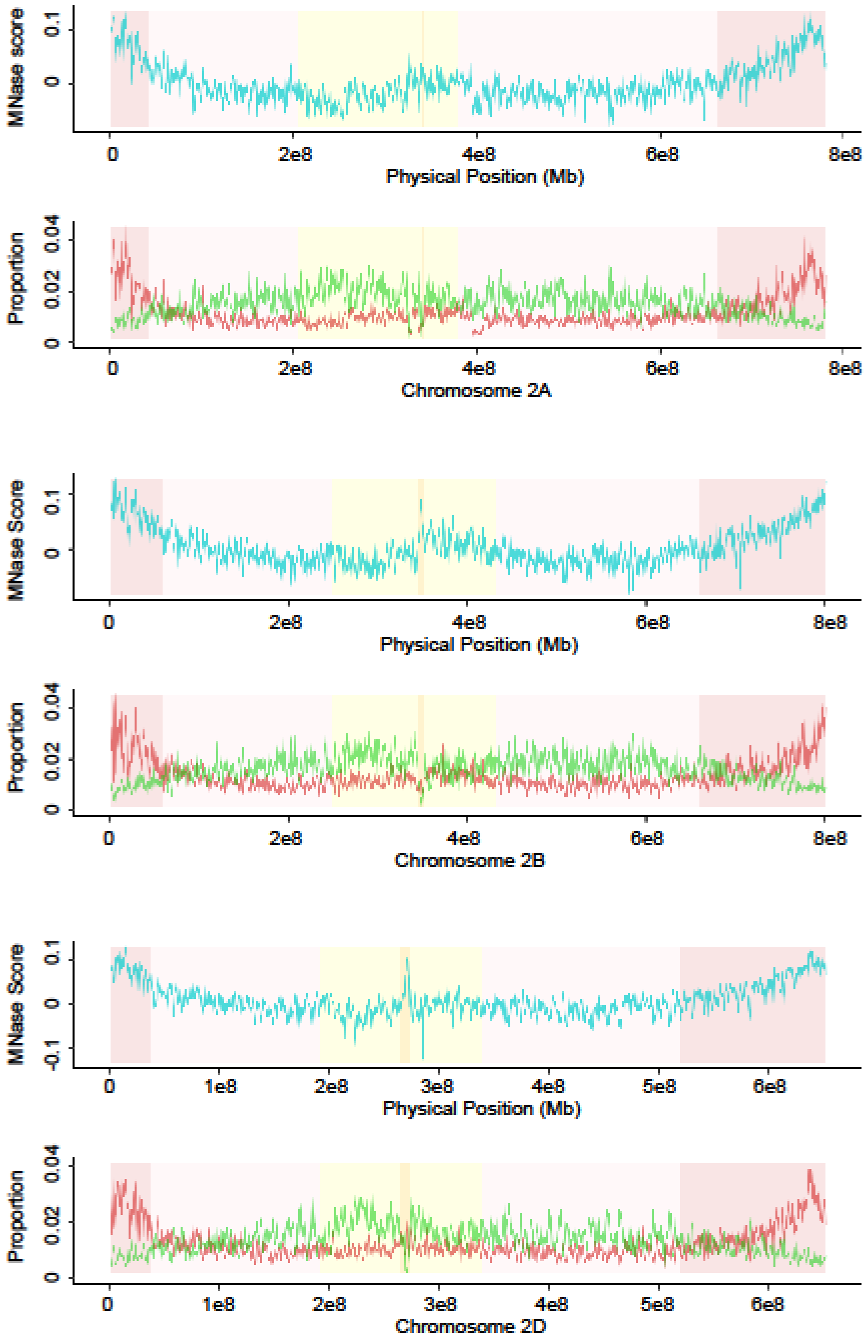


**Figure S4**. Distribution of chromatin accessibility across chromosomes 2A, 2B and 2D. For each chromosome, **top panel** shows distribution of DNS scores calculated for 1Mb windows, and **bottom panels** show proportion of MSF (red) and MRF (green) within 1Mb windows across chromosome. Genomic segments are shown in the background as dark pink for distal segments, light pink for interstitial segments, and pale yellow for the proximal region, location of the centromere is dark yellow.

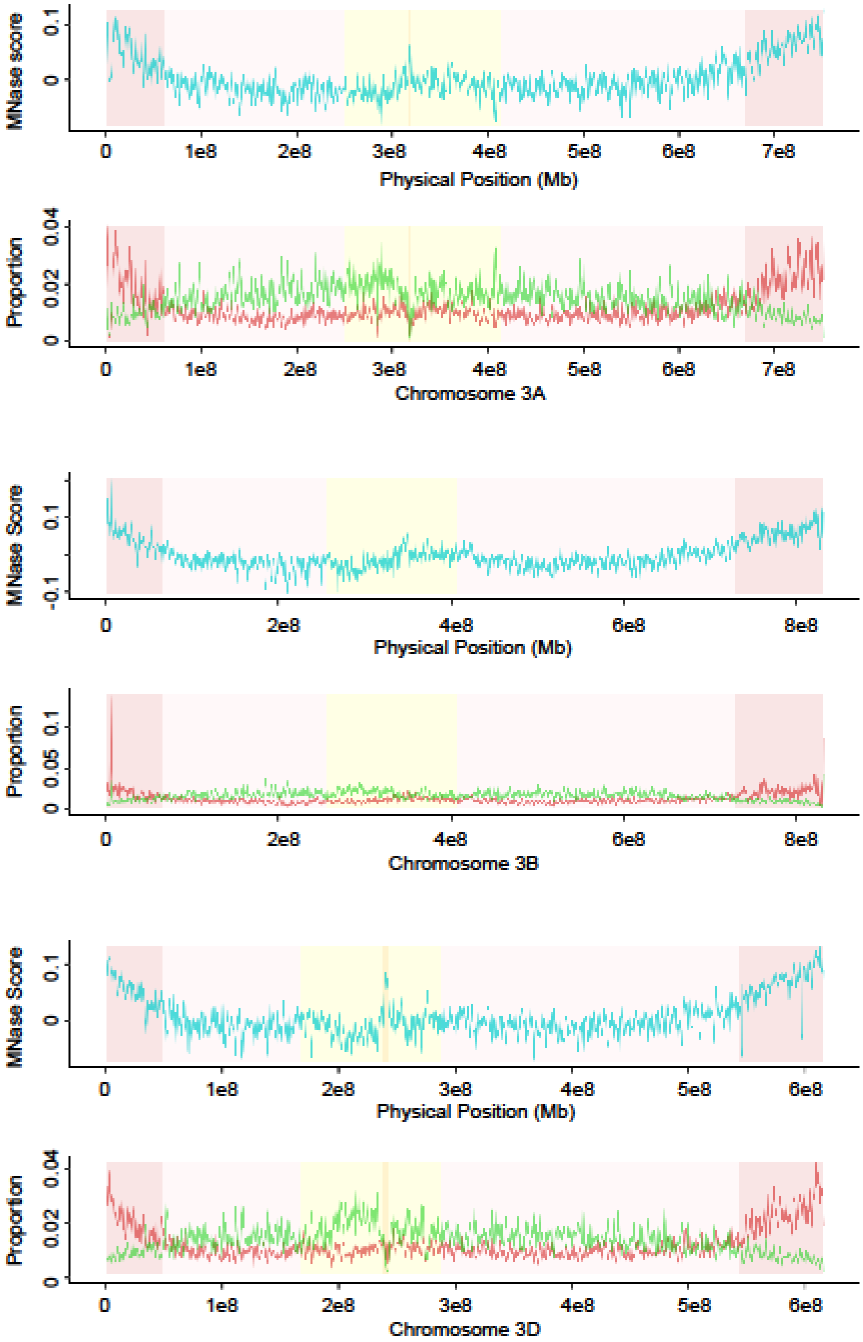


**Figure S5**. Distribution of chromatin accessibility across chromosomes 3A, 3B and 3D. For each chromosome, **top panel** shows distribution of DNS scores calculated for 1Mb windows, and **bottom panels** show proportion of MSF (red) and MRF (green) within 1Mb windows across chromosome. Genomic segments are shown in the background as dark pink for distal segments, light pink for interstitial segments, and pale yellow for the proximal region, location of the centromere is dark yellow.

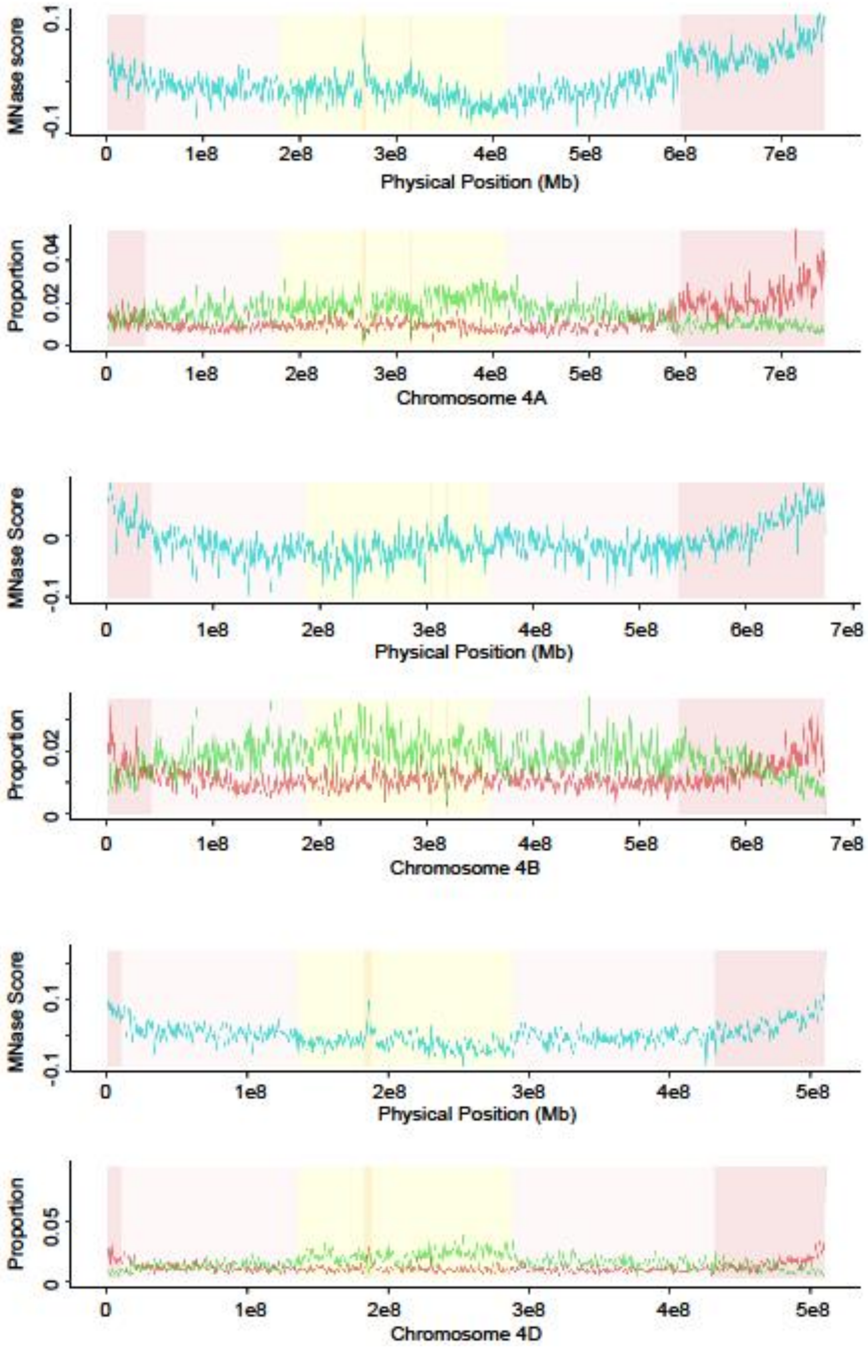


**Figure S6**. Distribution of chromatin accessibility across chromosomes 4A, 4B and 4D. For each chromosome, **top panel** shows distribution of DNS scores calculated for 1Mb windows, and **bottom panels** show proportion of MSF (red) and MRF (green) within 1Mb windows across chromosome. Genomic segments are shown in the background as dark pink for distal segments, light pink for interstitial segments, and pale yellow for the proximal region, location of the centromere is dark yellow.

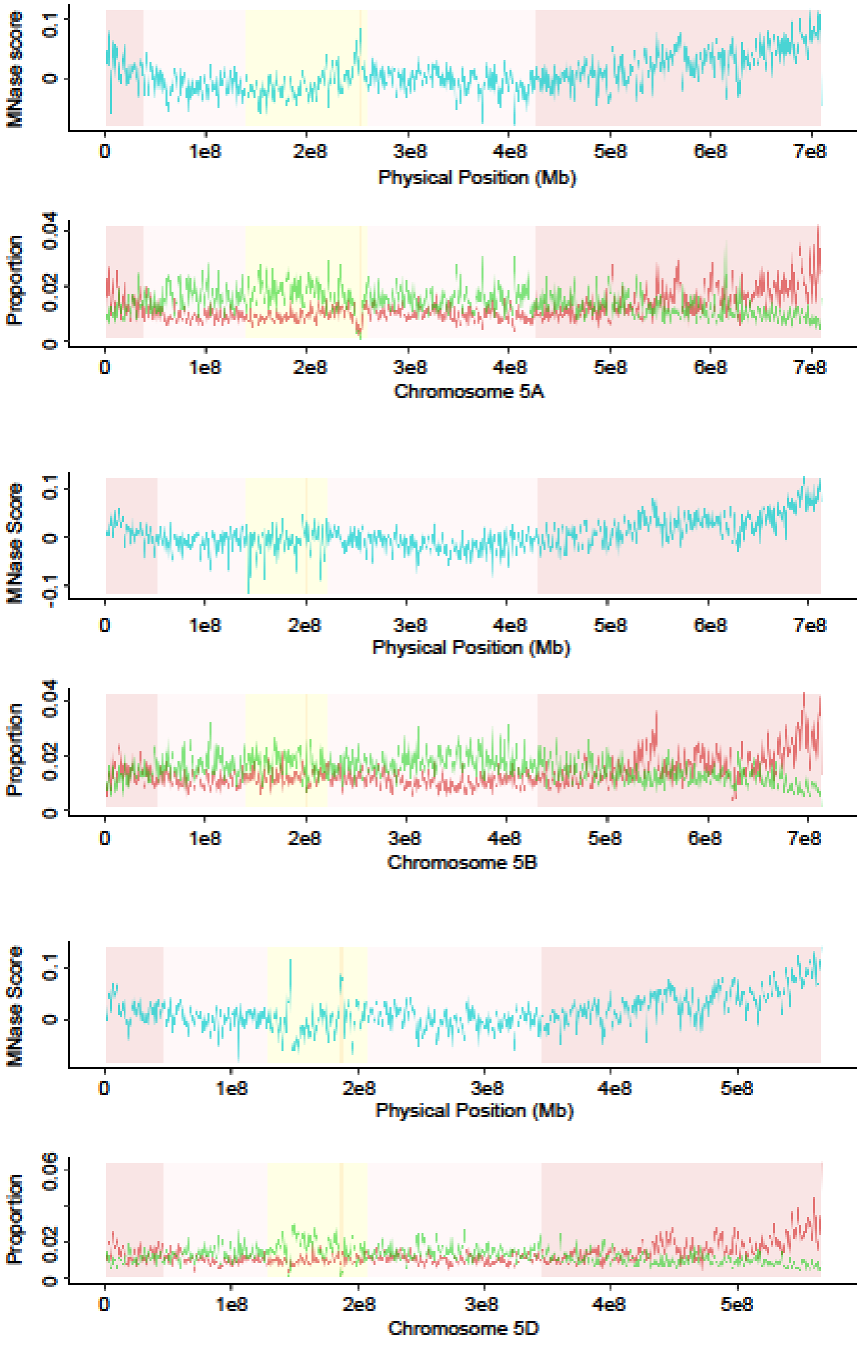


**Figure S7**. Distribution of chromatin accessibility across chromosomes 5A, 5B and 5D. For each chromosome, **top panel** shows distribution of DNS scores calculated for 1Mb windows, and **bottom panels** show proportion of MSF (red) and MRF (green) within 1Mb windows across chromosome. Genomic segments are shown in the background as dark pink for distal segments, light pink for interstitial segments, and pale yellow for the proximal region, location of the centromere is dark yellow.

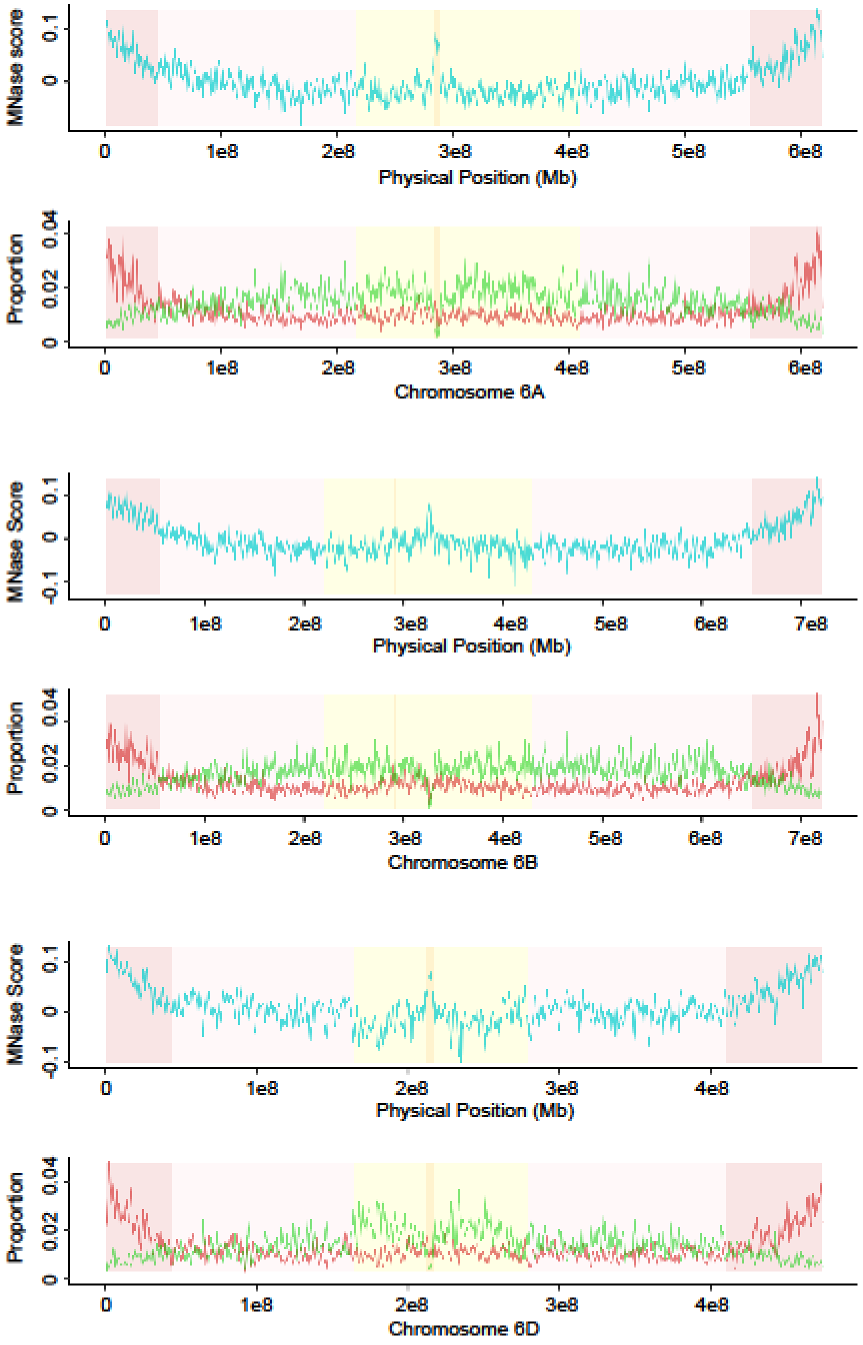


**Figure S8**. Distribution of chromatin accessibility across chromosomes 6A, 6B and 6D. For each chromosome, **top panel** shows distribution of DNS scores calculated for 1Mb windows, and **bottom panels** show proportion of MSF (red) and MRF (green) within 1Mb windows across chromosome. Genomic segments are shown in the background as dark pink for distal segments, light pink for interstitial segments, and pale yellow for the proximal region, location of the centromere is dark yellow.

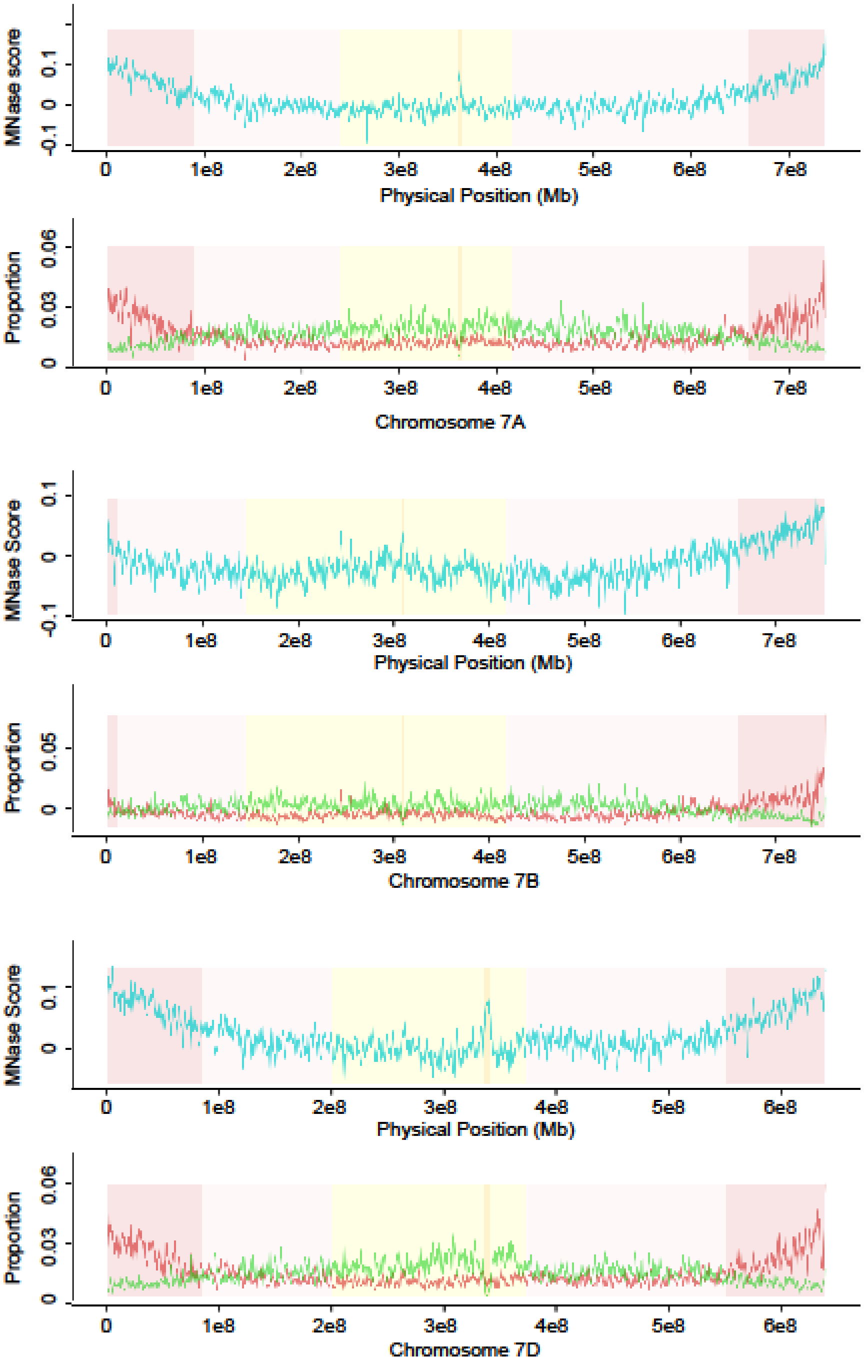


**Figure S9**. Distribution of chromatin accessibility across chromosomes 7A, 7B and 7D. For each chromosome, **top panel** shows distribution of DNS scores calculated for 1Mb windows, and **bottom panels** show proportion of MSF (red) and MRF (green) within 1Mb windows across chromosome. Genomic segments are shown in the background as dark pink for distal segments, light pink for interstitial segments, and pale yellow for the proximal region, location of the centromere is dark yellow.

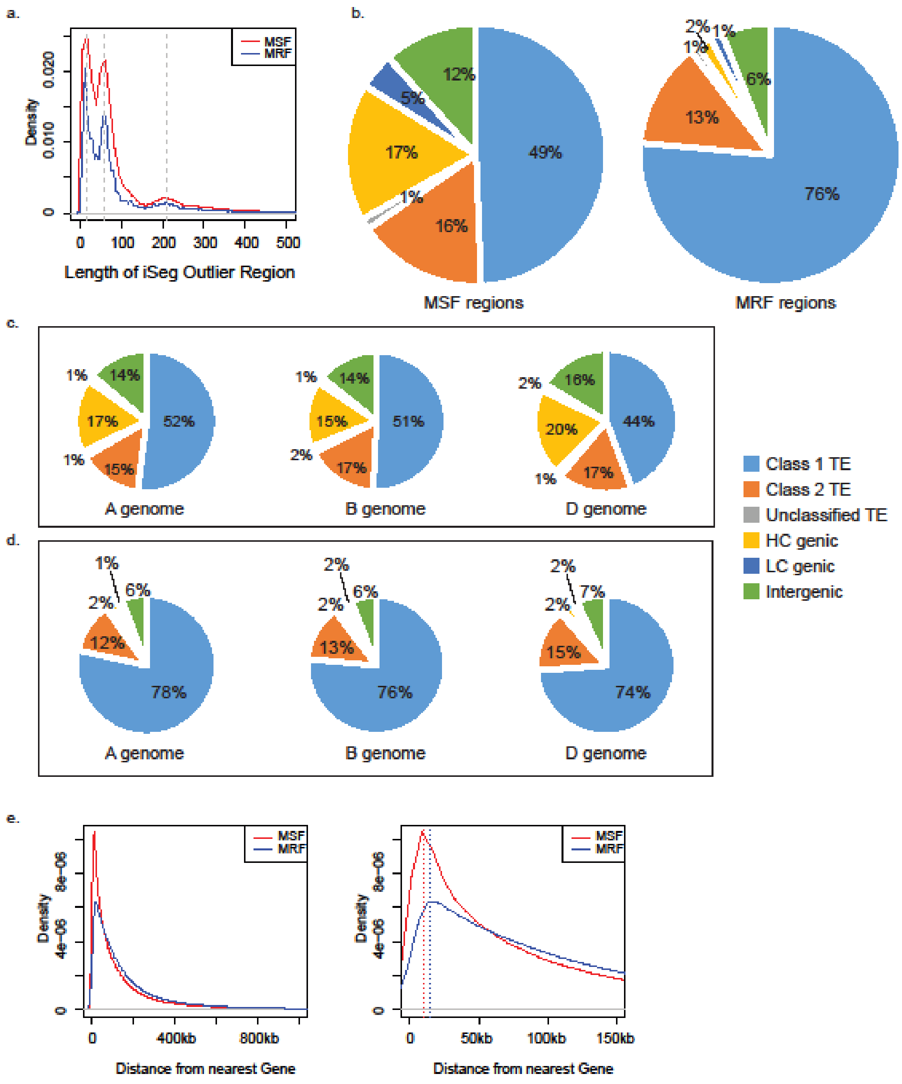


**Figure S10**. MSF and MRF outlier region descriptions. **(a)** Distribution of lengths of iSeg outlier regions for MSF (red) and MRF (blue) across the genome. Gray dashed lines represent the three peaks at 20bp, 60bp, and 210bp. **(b)** The majority of DNS outlier regions fall in Class 1 TEs (light blue), Class 2 TE regions (orange), or Unclassified TE regions (gray). MSF outliers have a significantly higher proportion of regions falling within 2 kb of annotated HC genes (yellow), and LC genes (dark blue), and the remaining MSF and MRF regions are intergenic (more than 2 kb from genes) and do not fall in annotated TE (green). **(c)** Breakdown of MSF regions by genome, and **(d)** MRF regions by genome. **(e)** For all intergenic MSF (red) and MRF (blue) regions, the distance from the nearest gene in the genome; the second panel is a narrowed down region of 150 kb, with density peaks marked by dashed line for both MSF (red) at 10 kb and MRF (blue) at 15 kb.

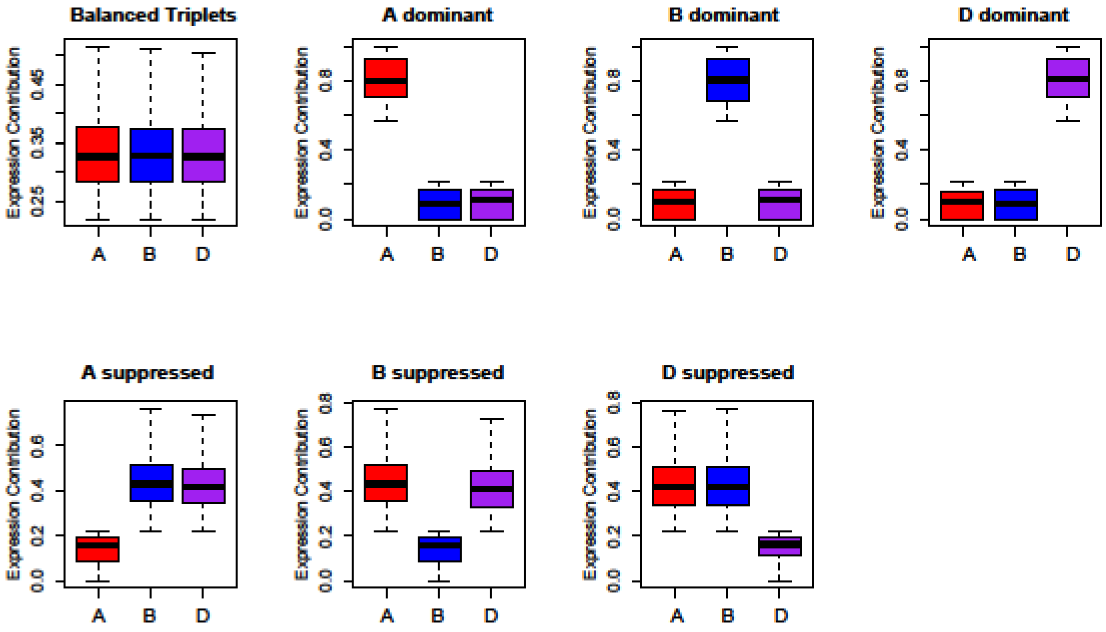


**Figure S11.** Categorized syntenic triplet expression contribution of 1:1:1 duplicated homeologous genes. Total expression for each triplet set is the sum of expression for (homeologous gene A + homeologous gene B + homoelogous gene D). Expression contribution is the proportion of expression that each genome's homeologous gene contributes to the overall expression of the triplets, *ie*. (A homelog expression / sum of triplet expression). Data shown for 12,601 syntenic expressed triplet sets, whose total expression was ≥ 0.5 TPM (Table S6). Categorization followed the method in Ramirez-Gonzalez, *et. al*. 2018. Expression was downloaded as a subset of Ramirez-Gonzalez, *et. al*. 2018 to match age and developmental state of DNS data. Distribution of expression contribution for each genome's homeologous gene for the A (red), B(blue), and D (purple) genomes, with respect to whole triplet's expression for each category.

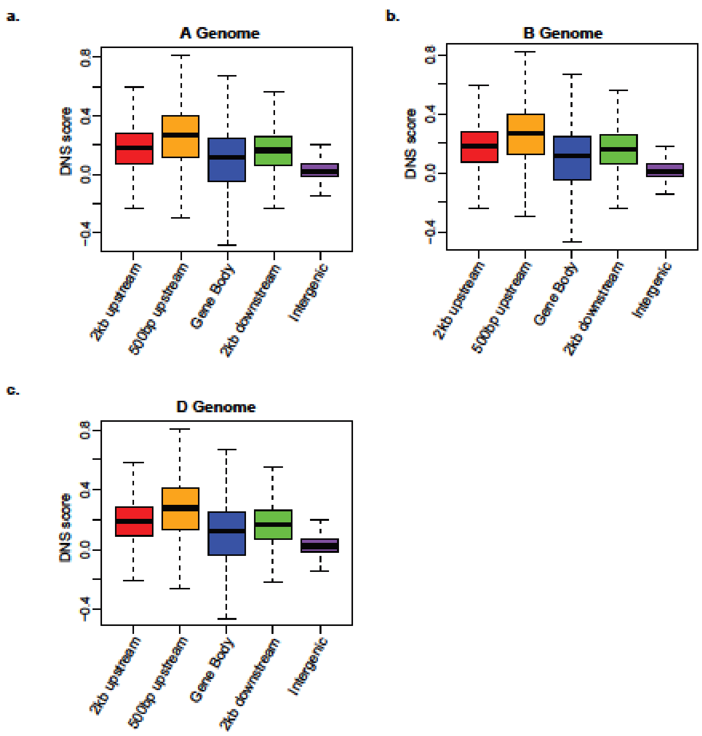


**Figure S12**. DNS scores around genes by genome. The distribution for the average DNS score for regions of 2kb upstream of CDS (red), 500 bp upstream of CDS (orange), gene body (blue), 2kb downstream of CDS (green), and intergenic regions (purple) for each of the HC genes in the **(a)** A genome, **(b)** B genome and **(c)** D genome.

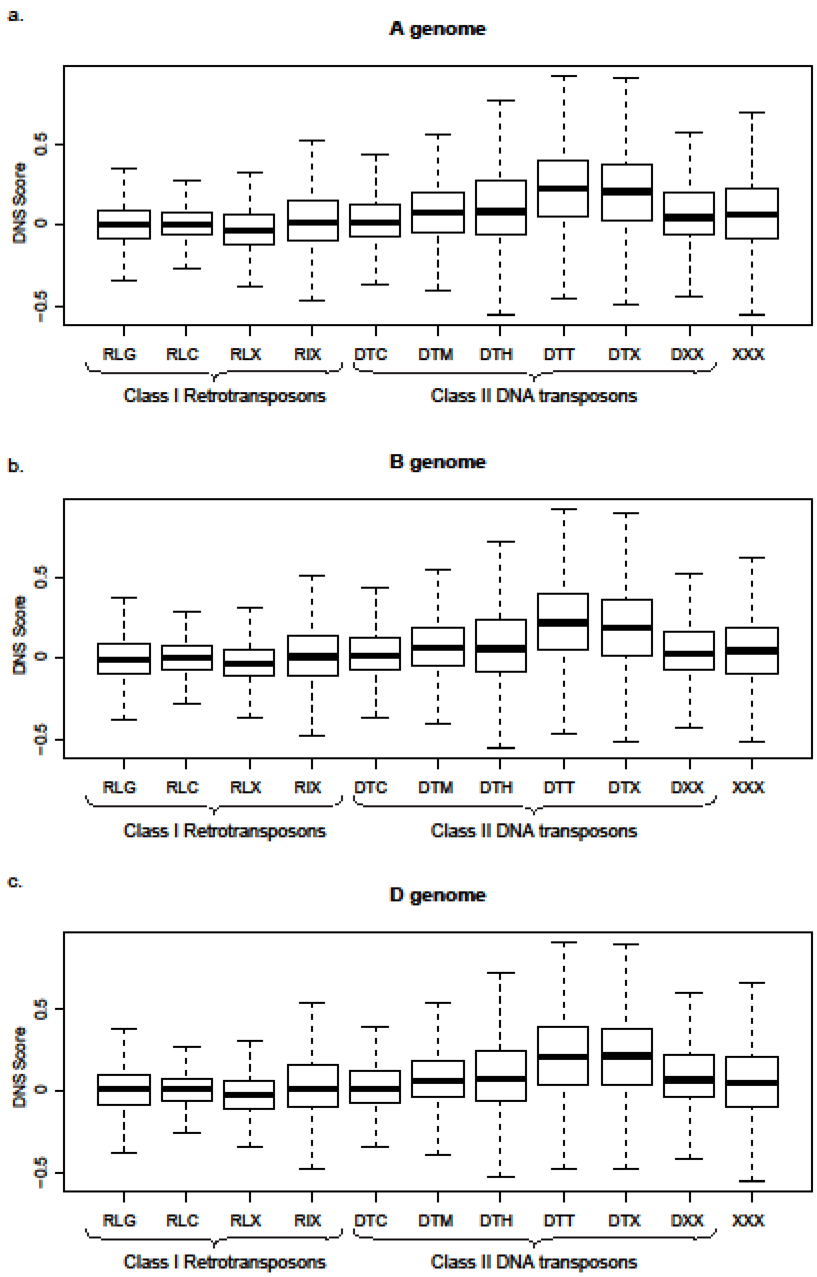


**Figure S13**. DNS score distribution for common TE superfamilies for the **(a)** A genome, **(b)** B genome, **(c)** D genome.

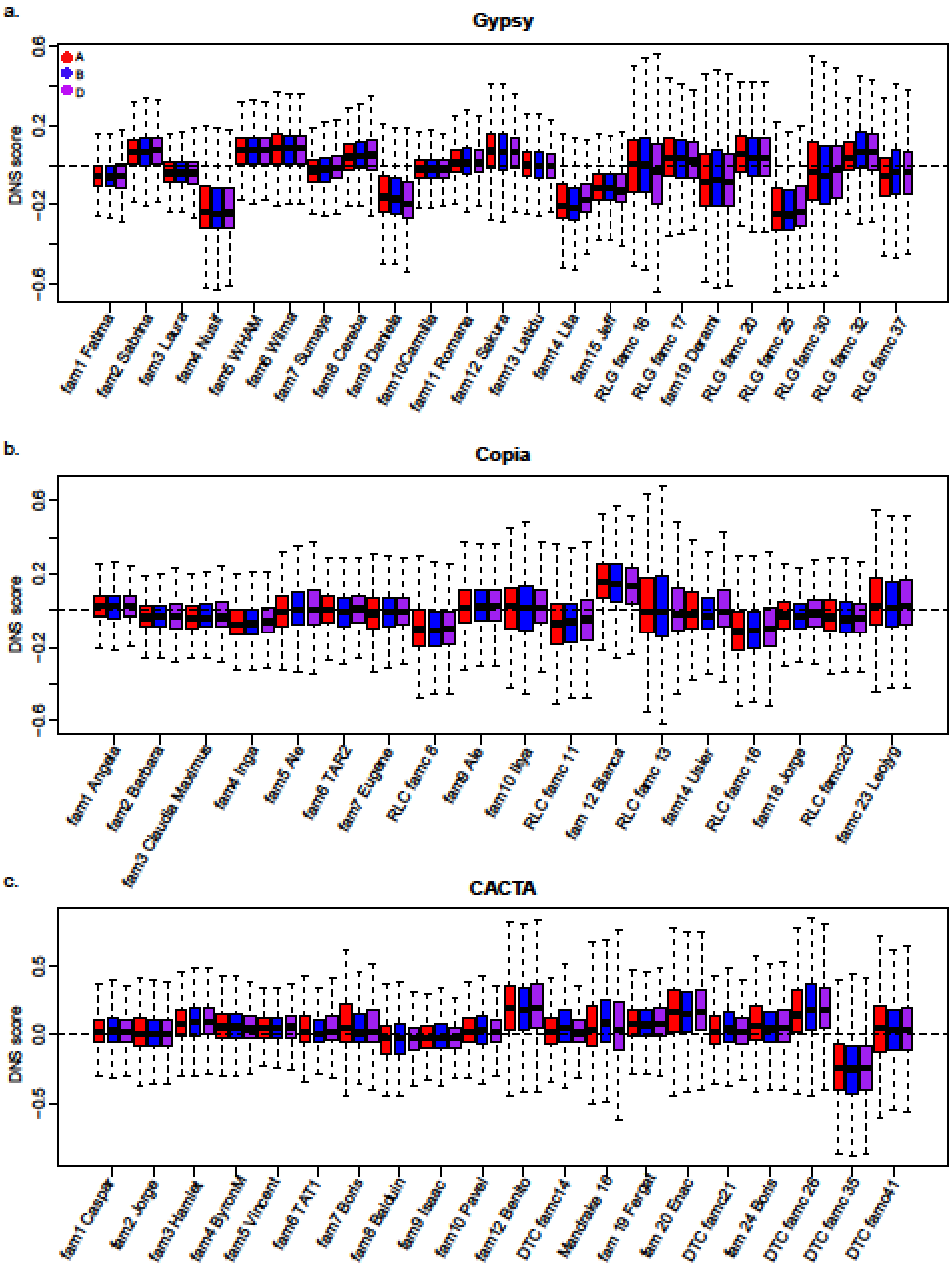


**Figure S14**. Distribution of DNS scores families within TE superfamilies **(a)** Gypsy, **(b)** Copia, **(c)** CACTA for the A (red), B (blue) and D (purple) genomes.

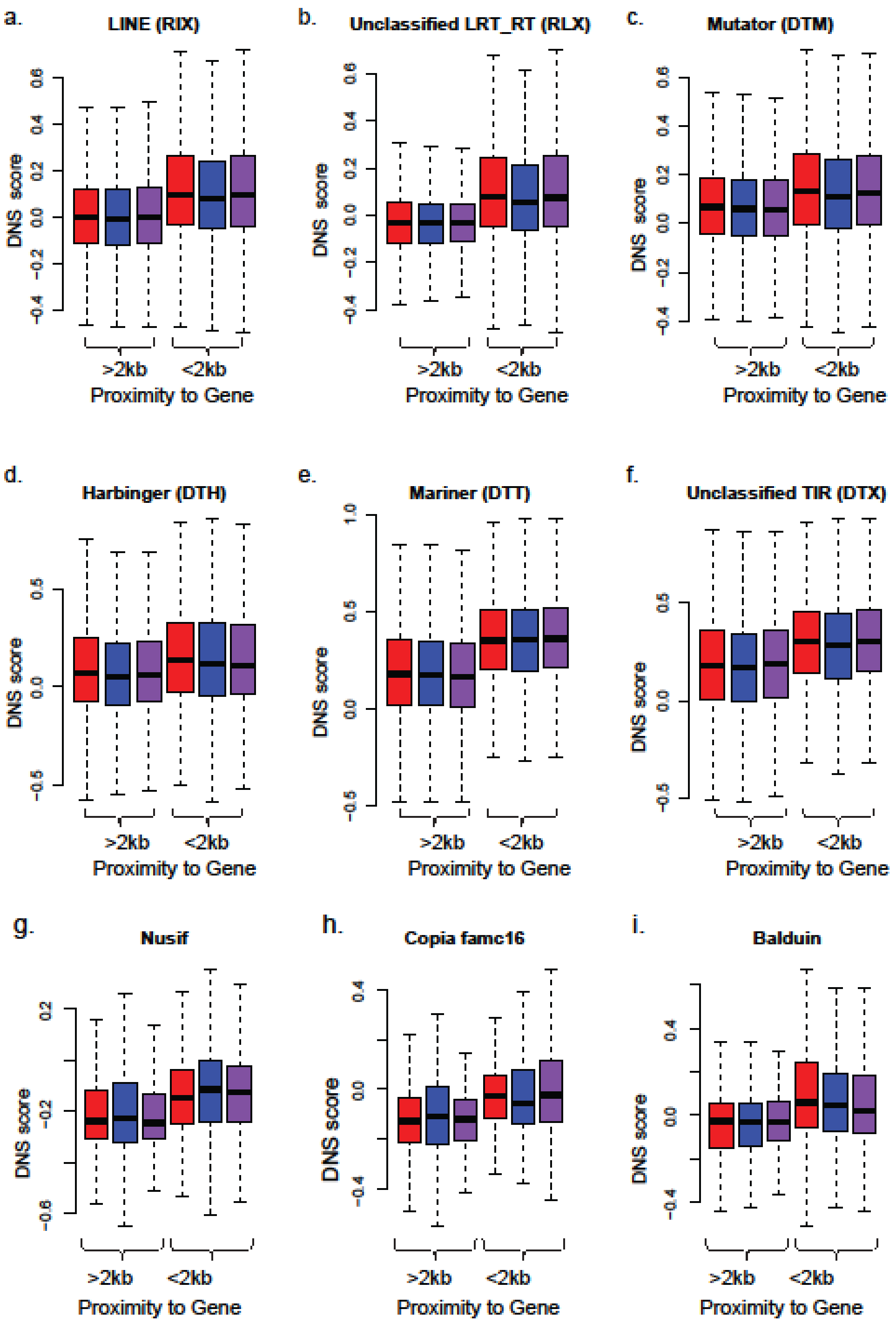


**Figure S15**. DNS scores of TE superfamilies and families located within and outside of 2 kb promoter regions of the genes. **(a)** Line retrotransposons, ($W_{RIX}$ = 800800000, *p-value* < 2.2e-16) **(b)** Unclassified LTRs, ($W_{RLX}$ = 627110000, *p-value* < 2.2e-16), **(c)** Mutator, ($W_{DTM}$ = 573740000, *p-value* < 2.2e-16), **(d)** Harbinger, ($W_{DTH}$ = 195940000, *p-value* < 2.2e-16) , **(e)** Mariner, ($W_{DTT}$ = 2288100000, *p-value* < 2.2e-16), **(f)** Unclassified DNA transposons, ($W_{DTX}$ = 935490000, *p-value* < 2.2e-16). **(g)** Gypsy family Nusif ($W_A$= 58313, $W_B$ = 68222, $W_D$ = 46973; all *p*-values < 2.2 x $10^{-16}$), **(h)** Copia family famc16 ($W_A$= 3523, *p-value* $_A$= 1.4 x $10^{-5}$; $W_B$ =

7864, *p-value* $_B$= 4 x $10^{-4}$; $W_D$ = 3190; *p*-value $_D$ < 2 x $10^{-3}$), **(i)** CACTA family Balduin ($W_A$= 11367, *p-value* $_A$= 2.2 x $10^{-9}$; $W_B$ = 28578, *p-value* $_B$= 7.5 x $10^{-9}$; $W_D$ = 39879; *p*-value $_D$ < 4 x $10^{-6}$).

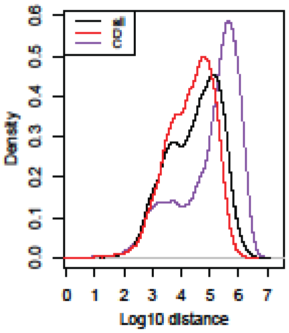


**Figure S16.** Comparison of Intergenic Distance for Distal and Centromeric Regions. Density of intergenic space between adjacent genes based on all segments in genome (black), distal regions only (red), and centromeric regions only (purple). The intergenic distance in centromeric region is much longer on average than in the distal regions, mean $_{Distal}$ = 70kb, mean $_{Cent.}$ = 418kb.

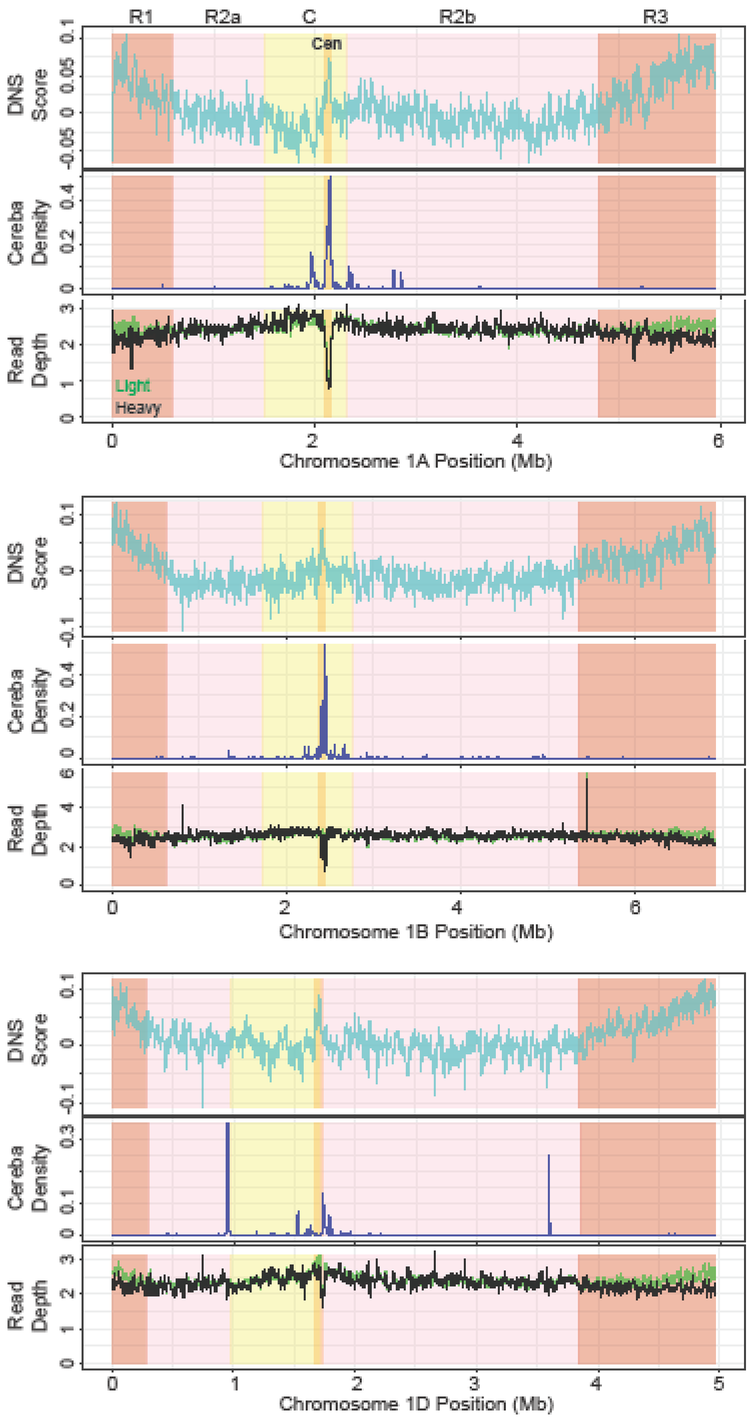


**Figure S17**. Sensitivity of centromeric chromatin to differential MNase digest. Distribution of DNS scores (top panel), Cereba LTR density (middle panel), and depth of read coverage obtained by light (green) and heavy (black) MNase digest (bottom panel) for homeologous chromosome group 1. Each chromosome is divided into five regions R1, R2a, C, R2b, and R3. The location of centromere (Cen- dark yellow) is based on previously published studies (The International Wheat Genome Sequencing Consortium (IWGSC) 2018) that used anti-CENH3 chromatin immunoprecipitation method.

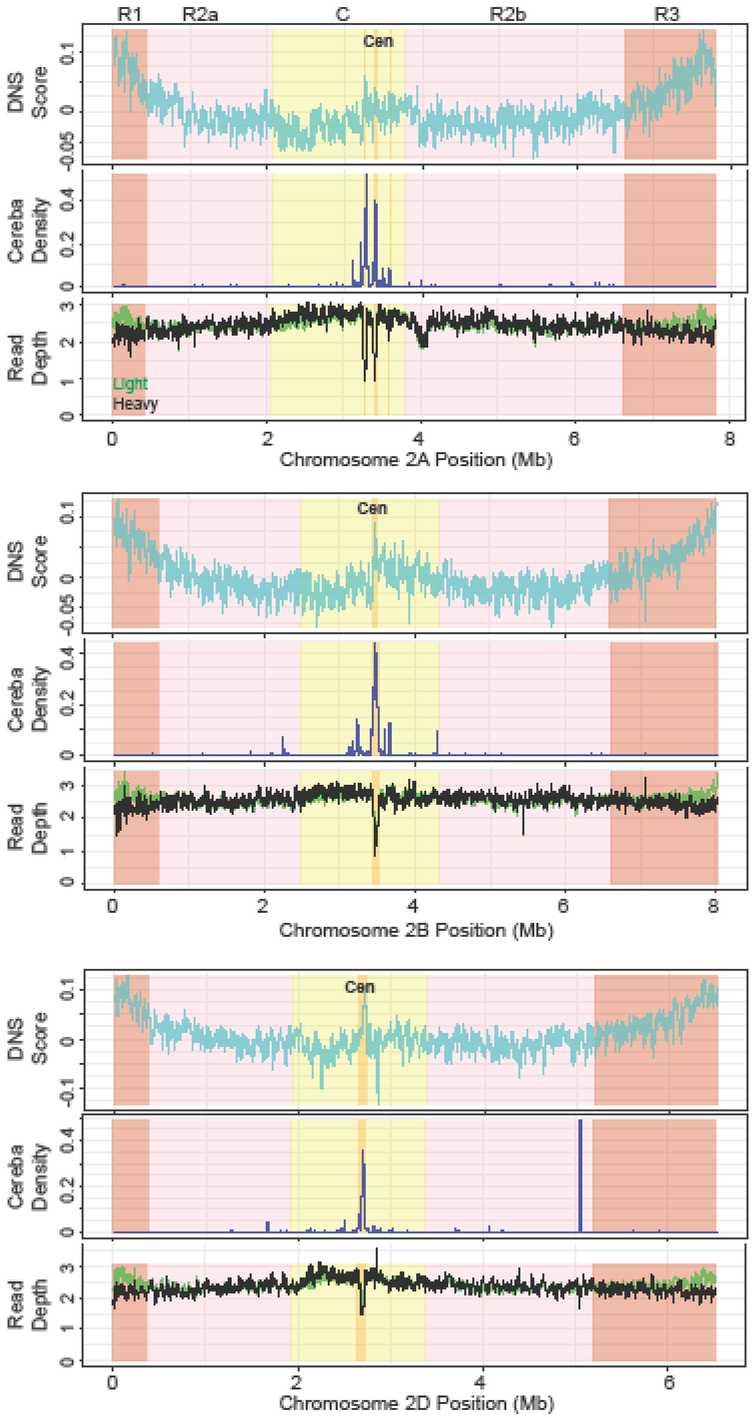


**Figure S17 continued**. Distribution of DNS scores (top panel), Cereba LTR density (middle panel), and depth of read coverage obtained by light (green) and heavy (black) MNase digest (bottom panel) for homeologous chromosome group 2.

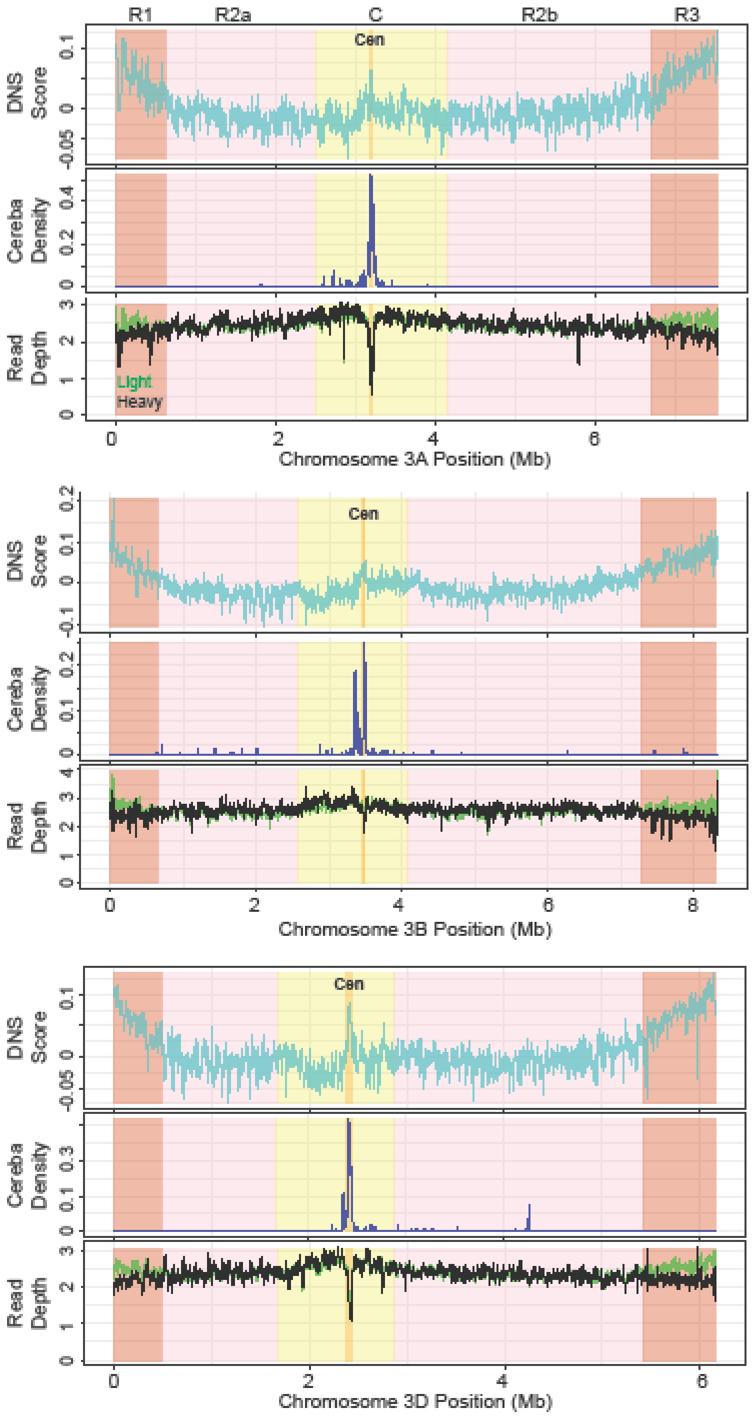


**Figure S17 continued**. Distribution of DNS scores (top panel), Cereba LTR density (middle panel), and depth of read coverage obtained by light (green) and heavy (black) MNase digest (bottom panel) for homeologous chromosome group 3.

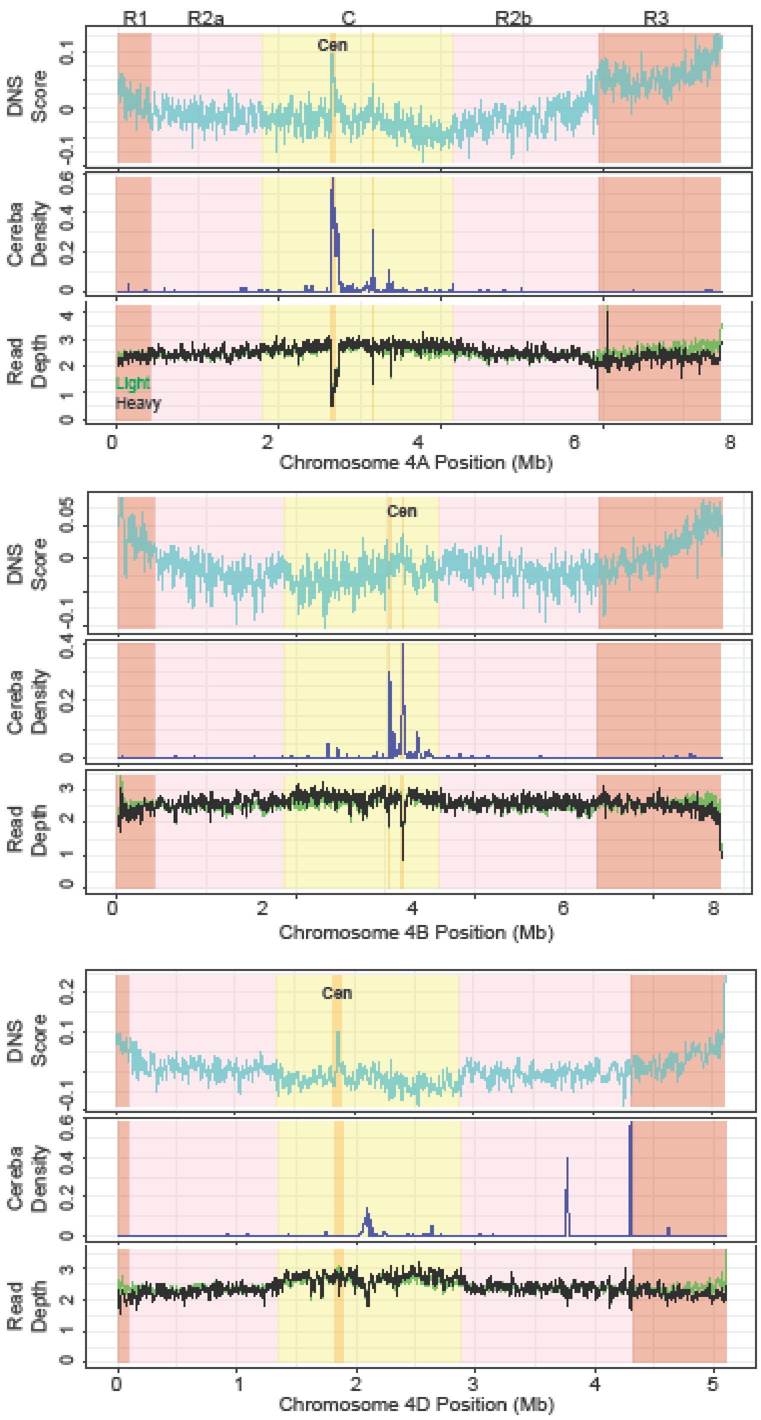


**Figure S17 continued**. Distribution of DNS scores (top panel), Cereba LTR density (middle panel), and depth of read coverage obtained by light (green) and heavy (black) MNase digest (bottom panel) for homeologous chromosome group 4.

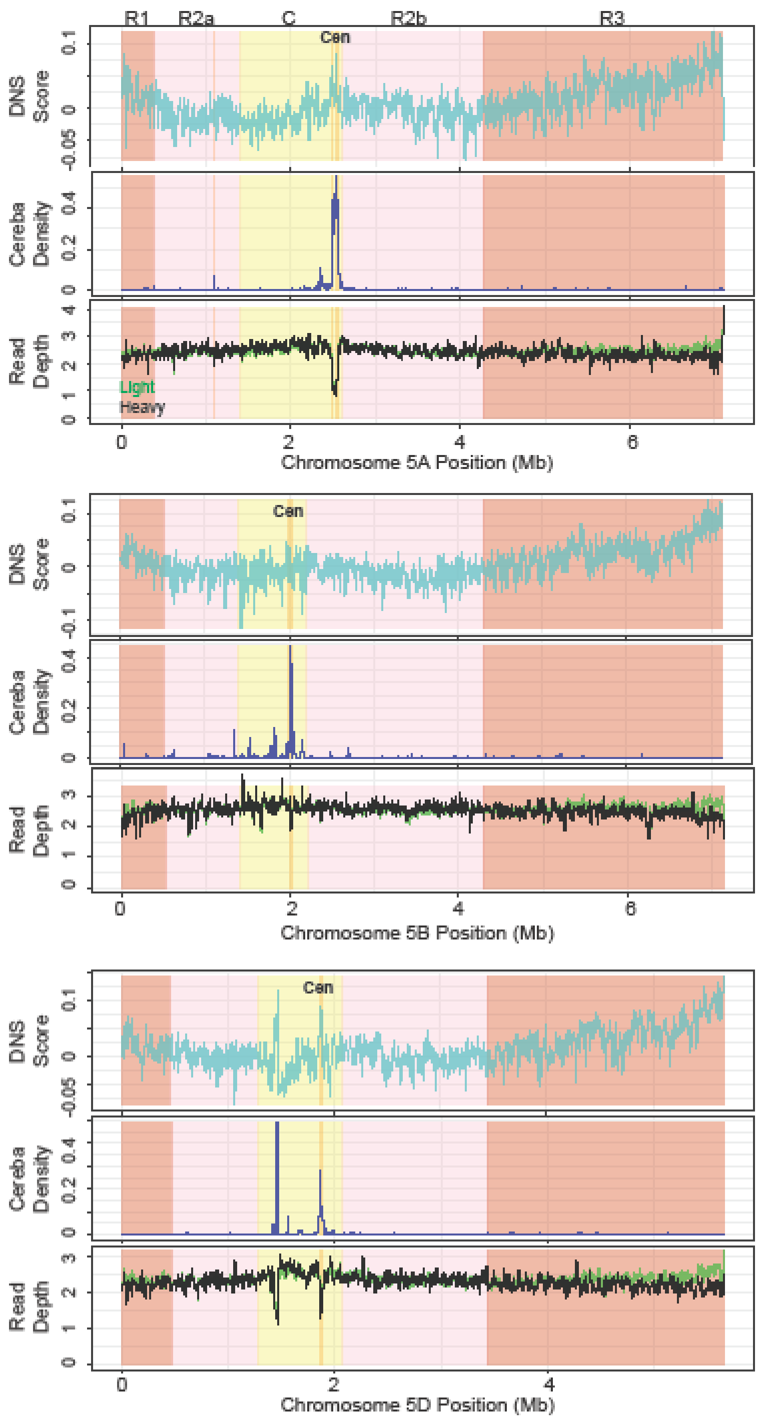


**Figure S17 continued**. Distribution of DNS scores (top panel), Cereba LTR density (middle panel), and depth of read coverage obtained by light (green) and heavy (black) MNase digest (bottom panel) for homeologous chromosome group 5.

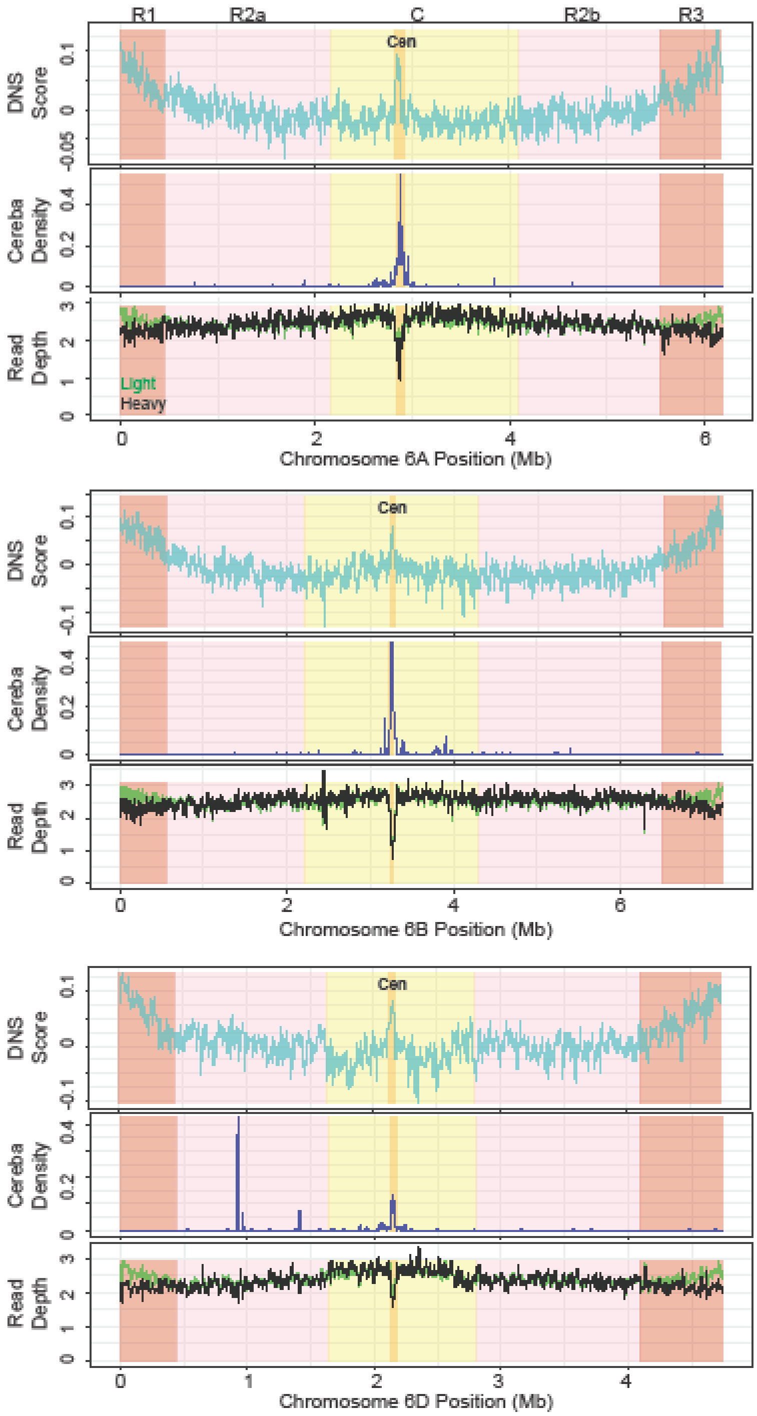


**Figure S17 continued**. Distribution of DNS scores (top panel), Cereba LTR density (middle panel), and depth of read coverage obtained by light (green) and heavy (black) MNase digest (bottom panel) for homeologous chromosome group 6.

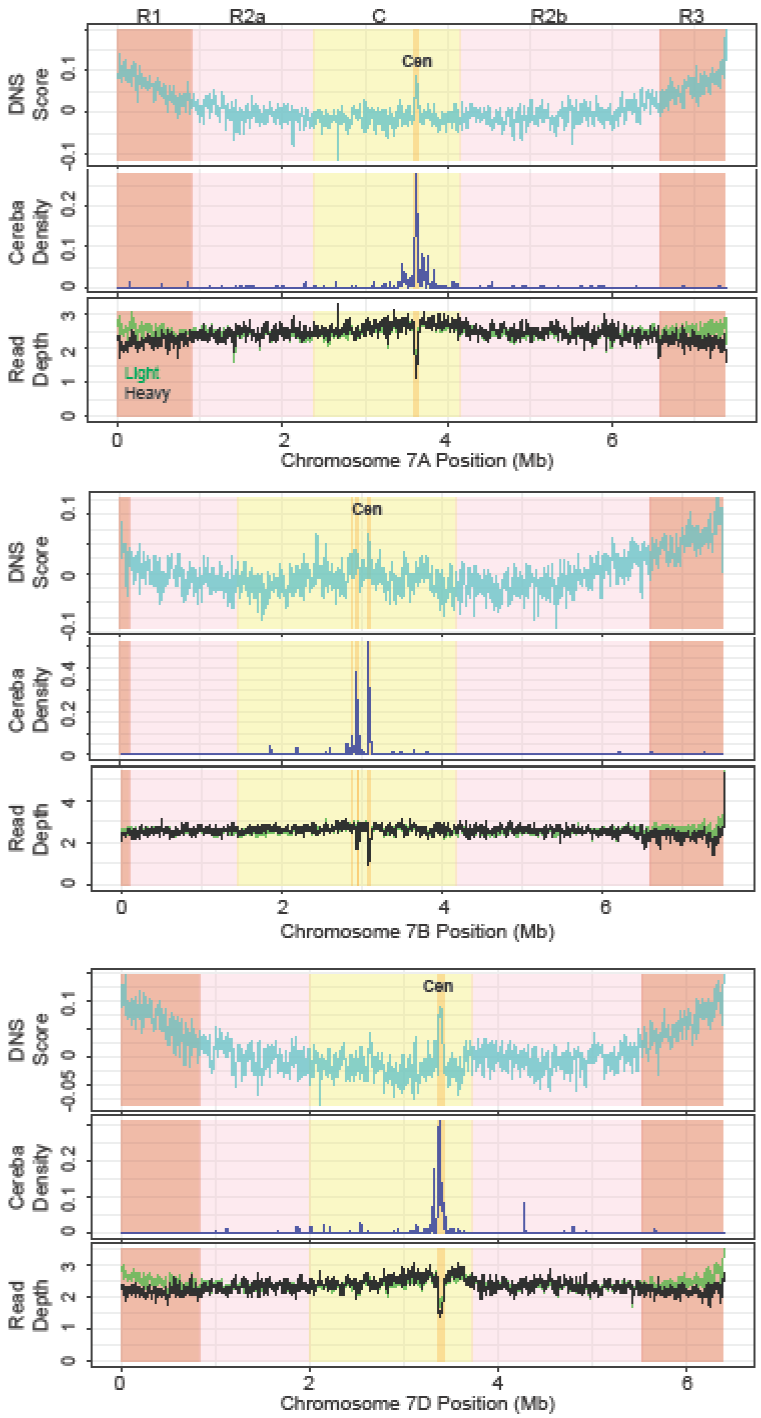


**Figure S17 continued**. Distribution of DNS scores (top panel), Cereba LTR density (middle panel), and depth of read coverage obtained by light (green) and heavy (black) MNase digest (bottom panel) for homeologous chromosome group 7.
